# Artemisinin-based hybrids produce intracellular proteasome inhibitors that overcome resistance in *Plasmodium falciparum*

**DOI:** 10.1101/2021.06.21.449268

**Authors:** Wenhu Zhan, Yi Jing Liu, Changmei Yang, Hao Zhang, Jacob C. Harris, Rong Wang, Songbiao Zhu, Julian Sherman, George Sukenick, Ana Rodriguez, Haiteng Deng, Carl F. Nathan, Laura A. Kirkman, Gang Lin

**Affiliations:** Department of Microbiology & Immunology, Weill Cornell Medicine, 1300 York Avenue, New York, NY 10065; Department of Medicine, Division of Infectious Diseases, 1300 York Avenue, New York, NY 10065; MOE Key Laboratory of Bioinformatics, Center for Synthetic and Systematic Biology, School of Life Sciences, Tsinghua University, Beijing 100084, China; NMR Analytical Core Facility, Memorial Sloan Kettering Cancer Center, New York, NY 10065; Division of Parasitology, Department of Microbiology, New York University School of Medicine, New York, NY, USA

**Keywords:** Malaria, artemisinin resistance, ART-*Pf*20S inhibitor hybrids, resistance circumvention.

## Abstract

Artemisinin resistant *Plasmodium falciparum* (*Pf*) is spreading despite combination chemotherapy (ACT). Here we report the design of artezomibs, single-molecule hybrids of an artemisinin and a *Pf*-selective proteasome inhibitor. Artezomibs exert a novel mode of action inside the malaria parasites. The artemisinin component covalently modifies parasite proteins, which become substrates of the *Pf* proteasome. The proteasomal degradation products that bear the proteasome inhibitor component of the hybrid then inhibit *Pf* proteasomes, including those with mutations that reduce binding affinity of the proteasome inhibitor component on its own. We demonstrated that artezomibs circumvent both artemisinin resistance conferred by Kelch13 polymorphism and resistance to the proteasome inhibitor associated with mutations in *Pf* proteasomes. This mode of action may enable the use of a single molecule with one pharmacokinetic profile to prevent the emergence of resistance.

## Main

Artemisinin (ART) is currently the backbone of the treatment for malaria, a protozoal infection responsible for 200 million cases and almost half a million deaths each year ^1^. ART is a pro-drug that it is activated by hemoglobin-derived heme within the parasites. Activation converts ART to radicals that cause extensive oxidative damage to lipids and proteins. Oxidized proteins overload the parasites’ ubiquitin-proteasome degradation system (UPS), leading to parasite death ^2–6^. With increasing cases of malaria recrudescence following ART monotherapy, the use of ART-based combination therapy (ACT) was implemented. However, ACT treatment failure is now widespread across Southeast Asia and has been associated with mutations in a protein called Kelch13 (K13). Importantly, K13 mutations are now appearing in Africa and South America,^7–10^ portending the potential spread of ART resistance in the regions of the world where malaria is most widespread. Novel approaches are therefore needed to prevent a potential public health crisis in regions affected by ART resistance. The hallmark of ART resistance is increased tolerance to ART at the early ring stage of the parasites’ erythrocytic cycle. Multiple mechanisms of resistance have been associated with K13 polymorphisms ^11,12^, including reduced ART activation arising from defects in hemoglobin catabolism that reduce the abundance of free heme ^13^, reduction in proteotoxic stress ^14^, and prolongation of the ring stage of intra-erythrocytic *Pf* development ^15^. ACT is additionally limited by the divergent pharmacokinetic profiles of the individual drugs; intermittent de-facto monotherapy forfeits their intended synergistic effect against the emergence of ART resistance ^16^.

Treatment of *Pf* with ART leads to accumulation of polyubiquitinated proteins ^2^. Polyubiquitination targets proteins for degradation in the proteasome (*Pf*20S). Killing of the malaria parasite by ART occurs when protein damage exceeds the capacity of the protein degradation machinery. *Pf* parasites at erythrocytic, liver, gametocyte and gamete activation stages are highly susceptible to proteasome inhibition, suggesting essential functions of *Pf*20S in all lifecycle stages and thus making the *Pf* proteasome an appealing target for antimalarials development ^17–25^..

We and others have demonstrated that various classes of proteasome inhibitors with selectivity against the malarial proteasome over human proteasomes showed synergistic anti- malarial effects with ART ^18,25^. The synergy may arise in part because treatment of *Pf* with ART leads to accumulation of misfolded proteins with toxic effects and proteasome inhibition prevents the breakdown and removal of damaged proteins. As with other antimalarials, *Pf* parasites can develop resistance to proteasome inhibitors, albeit this was demonstrated *in vitro* and required a high parasite inoculum compared to other compounds in development, suggesting a high in vitro barrier to resistance and in some cases, only a minor shift in EC_50_ could be achieved ^17,18,25^. Given that proteasome inhibitors not only kill *Pf* on their own but also make the parasites more susceptible to ART, we hypothesized that linking a proteasome inhibitor to an ART analog through a tether could yield a hybrid compound with the ability to hijack the parasite ubiquitin proteasome system to produce a host of proteasome inhibitors with the ability to overcome resistance to each of the hybrid’s two constituent chemophores. We reasoned that an ART-proteasome inhibitor hybrid would yield ART-modified proteins whose proteasomal degradation products bearing a proteasome inhibitor moiety could inhibit the function of *Pf*20S by binding to its active proteolytic subunits. By binding distal to the *Pf*20S active sites, the extended peptides of the degradation products could compensate for a loss of binding affinity caused by point mutations near the active sites that would otherwise reduce the efficacy of the proteasome inhibitor. These peptides bearing a proteasome inhibitor via an ART component would only be formed in parasites treated with the hybrid compounds and not when the two chemophores are administered to humans in a 1:1 ratio. Here we report combining the ART and proteasome inhibitor moieties into one small molecule, termed an artezomib (ATZ), that overcomes resistance to its individual components and potentially prevents the emergence of resistance to each.

## Results

### Design and test of ATZs

To establish if an ART interfere with the binding of a *Pf*20S inhibitor to the binding pockets of the *Pf*20S, we conjugated a commercially available artesunate with PKS21224 and PKS21208, two asparagine ethylenediamines, a novel class of proteasome inhibitors ^25,26^, where isoxazoly was replaced with succinate (Scheme S1). The ART moiety was coupled at the P4 position, as the S4 pocket is partially exposed to solvent. We hypothesized that, at this position, the bulky artemisinin would not interfere with the binding of the rest of the molecule to the active site of the *Pf*20S β5 subunit. WZ-06 and WZ-13 were synthesized (Scheme S2) and their structures confirmed. WZ-20 was synthesized as an inactive proteasome inhibitor control. We next determined their IC_50_ values against *Pf*20S and human constitutive (c-20S) and immunoproteasomes (i-20S) (Table S1). Artesunate itself does not inhibit the β5 subunits of *Pf*20S, human i-20S or human c-20S. In contrast, conjugates WZ-06 and WZ-13 were potent against *Pf*20S β5 at 6 nM and 2 nM, respectively. The data suggested that ARTs at the P4 position do not interfere with the binding of AsnEDAs to *Pf*20S. However, the semi-ketal ester of the artesunate is not stable in human blood plasma, making it difficult to interpret the activity of ester-based ATZs against *Pf* parasites in red blood cells.

We thus designed and synthesized four new ATZs (Figure 1A and Schemes S3 – S5) with a stable amide instead of an ester tether to improve the stability. In accordance, we replaced the control compound artesunate with ART1, where the semi-ketal ester was changed to a cyclic ether with a carboxamide, and replaced the proteasome inhibitor control WZ-20 with PI01, the analog of the proteasome inhibitor moiety of ATZs. Deoxy-ATZ4 was also synthesized as a control for ATZ4 (Scheme S5). This compound cannot be activated by heme because it lacks the endoperoxide in the ART-related moiety. We determined their IC_50_ values against *Pf*20S, c-20S and i-20S (Figure 1B and Table 1). ART1 showed weak inhibitory activity against *Pf*20S at 9.23 μM. The IC_50_ of ATZ1 against *Pf*20S increased 10.5-fold to 0.063 μM compared to PI01. ATZ3, with a propionate linker between ART1 and the proteasome inhibitor, displayed 106-fold and 760-fold selectivity against *Pf*20S over i-20S and c-20S, while ATZ4 with a butyrate linker showed 45-fold and 250-fold selectivity, respectively. ATZ3 and ATZ4 showed increased selectivity in enzyme inhibition compared to ATZ1 and ATZ2. Deoxy-ATZ4 showed comparable IC50 values to those of ATZ4, indicating no significant effects from either the endoperoxide- or deoxy-ART moiety. The results also suggest that the propionate linker best balances potency and selectivity among these compounds.

**Figure 1.**
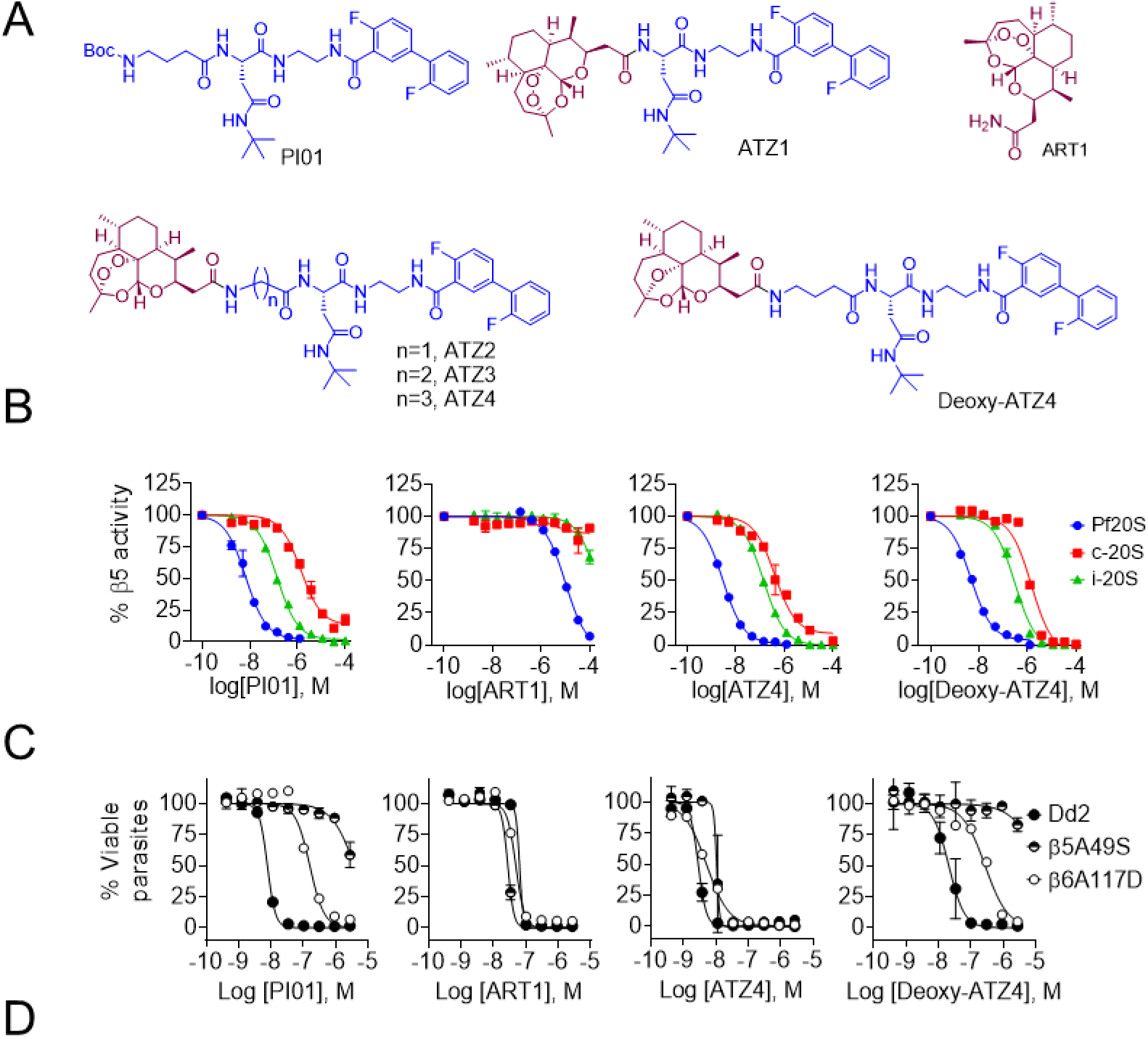
Design of hybrids of ART and proteasome inhibitors and their inhibition of proteasomes and of parasite growth. (A) Structures of proteasome inhibitor, ART analog and hybrids. (B) Inhibition of Pf20S, human c-20S and i-20S. (C) Growth inhibition of Dd2, Dd2β5A48S and Dd2β6A117D by PI01, ART1, ATZ4 and deoxy-ATZ4. (D) Table of proteasome inhibition activity and *Plasmodium* growth inhibition by the compounds.

**Table 1.** Proteasome inhibition activity and *Plasmodium* growth inhibition of hybrids.

| ID | IC<sub>50</sub> (nM) |  |  | EC<sub>50</sub> (nM) |  |  |  |
| --- | --- | --- | --- | --- | --- | --- | --- |
|  | Pf20S | Hu i-20S | Hu c-20S | Pf Dd2 | Dd2(β6A117D) | Dd2(β5A49S) | HepG2 |
| ART1 | 9,200 ± 720 | > 100,000 | > 100,000 | 43.7 ± 12.7 | 27.9 ± 11 | 30.2 ± 2.6 | > 100,000 |
| PI01 | 6.0 ± 0.8 | 200 ± 46 | 2,000 ± 240 | 7.4 ± 0.2 | 121 ± 20 | >2,770 | > 100,000 |
| ATZ1 | 63.2 ± 11 | 650 ± 94 | 3,600 ± 490 | 4.5 ± 1.6 | 4.1 ± 0.4 | 10.6 ± 4.2 | 7,300 ± 2,100 |
| ATZ2 | 2.9 ± 0.6 | 73 ± 19 | 210 ± 40 | 3.0 ± 0.6 | 4.4 ± 1.1 | 10.9 ± 2.6 | 5,800 ± 620 |
| ATZ3 | 2.4 ± 0.4 | 260 ± 51 | 18,00 ± 240 | 4.4 ± 1.5 | 12.6 ± 0.8 | 13.1 ± 2.9 | 20,300 ± 4,700 |
| ATZ4 | 4.3 ± 0.9 | 200 ± 57 | 890 ± 140 | 4.3 ± 1.3 | 6.7 ± 2.2 | 5.9 ± 1.9 | 9,400 ± 1,700 |
| Deoxy-ATZ4 | 4.4 ± 0.5 | 300 ± 22 | 1100 ± 46 | 21.6 ± 1.4 | 246 ± 44.5 | >2,770 |  |

All data are means of at least three independent experiments and are presented with standard deviation. All data were round to three or fewer significant figures.

We tested these compounds against *Pf* Dd2 and two proteasome inhibitor-resistant strains derived from Dd2, Dd2(β6A117D) and Dd2(β5A49S), which harbor a mutation in the β6 subunit (A117D) or in the β5 subunit (A49S) of *Pf*20S, respectively ^17,25^ (Figure 1C and Table 1). As expected, the growth inhibitory activity of ART analog ART1 was similar for Dd2 and the proteasome inhibitor mutants. Also as expected, Dd2(β6A117D) and Dd2(β5A49S) were 16- and >370-fold more resistant to PI01, respectively, than the parental strain Dd2. Since ATZs were more potent than ART1 against *Pf* Dd2, we reasoned that the anti-*Pf* activity of ATZs was not only derived from the ART moiety, but also from the proteasome inhibitor moiety. In agreement with that interpretation, the ATZs were as potent as PI01 in inhibiting the growth of *Pf* Dd2, and their inhibition activities were only slightly less against the mutant strains: ≤ 2.9-fold for Dd2 (β6A117D) and ≤ 3.6-fold for Dd2 (β5A49S), representing ≥ 5-fold and >100-fold improvement over PI01 against the respective strains. The deoxy-ATZ4 showed reduced activity against Dd2 comparing to PI01, and much reduced activity against Dd2 (β6A117D) and Dd2 (β5A49S). Thus, the ATZs substantially overcame resistance to the proteasome inhibitor moiety alone that were conferred by point mutations in *Pf*20S.

### Activation of ATZ produces proteasome inhibitors in a model system

We next devised a model system to explore the mechanism of action of the hybrids and test the hypothesis that degradation products of a protein covalently attached to an ATZ can lead to inhibition of 20S (Figure 2A). Because of the difficulty in obtaining large quantity of purified *Pf*20S, we used human i-20S for a proof of concept. β-casein is intrinsically unstructured and can be degraded by 20S and PA28α without a requirement for ubiquitination. We incubated β-casein with hemin and ascorbate in the presence of highly potent i-20S inhibitor ATZ2, PI01, or ART1 in two identical groups, for 4 hours at 25 °C. One group was dialyzed to remove small molecules; the other group was not dialyzed. Treated β-casein samples were then incubated with i-20S with PA28α and aliquots were removed as indicated (Figure 2B), as described ^3^. Degradation of β-casein treated with ATZ2 was markedly reduced, whereas the degradation of β-casein treated with PI01 or ART1 alone was almost complete by five hours (Figure 2B, left). As expected, in a control experiment done without removing small molecules from the reaction mixtures by dialysis, both PI01 and ATZ2 reduced the degradation of β-casein compared to ART1 (Figure 2B, right). We conducted proteomic analyses of PI01-, ART1- and ATZ2-treated β-casein in order to identify ART1 and ATZ2 modified β-casein peptides (Scheme S6 and Table S2). We unambiguously identified the peptide SLVYPFPGP^80^ from ATZ2 treated β-casein, in which proline-80 was modified by ATZ2 (Figure 2C and Table S3), and the peptide F^67^AQTQSLVYPFPGPIPN from ART1 treated β-casein, wherein phenylalanine-67 was modified by ART1 (Figure 2D and Table S4), confirming the covalent modification of β-casein by the activated artemisinin moiety in both ART1 and ATZ2. The result suggests that degradation of ART-damaged protein by the proteasome was not affected by the covalent modification by ART, yet the proteasome inhibitor-coupled ART, ATZ, inhibited the degradation of β-casein either directly or via the degradation products of ATZ-containing oligopeptides (Figure 2A).

**Figure 2.**
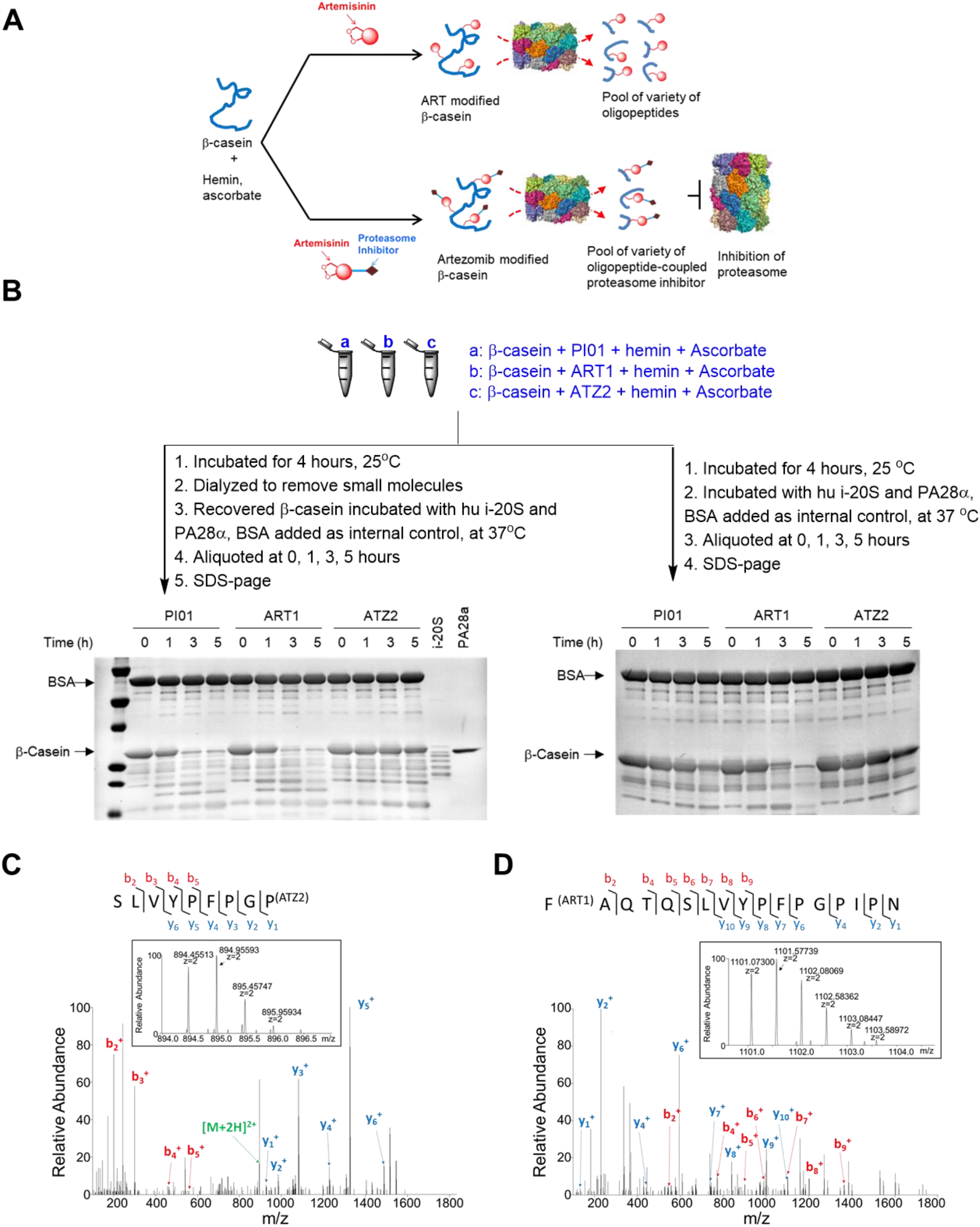
Mode of action of ATZ in the degradation of β-casein by 20S. (**A**) Illustration of degradation of β-casein by human i-20S following incubation with ART or ATZ activated by hemin and ascorbate. (**B**) Degradation of β-casein. β-casein was treated under indicated conditions (a, b or c). Left panel: after dialysis to remove the inhibitors, hemin and ascorbate, the treated β-casein was incubated with i-20S and PA28α with bovine serum albumin as an internal control. Aliquots were taken at indicated times and samples run on SDS-page and stained with Coomassie blue. Right panel: without dialysis, aliquots were taken from each reaction at indicated time points and samples run on SDS-page and stained with Coomassie blue. Representative images of three independent experiments. (C) The MS/MS spectrum of the ATZ2 modified peptide SLVYPFPGP80. The inserted mono-isotope peak at m/z 894.45557 matches the theoretical mass of the aforementioned peptide modified by ATZ2. This peptide was not observed in PI01 treated nor in ART1-treated β-casein samples through manual check. (**D**) The MS/MS spectrum of the ART1 modified peptide F67AQTQSLVYPFPGPIPN. The inserted mono-isotope peak at m/z 1101.07361 matches the mass of the aforementioned peptide modified by ART1. This peptide was not observed in PI01 treated nor in ATZ2-treated β-casein samples through manual check.

### ATZs overcome malaria resistances to ARTs

Next, we investigated if the mode of action of ATZs could circumvent the ART resistance conferred by the K13 polymorphism. Resistance to ART by the malaria parasite is not due to a shift in EC_50_, but rather is a partial resistance seen only in the early ring stage of the erythrocytic cycle. We conducted a ring-stage survival assay (RSA) with strains Cam3.I^rev^ and Cam3.I^R539T^; the latter strain has a Kelch13 polymorphism and is resistant to ART ^11^. Highly synchronized 0-3 hour ring stage parasites were treated with DMSO, dihydroartemisinin (DHA), ART1, PI01, ATZ3 or ATZ4 at the indicated concentrations for 6 hours (Figure 3). We then washed off the compounds and maintained the parasite cultures at 37 °C for a further 66 hours. Live parasites in each condition were analyzed by flow cytometry and survival expressed relative to the DMSO control. As expected, Cam3.I^R539T^ was highly resistant to DHA and slightly resistant to ART1 (Figure 3A), whereas Cam3.I^R539T^ was as susceptible as Cam3.I^rev^ to ATZ3 and ATZ4, respectively. Interestingly, Cam3.I^R539T^ was more sensitive than Cam3.I^rev^ to the proteasome inhibitor PI01. These data confirmed that ATZs can circumvent ART resistance associated with Kelch13 mutation at the early ring stages. In the ring survival assay (RSA), even DHA-sensitive parasites pulsed with DHA will eventually recover normal growth, therefore, more extended RSAs have been proposed to better reflect clinical efficacy.^27^ Accordingly, we performed an extended RSA by pulsing parasites as in the standard RSA and then monitoring parasite growth over seven days. As expected, parasites pulse-treated with DHA, PI01 or ART1 re-established normal growth. In contrast, parasites of both the ART sensitive and resistant lines had significantly lower parasitemia on day 7 (Figure 3B), indicative of a prolonged growth inhibition profile of ATZs.

**Figure 3.**
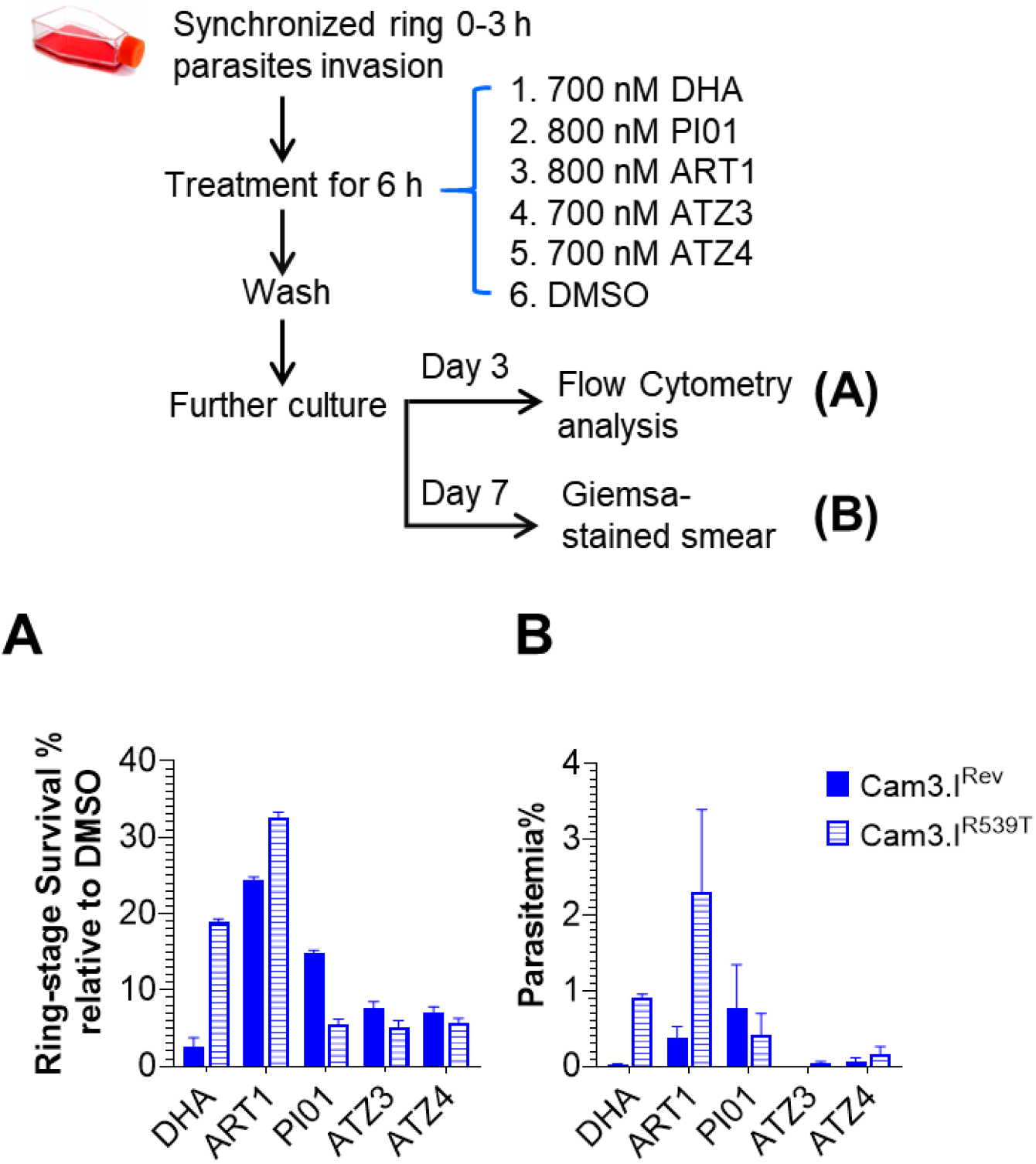
Effects of compounds in ring-stage survival assay. Red blood cells infected with highly synchronized ring-stage parasites were treated with DMSO, DHA, ART1, PI01, a 1:1 mixture of PI01 and ART1, ATZ3 or ATZ4 at indicated concentrations. After 6 h the compounds were washed off. (**A**) The parasite cultures were allowed to grow for 66 hours. Viable parasites were analyzed by flow cytometry and their numbers normalized to values for the DMSO control. (**B**) Aliquots of parasites from (A) were cultured for a further 96 hours. Parasitemia was quantified by Giemsa-stained smears.

### Mode of action of ATZ

To relate growth inhibition to pharmacodynamic effect, we used a covalently reactive, irreversible probe compound, MV151 ^28^ to label *Pf*20S. We first established that 1 hour incubation of either PI01 or ATZ4 with Dd2 *Pf*20S prior to labeling with MV151 dose-dependently blocked labeling of the *Pf*20Sβ5 of Dd2 wild type by the probe (Figure 4A). Under the same conditions, the β6A117D and β5A49S mutations prevented PI01 and ATZ4 from inhibiting the labeling *Pf*20S(β6A117D) and *Pf*20S(β5A49S) in lysates of Dd2(β6A117D) and Dd2(β5A49S) parasites (Figure 4), indicating that the mutations reduced the binding affinity of PI01 and ATZ4 to *Pf*20S β5. The seemingly contradictory results between the growth inhibition potency of ATZ3 and ATZ4 against the Dd2 and two *Pf2*0S inhibitor resistant strains (Table 1) and the enzyme labeling (Figure 4) can be explained by ATZs being transformed to intracellularly retained moieties with proteasome inhibiting potential that arose inside the parasites following activation of ATZs and that were active not only against wild type Pf20S but also against Pf20S with the β6A117D and β5A49S mutations.

**Figure 4.**
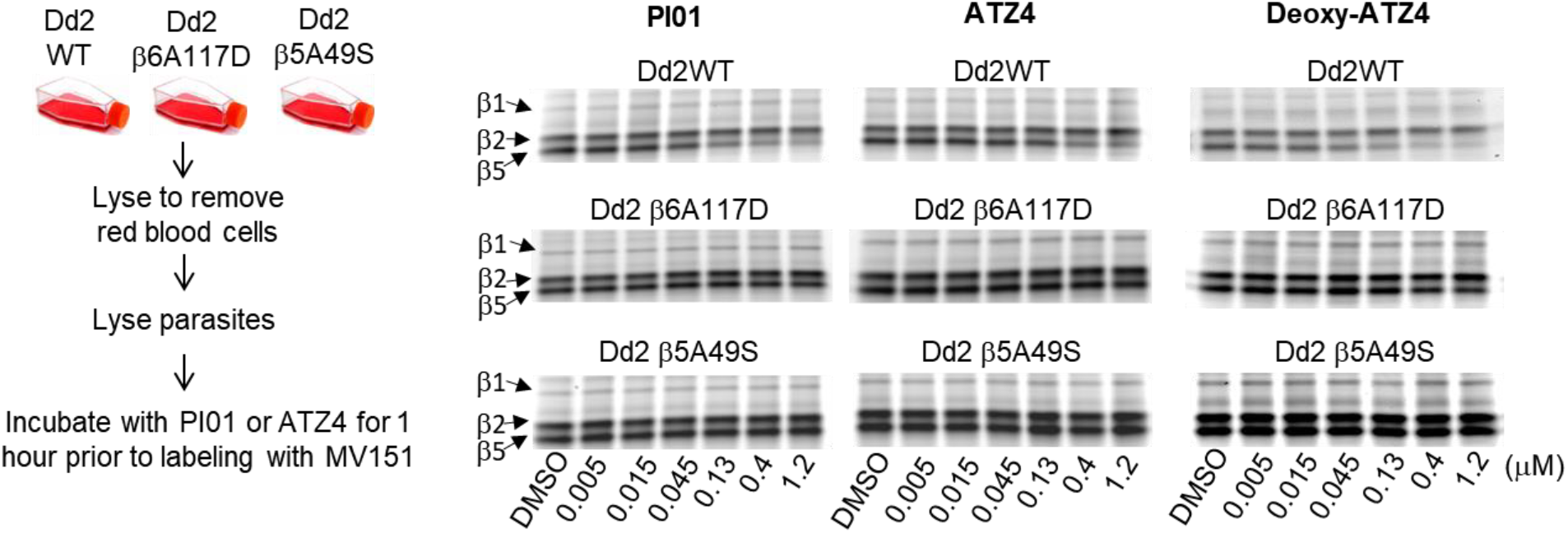
Inhibition of Pf20S, Pf20S(β6A117D) and Pf20S(β5A49S) by PI01 or ATZ4 in lysates of Dd2, Dd2(β6A117D) and Dd2(β5A49S) parasites, respectively, was assessed by their ability to block labeling of the parasites’ proteasomes by the activity-based fluorescent probe MV151 with 1 hour preincubation.

To test this hypothesis, we pulse-treated late stage Dd2 wild type, Dd2(β6A117D) and Dd2(β5A49S) parasites with DMSO as a vehicle control, DHA (700 nM), PI01 (800 nM), ART1 (800 nM), ATZ4 (700 nM), a 1:1 combination of PI01, ART1 and deoxy-ATZ4 for six hours. After thoroughly washing off the compounds and recovering the parasites from the red blood cells, we lysed the parasites and labeled *Pf*20S with MV151 (Figure 5). The labeling of *Pf*20S in lysates from parasites treated with DHA, PI01, ART1, the combination of PI01 with ART1 (1:1), and deoxy-ATZ4 was not inhibited compared to the labeling of *Pf*20S β5 by MV151 in DMSO treated samples. This demonstrated the effectiveness of the washing procedure at lowering the intracellular concentration of compounds below a functionally detectable level, because when the washing steps were omitted, PI01 and ATZ4 did block labelling of *Pf*20S β5 by MV151 (Figure S1), confirming that both of the compounds could enter red bloods cells and the parasites within them. Despite the extensive washing, *Pf*20S in the lysate from the parasites treated with ATZ4 showed substantial inhibition of labeling of *Pf*20S β5 by MV151 (Figure 4B). Moreover, this activity was similarly effective against intracellular *Pf*20S β5 with the β6A117D and β5A49S mutations (Figure 5), in contrast to the minimal inhibition of *Pf*20S β6A117D and β5A49S labeling by ATZ4 itself (Figure 4). These results are consistent with transformation of ATZ4 in the parasites to persistently-retained intracellular moieties capable of sustained inhibition of *Pf*20S β5.

**Figure 5.**
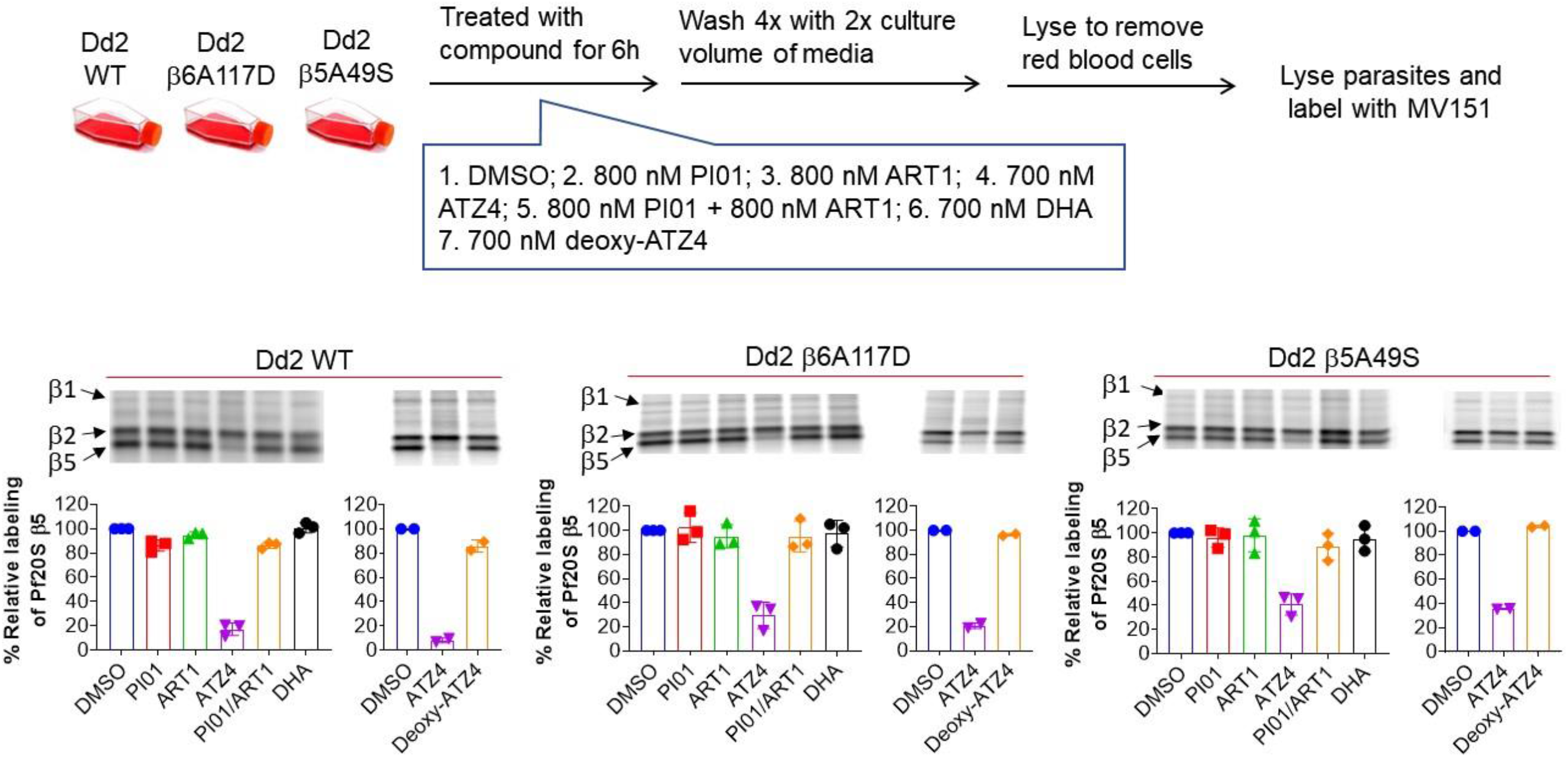
Pf20S inhibition assessed in Dd2, Dd2(β6A117D) and Dd2(β5A49S) cultures after compounds were washed off. (**A**) Scheme of experiment design. Parasites were treated with DMSO, PI01, WZ1840, ATZ4, PI01/ART1 (1:1), DHA or deoxy-ATZ4 at indicated concentrations for 6 hours and compounds were then washed off prior to hypotonic lysis of red blood cells. Parasites were then lysed and labelled with MV151. (**B**) Fluorescent scanning images of representative SDS-page gels of Pf20S labeling by MV151. Quantification of labeling was performed with ImageJ. Percentages of labeling inhibition were calculated by normalizing the ratios of density of β2 / β5 bands to that of DMSO treated. Data were means of two or three independent experiments.

**Figure 6.**
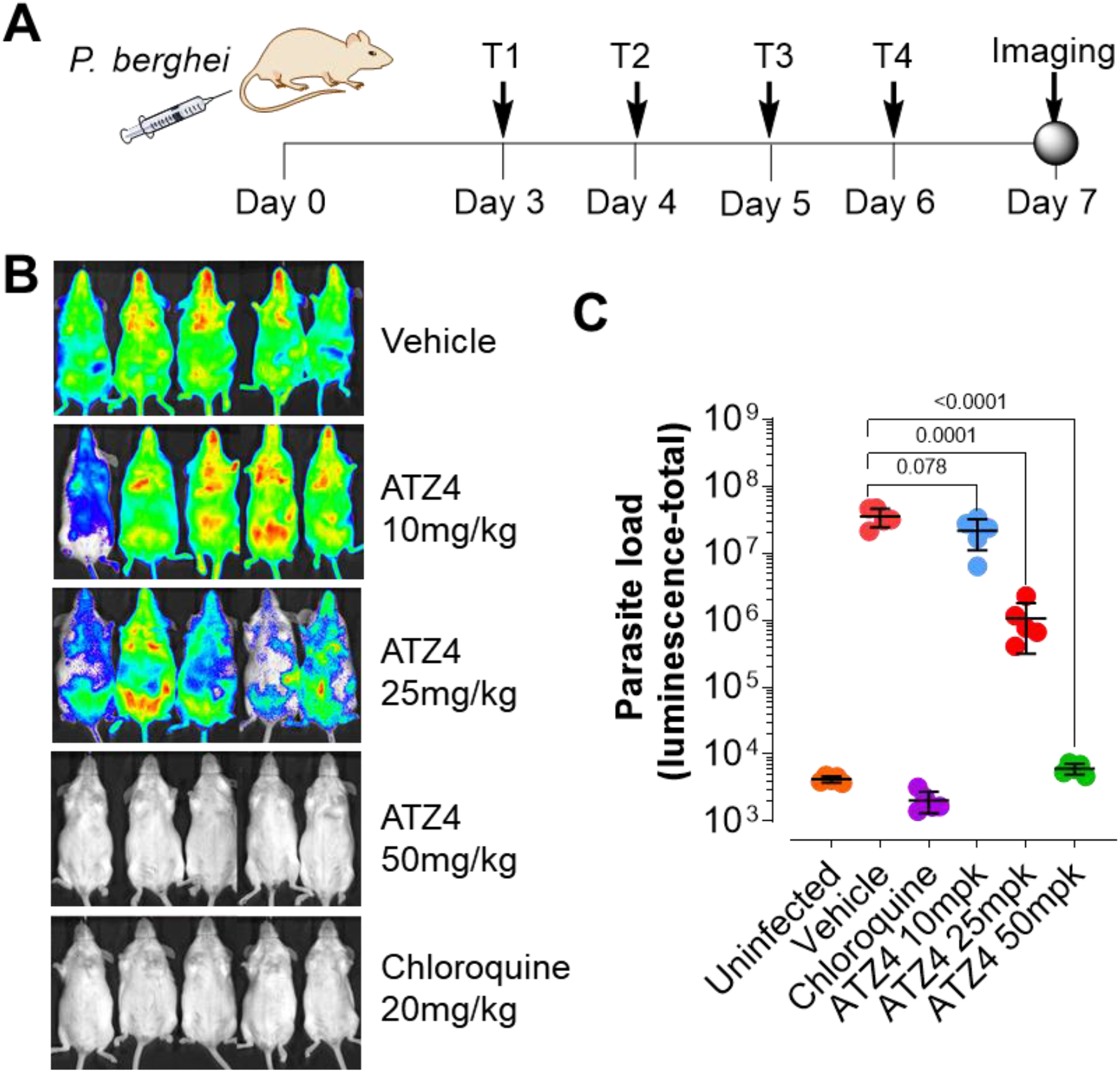
In vivo efficacy of ATZ4 in mice infected with *P. berghei* ANKA lux. A) Scheme of the in vivo efficacy study with Swiss Webster mice. *P berghei* ANKA lux was injected in the tail vein on day 0. ATZ4 was administered *i.p.* at 10, 25 and 50 mg/kg on day 3 to day 6 once daily. Chloroquine was used as positive control at 20 mg/kg via *i.p.* Luminescence density from each mouse was recorded on day 7. B) Luminescence images of mice in respective treatment groups. **C**) Quantification of parasite load in luminescence in each group in **B**). Experiments were repeated twice, and a representative is shown.

### In vivo efficacy of ATZ

To learn if an ATZ possessed in vivo efficacy, we infected mice with *P. berghei* ANKA Lux on day 0 and started once-daily treatment on day 3 for 4 days. Parasite loads in each mouse were quantified by luminescence on day 7. As expected, chloroquine at 20 mg/kg reduced the parasite load below the limit of detection. ATZ4 demonstrated dose-dependent therapeutic effects, with no detectable effect at 10 mg/kg, 1.2 log_10_ reduction in parasite load at 25 mg/kg, and > 3 log_10_ reduction at 50 mg/kg, to below the limit of detection.

## Discussion

In summary, stable, covalent conjugates of a proteasome inhibitor and an ART analog, termed ATZs, retain both proteasome inhibitory activity and the reactive alkylating activity of ART. These effects are not only synergistic against growth of *Pf* but can overcome resistance to either moiety. The ability to overcome resistance conferred by point mutations in *Pf*20S is associated with ATZ-dependent formation of proteasome-inhibitory activity that is not removed from the parasites by washing procedures that remove ATZ itself. We ascribe this more robust proteasome-inhibitory activity to the demonstrable formation of proteasomal degradation products of ATZ-damaged proteins. The oligopeptides to which the ART-derived radicals are attached appear to stabilize presentation of the proteasome inhibitory moiety of the ATZ at the *Pf*20S active site, compensating for the reduced binding affinity conferred by the point mutations. In short, with the malaria parasite’s dependence on the proteasome to remove ART-damaged proteins, ATZ hybrids hijack the parasite protein degradation machinery to create a pool of proteasome inhibitor-containing oligopeptides. Because the actions of ART and the improved action of the proteasome inhibitor are delivered by a single molecule, a single pharmacokinetic profile will preclude temporary exposure to only one of the components in the combination.

## Methods

### In vitro cultivation

*P. falciparum* laboratory lines were grown under standard conditions in RPMI 1640 medium with 0.5% Albumax II (Invitrogen), 5% hematocrit, 0.25% sodium bicarbonate and 0.1 mg/ml gentamicin. Parasites were placed in an incubator under 90% nitrogen, 5% carbon dioxide and 5% oxygen at 37 °C. Two Dd2-derived resistant strains (Dd2β5A49S and Dd2β6A117D) were developed in-house and identified as described.^17,25^

### IC50 determination

IC_50_ values of all compounds against Pf20S β5, human c-20S β5c and i-20S β5i were determined in a 96-well format as described.^17,25,26^ Briefly, 1 μL of compound in a 3-fold series dilution in DMSO at final concentrations from 100 μM to 0.0017 μM was mixed with 99 μL of reaction buffer containing the corresponding proteasome, substrate and activator in a black 96-well plate with a solid bottom. Buffer 50 mM Tris, 5 mM MgCl_2_, 1 mM DTT, pH 7.4 was for Pf20S β5 and buffer 20 mM HEPES, 0.5 mM EDTA and 0.1 mg/mL BSA, pH7.5 for human β5c and β5i. The fluorogenic substrate suc-LLVY-AMC was used for Pf20S and c-20S at final concentration 25 μM, and Ac-ANW-AMC was used as substrate of i-20S at final concentration 15 μM. Activator PA28α at final concentration of 12 nM was used for Pf20S assay in the presence of 0.5 μM of WLW-VS, whereas 0.02% SDS was used in the assays for c-20S and i-20S. Final concentrations of Pf20S, c-20S and i-20S were 1 nM, 0.2 nM and 0.4 nM, respectively. The fluorescence of the hydrolyzed AMC at λex=360nm and λex= 460 nm in each well was followed for 1-2 hours. Linear ranges of the time course were used to calculate the velocities in each well, which were fit to a dose-dependent inhibition equation to estimate the IC_50_ values in PRISM (GraphPad).

### Anti-malarial activity in erythrocytic stage

Parasite growth inhibition assays were performed as reported.^25^ Drug assays were performed on parasites cultured in sterile 96-well plates at a total 200 μL volume per well and a 0.5% initial parasitemia and 2% hematocrit. Plates were placed in an airtight chamber flushed with 5% oxygen, 5% carbon dioxide and 90% nitrogen for 72 hours. Plates were then placed in the −80 °C freezer to promote cell lysis upon thawing. When thawing was complete, 100 μL of SYBR Green diluted in lysis buffer (0.2 μL SYBR Green per ml lysis buffer) was added to each well and the plates were shaken in the dark at room temperature for 1 h. Fluorescence was then recorded in a SpectraMax Gemini plate reader using λex=490 nm / λem=530 nm. Data analysis was performed with Graphpad Prism software. Counts were normalized and plotted by non-linear regression to yield EC_50_ values.

### Ring Survival Assay

Ring survival assays (RSA) were performed as described ^11^. Parasite cultures, IPC5202 (Cam3.I^R539T^), an artemisinin resistant parasite line from Cambodia, and the genetically engineered artemisinin sensitive revertant Cam3.I^rev^, were synchronized several times with 5% sorbitol and then a Percoll-sorbitol gradient was used to obtain tightly synchronized late stage parasites. Isolated late stage parasites were then allowed to reinvade fresh red blood cells for three hours ring stage parasites were confirmed by microscopy before the cultures were again subjected to 5% sorbitol to obtain 0-3 hour rings. The isolated ring stage cultures were then plated into a 96 well plate at 0.5% parasitemia at the corresponding drug concentrations: DHA 700 nM, PI01 800 nM, ART1 800 nM, ATZ3 700 nM, ATZ4 700 nM. Plates were incubated at 37 °C in standard gas conditions for six hours before the plates were spun and washed to remove medium with compound and replenished with fresh medium. Plates were then incubated for an additional 66 hours and parasite growth was then assessed using flow cytometry and nucleic acid stains HO (Hoechst 33342) and TO (thiazole orange).

### Parasite regrowth assay

Parasite were synchronized as described for the RSA and treated with the same drug concentrations for six hours in a 96 well plate. After six hours of drug/compound exposure and washing, the 200 μL culture was then transferred to a 3 mL culture which was monitored for seven days. Parasitemia was checked by microscopy on day 7.

### Inhibition of Pf20S, Pf20S(β6A117D0 and Pf20S(β5A49S) by PI01 and ATZ4

Cell free lysates of *P. falciparum* Dd2 wild-type and two Dd2-derived resistant (Dd2◻5A49S and Dd2β6A117D) were used. 5 - 10 μg of total lysate proteins were incubated with PI01 or ATZ4 at the indicated concentrations for 1 h at 37 °C prior to addition of MV151 at a final concentration of 2 μM and incubated for a further 1 h at 37 °C. The samples were then heated with 4X SDS loading buffer at 95 °C for 10 min and run on 12% Novex™ Bis-Tris Protein Gels. The gels were rinsed with double distilled H_2_O and then scanned on the Typhoon Scanner.

### Intraparasitic hybrid activation and proteasome inhibition assay

*Pf* Dd2, Dd2(β6A117D) and Dd2(β5A49S) parasites were grown synchronized to a high parasitemia (5-8%). At the early trophozoite stage, 5 mL of parasite-infected red blood cells were exposed to DMSO, PI01 (800 nM), ART1 (800 nM), ATZ4 (700 nM), and a mixture of ART1 and PI01 in a 1:1 ratio both at 800 nM for 6 hours. After centrifugation at 3600 rpm for 3 minutes, the supernatant was removed, and red cells were washed with complete media once and resuspended in 14 mL of fresh complete media. The cultures were then placed back in the incubator and shaken for 10 minutes. The wash procedure was repeated 4 times. After the last wash, the red blood cell pellets were placed on ice and washed with PBS 1 mL once and then lysed with 10% saponin to obtain parasite pellets. Parasite pellets were kept on ice and washed with cold PBS until supernatant was clear (approximately 3 times). Pellets were stored at −80 °C until analysis. The frozen *Pf* pellets were thawed on ice and resuspended in 2 × pellet volume of lysis buffer containing 20 mM Tris-HCl, 5 mM MgCl_2_ and 1 mM DTT, pH 7.4. The mixtures were kept on ice for 1 h and vigorously vortexed every 5 min, then centrifuged at 15000 rpm for 20 min at 4 °C. The supernatants were collected and their concentrations were determined by BCA protein assay. Equal amounts of lysates were incubated with MV151 at a final concentration of 2 μM for 1 h at 37 °C in a 1.5 mL Eppendorf tube wrapped in aluminum foil. The samples were then heated with 4X SDS loading buffer at 95 °C for 10 min and run on a 12% Novex™ Bis-Tris Protein Gel with MOPS SDS running buffer. The gel was rinsed with double distilled H_2_O and scanned at the TAMRA channel on a Typhoon Scanner (GE Healthcare). In an alternative procedure, wash steps of parasite-infected red blood cells in the above-stated procedure were omitted to examine the permeability of PI01 and ATZ4. However, parasite pellets were thoroughly washed to remove red blood cell constituents to avoid interferences in the MV151 labeling assays.

### HepG2 cell viability assay

HepG2 cells were plated in 96-well plate (5,000 cells per well) format and treated with various concentrations of test compounds or DMSO for 72 h. Cell viability was measured using CellTiter-Glo^®^ Assay (Promega) as per the manufacturer’s instructions.

### Degradation of the modified β-casein by i-20S

β-casein dissolved in PBS (10 μM) was incubated with 100 μM of PI01, ART1 or ATZ2 in the presence of sodium ascorbate (200 μM) and hemin (100 μM) at r.t. for 4 h. The samples were transferred to Slide-A-Lyzer MINI Dialysis Devices (10K MWCO, Thermo Scientific™ 88401) and placed into tubes containing the dialysis buffer (20 mM HEPES and 0.5 mM EDTA, pH7.5) and dialyzed overnight at 4 °C with fresh dialysis buffer changing every 4 h. After dialysis, the samples were collected and further incubated with i-20S (50 nM), PA28α (0.5 μM), and bovine serum albumin (10 μM) at 37 °C. Aliquots from the reaction mixtures were removed at designated time intervals, mixed with SDS sample loading buffer, and were run on a SDS-PAGE (4-20%, Tris-Glycine) and stained with Coomassie blue. For the control experiment in **Figure 2B**, samples were prepared with the same method above, except that dialysis step was skipped.

### Protein Sample Preparation for Mass Spectrometry Analysis

β-casein was treated as aforementioned. After removing the inhibitors, hemin and ascorbate by dialysis, the treated β-casein samples were run on SDS-page and stained with Coomassie blue G-250. The gel bands of β-casein were cut into pieces. Samples were reduced with 5 mM dithiothreitol in 50 mM ammonium bicarbonate buffer for 50 min at 55℃ and then dried by acetonitrile. Next, the samples were alkylated with 12.5 mM iodoacetamide in 50 mM ammonium bicarbonate buffer for 45 min in the dark at room temperature and dried by acetonitrile. The samples were then digested by trypsin or chymotrypsin at 37 ℃ overnight. The digestion was stopped with 10% trifluoracetic acid, after which the digested peptides were extracted twice with 1% formic acid in 50% acetonitrile aqueous solution, and then evaporated to dryness on a Speedvac and resuspended in 20 μL of formic acid/H_2_O (v:v = 0.1%/99.9%) with sonication.

### LC-MS/MS

For LC-MS/MS analysis, the fragment peptides were separated by a 120-min gradient elution method at a flow rate of 0.3 μL/min with a Thermo-Dionex Ultimate 3000 HPLC system that is directly interfaced with a Thermo Orbitrap Fusion Lumos mass spectrometer. The analytical column was a homemade fused silica capillary (75 μm inner-diameter, 150 mm length; Upchurch, Oak Harbor, WA, USA) packed with C-18 resin (pore size 300 Å, particle size 5 μm; Varian, Lexington, MA, USA). Mobile phase A was 0.1% formic acid in water, and mobile phase B is 100% acetonitrile and 0.1% formic acid. The Thermo Orbitrap Fusion Lumos mass spectrometer was operated in the data-dependent acquisition mode using Xcalibur 4.0.27.10 software. A single full-scan mass spectrum was done in the Orbitrap (300 −1500 m/z, 120,000 resolution). The spray voltage was 1850 V and the Automatic Gain Control (AGC) target was 200,000. This was followed by 3-second data-dependent MS/MS scans in an ion routing multipole at 30% normalized collision energy (HCD). The charge state screening of ions was set at 1-8. The exclusion duration was set at 8 seconds. Mass window for precursor ion selection was set at 2 m/z. The MS/MS resolution was 15,000. The MS/MS maximum injection time was 150 ms and the AGC target was 50,000.

### Mass Data Processing

Data were searched against the bovine casein database from the Uniprot by using Proteome Discoverer 1.4 software (Thermo Scientific) and peptide sequences were determined by matching protein database with the acquired fragmentation pattern by SEQUEST HT algorithm. The following search parameters were used: the precursor mass tolerance was set to 10 ppm and fragment mass tolerance was 0.02 Da; No-Enzyme (Unspecific); Modification (MOD) A_1_ (811.39624 Da, **Table S2**, any amino acids), MOD A_2_ (751.37511 Da, **Table S2**, any amino acids), MOD B_1_ (325.18837 Da, **Table S2**, any amino acids), MOD B_2_ (265.16725 Da, **Table S2**, any amino acids), carbamidomethyl of cysteines (+57.02146 Da), oxidation of methionines (+15.99492 Da) as the variable modifications. Only peptides with the strict target false discovery rate (FDR) below 1% were considered as high-confidence hits.

### In vivo efficacy of ATZ4 in mice infected with *P. berghei*

Female 4-6 weeks old Swiss-Webster mice were infected on day 0 via *i.p.* injection with 10^3^ (~5×10^−5^ % infection) *P. berghei-*Luccon ^29^ infected erythrocytes. Treatments started from day 3 to day 6 for 4 days. Chloroquine (20 mg/kg) in 0.5% hydroxymethyl cellulose, 0.4% Tween-80 and ATZ4 (10, 25, 50 mg/kg) in 5% DMSO, 5% Tween-80 sterile water were administered via *i.p.* once daily. On day 7, anesthetized mice were i.p. injected 150 mg/kg of D-luciferin potassium-salt (Goldbio, LUCK-100) dissolved in PBS. Mice were imaged 5 to 10 minutes after injection of luciferin with an IVIS 100 (Xenogen, Alameda, CA) and the data acquisition and analysis were performed with the software LivingImage (Xenogen).

## ASSOCIATED CONTENT

### Supporting Information

Supplementary schemes, figure and supplementary tables; detailed experimental procedures including synthesis and characterization.

## Author Contributions

Conceptualization: W.Z. and G.L. Supervision: G.L. and L.A.K. Investigation: W.Z., J.Y.L., C.Y., H.Z., J.C.H., R.W., S.Z., J.S., G.S., A.R., H.D., C.F.N., L.A.K. and G.L. Methodology: W.Z., J.Y.L., C.Y., H.Z., R.W., A.R., G.S., H.D., C.F.N., L.A.K. and G.L. Writing: W.Z., C.Y., H.D., C.F.N., L.A.K. and G.L. All authors read, edited and approved the final manuscript.

## Data Availability

All study data are included in the article and/or SI Appendix.

## Notes

The authors declare the following competing financial interest(s): Cornell University has filed a provisional patent application on these artemisinin proteasome inhibitor hybrids. G. Lin, W. Zhan, H. Zhang, C. Nathan and L. Kirkman are listed as inventors. This statement will be revised once more information is available before publication.

## ACKNOWLEDGMENT

This work is supported by R21AI153485 (GL), in part by R01AI143714 (GL), in part by a Brockman Medical Foundation Medical Research Grant (LK) and by the Milstein Program in Chemical Biology and Translational Medicine. This research was funded in part through the NIH/NCI Cancer Center Support Grant P30 CA008748.The Department of Microbiology and Immunology is supported by the William Randolph Hearst Foundation.

## ABBREVIATIONS

ART: artemisinin
ACT: artemisinin combination therapy
ATZ: artezomib – artemisinin-proteasome inhibitor hybrid
c-20S: human constitutive proteasome
i-20S: human immunoproteasome
*Pf*: *Plasmodium falciparum*
*Pf*20S: *Pf* proteasome
Dd2(β6A117D): *Pf* Dd2 strain with a mutation A117D on β6 subunit of *Pf*20S
Dd2(β5A49S): *Pf* Dd2 strain with a mutation A49S on β5 subunit of *Pf*20S;
*Pf*20S(β6A117D): *Pf*20S with a mutation A117D on β6 subunit
*Pf*20S(β5A49S): *Pf*20S with a mutation A49S on β5 subunit

