## Supporting information for "Artemisinin-based hybrids produce intracellular proteasome inhibitors that overcome resistance in *Plasmodium falciparum*"

**Table of Contents**

**S2 – Supplementary Schemes and Figure**

**S6 – Supplementary Tables**

**S8 – Methods**

**S20 – References**

**S22 – $^{1}$H and $^{13}$C NMR Spectra**

**S36 – LCMS**

**S45 – HRMS**

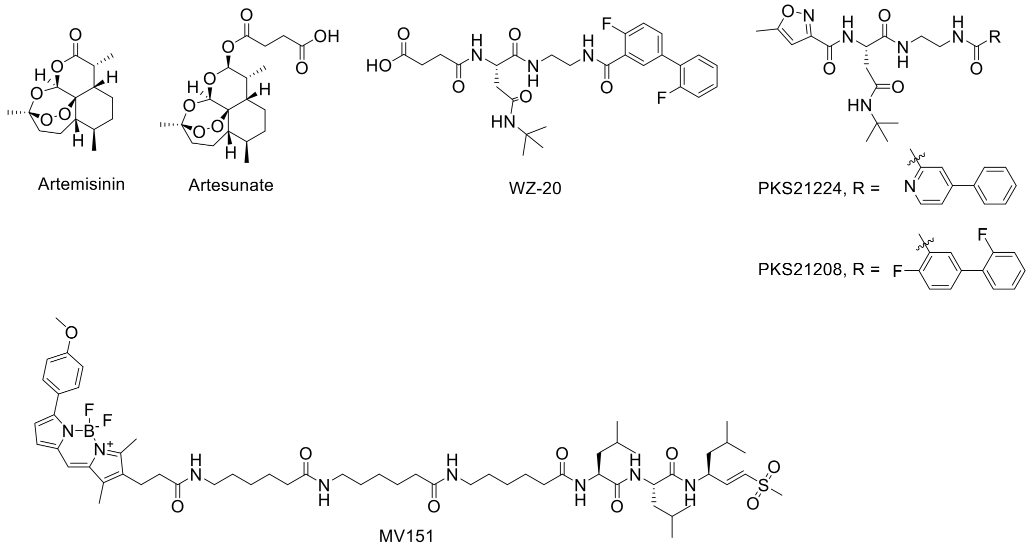

**Scheme S1**. Structures of Artemisinin, Artesunate, AsnEDA-based proteasome inhibitors PKS21224 and PKS21208, control compound WZ-20 and proteasome labeling probe MV151.

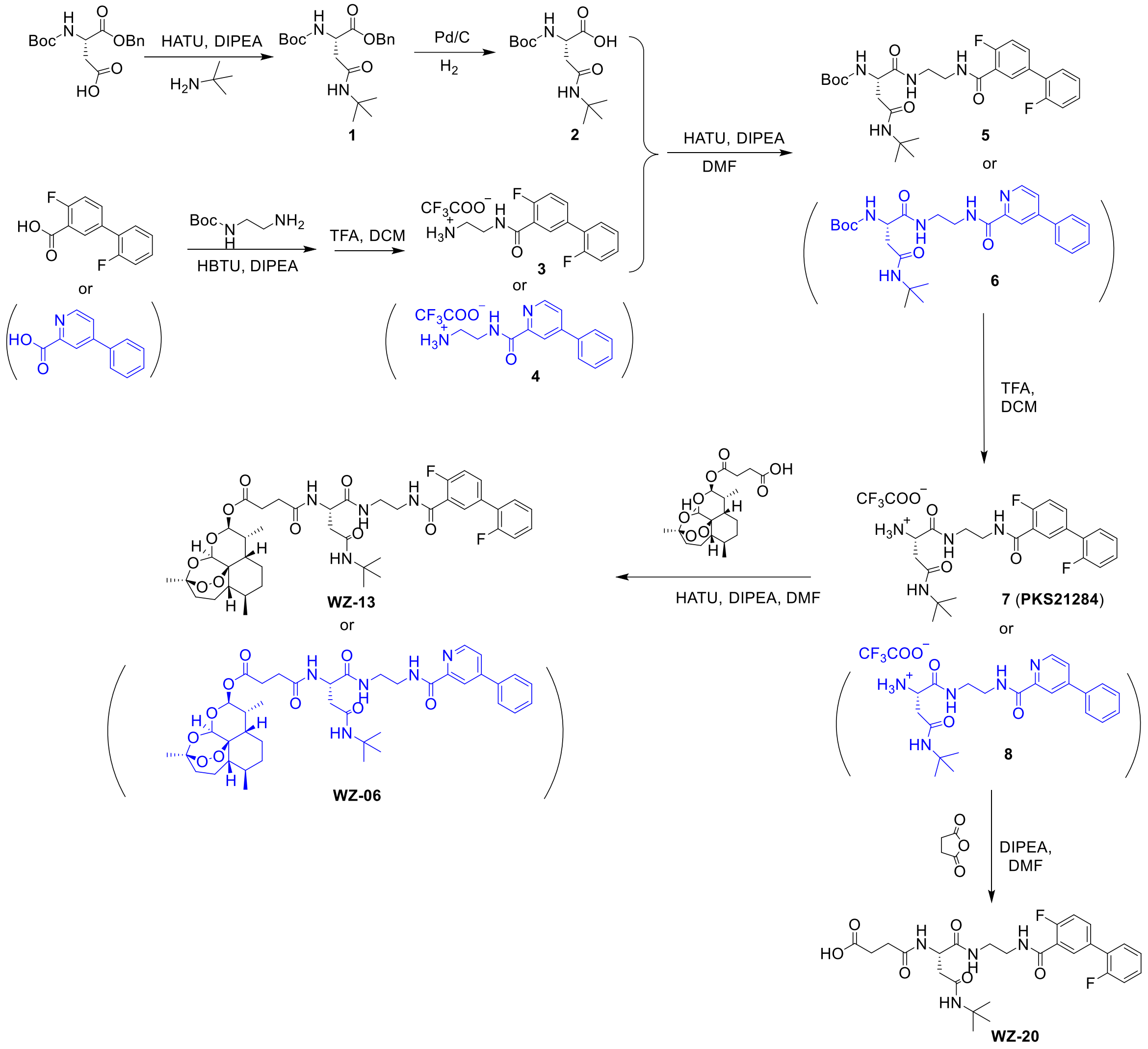

**Scheme S2**. Synthetic route of Artesunate-based hybrids WZ-13 and WZ-06 and control WZ-20.

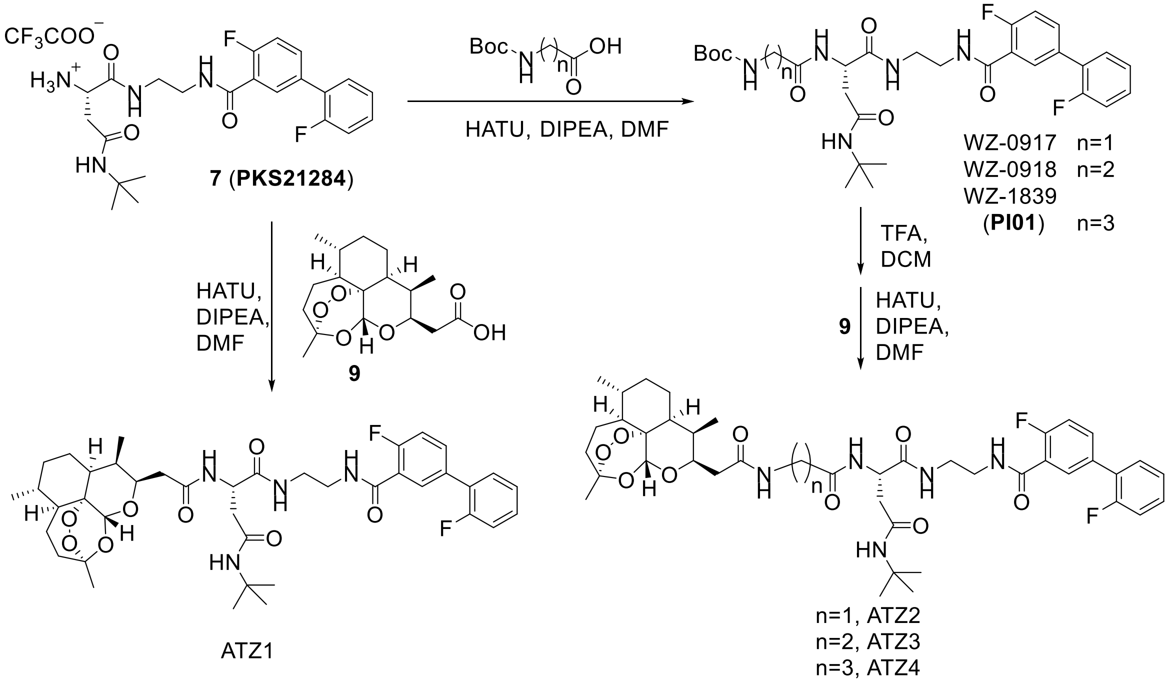

**Scheme S3**. Synthetic route of ART-based hybrids ATZ1~4.

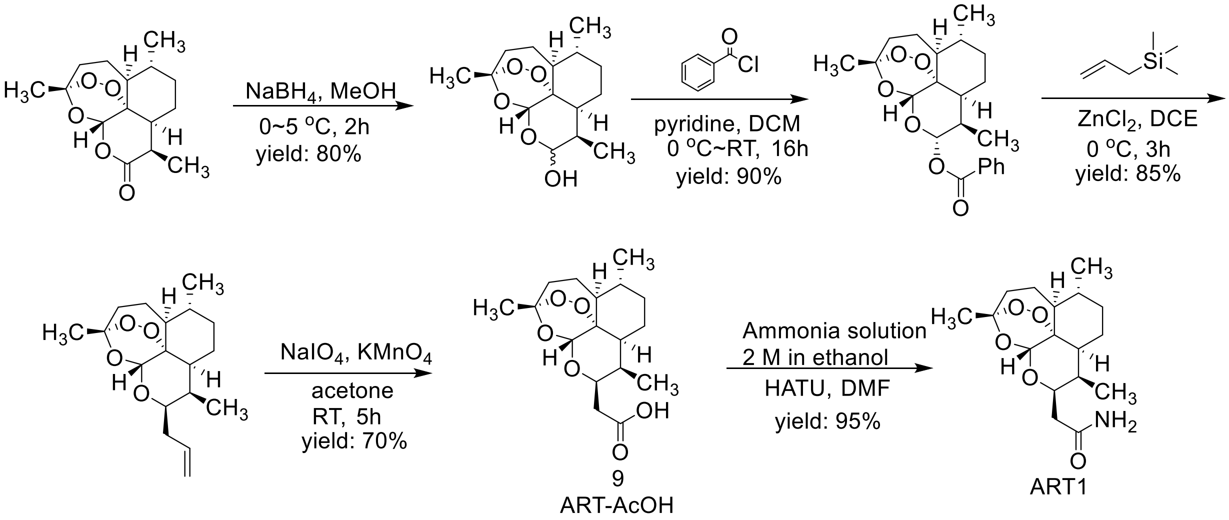

**Scheme S4**. Synthetic route of ART-AcOH (9) and ART1.

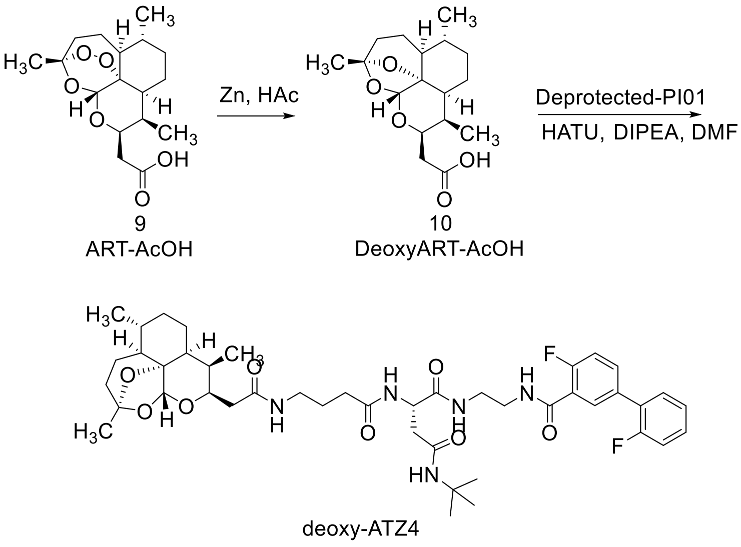

**Scheme S5**. Synthesis of deoxy-ATZ4.

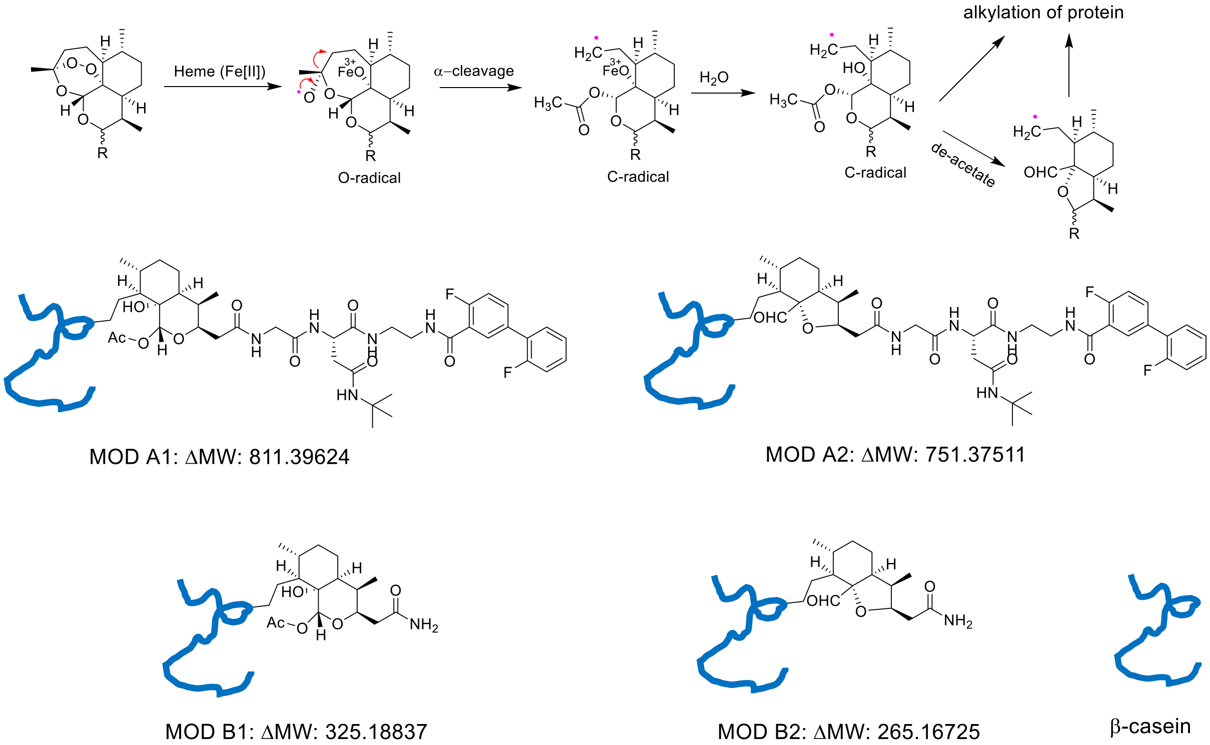

**Scheme S6**. Heme-induced activation of the endoperoxides, yielding reactive radical intermediates of ART1 and ATZ2 capable of two types of covalent modification of β-casein.

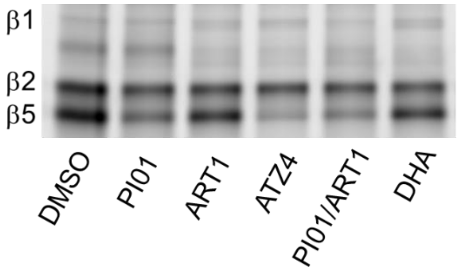

**Figure S1**. Labelling inhibition of *Pf*20S in Dd2 parasites treated with DMSO, PI01, ART1, ATZ4, PI01/ART1 (1:1) or DHA, assessed by their ability to block labeling of the parasites' proteasomes by MV151. Parasites were treated with indicated compounds for 6 hours and extracellular compounds were removed prior to hypotonic lysis of red blood cells.

**Table S1**. $IC_{50}$ values of compounds against β5 of *Pf*20S, i-20S and c-20S.

| | $IC_{50}$ (nM) | | | $EC_{50}$ (nM) |
|---|---|---|---|---|
| | *Pf*20S | i-20S | c-20S | *Pf* 3D7 |
| Artesunate | >100,000 | >100,000 | >100,000 | 6.2 ± 4.0 |
| WZ-20 | 6.0 ± 0.005 | 3,490 ± 1,160 | 34,300 ± 8,600 | 50.8 ± 0.7 |
| WZ-06 | 13.5 ± 7.6 | 1,100 ± 50 | 6,610 ± 1,090 | 3.6 ± 0.01 |
| WZ-13 | 5.5 ± 3.8 | 158 ± 50 | 440 ± 110 | 2.5 ± 0.7 |

**Table S2**. Two types of modification of β-casein by ATZ2 or ART1 as shown in **Scheme S4**. (A) For ATZ2, MOD $A_1$ Δmass = 811.39624 and MOD $A_2$ Δmass = 751.37511 ($A_1$-acetate); (B) For ART1, MOD $B_1$ Δmass =325.18837 and MOD $B_2$ Δmass =265.16725 ($B_1$-acetate).

| Tagged molecules | Modification | Molecular formula | Theoretical mass (monoisotopic) |
|---|---|---|---|
| ATZ2 | MOD $A_1$ | $C_{42}H_{55}F_2N_5O_9$ | 811.39624 |
| | MOD $A_2$ | $C_{40}H_{51}F_2N_5O_7$ | 751.37511 |
| ART1 | MOD $B_1$ | $C_{17}H_{27}NO_5$ | 325.18837 |
| | MOD $B_2$ | $C_{15}H_{23}NO_3$ | 265.16725 |

**Table S3**. Fragment ions of peptide (SLVYPFPGP[80]) modified by ATZ2. Red and blue numbers indicate fragments that were matched with theoretical masses of corresponding fragments; black numbers indicate fragments not detected.

| #1 | $b^+$ | $b^{2+}$ | Seq. | $y^+$ | $y^{2+}$ | #2 |
|---|---|---|---|---|---|---|
| 1 | 88.03931 | 44.52329 | S | | | 9 |
| 2 | 201.12338 | 101.05533 | L | 1700.87809 | 850.94268 | 8 |
| 3 | 300.19180 | 150.59954 | V | 1587.79402 | 794.40065 | 7 |
| 4 | 463.25512 | 232.13120 | Y | 1488.72560 | 744.86644 | 6 |
| 5 | 560.30789 | 280.65758 | P | 1325.66228 | 663.33478 | 5 |
| 6 | 707.37631 | 354.19179 | F | 1228.60951 | 614.80839 | 4 |
| 7 | 804.42908 | 402.71818 | P | 1081.54109 | 541.27418 | 3 |
| 8 | 861.45055 | 431.22891 | G | 984.48832 | 492.74780 | 2 |
| 9 | | | P-ATZ2 | 927.46685 | 464.23706 | 1 |

**Table S4**. Fragment ions of peptide (F[67]AQTQSLVYPFPGPIPN) modified by ART1. Red and blue numbers indicate fragments that were matched with theoretical masses of corresponding fragments; black numbers indicate fragments not detected.

| #1 | $b^+$ | $b^{2+}$ | Seq. | $y^+$ | $y^{2+}$ | #2 |
|---|---|---|---|---|---|---|
| 1 | 473.26407 | 237.13567 | F-ART1 | | | 17 |
| 2 | 544.30119 | 272.65423 | A | 1728.89561 | 864.95144 | 16 |
| 3 | 672.35977 | 336.68352 | Q | 1657.85849 | 829.43288 | 15 |
| 4 | 773.40745 | 387.20736 | T | 1529.79991 | 765.40359 | 14 |
| 5 | 901.46603 | 451.23665 | Q | 1428.75223 | 714.87975 | 13 |
| 6 | 988.49806 | 494.75267 | S | 1300.69365 | 650.85046 | 12 |
| 7 | 1101.58213 | 551.29470 | L | 1213.66162 | 607.33445 | 11 |
| 8 | 1200.65055 | 600.82891 | V | 1100.57755 | 550.79241 | 10 |
| 9 | 1363.71387 | 682.36057 | Y | 1001.50913 | 501.25820 | 9 |
| 10 | 1460.76664 | 730.88696 | P | 838.44581 | 419.72654 | 8 |
| 11 | 1607.83506 | 804.42117 | F | 741.39304 | 371.20016 | 7 |
| 12 | 1704.88783 | 852.94755 | P | 594.32462 | 297.66595 | 6 |
| 13 | 1761.90930 | 881.45829 | G | 497.27185 | 249.13956 | 5 |
| 14 | 1858.96207 | 929.98467 | P | 440.25038 | 220.62883 | 4 |
| 15 | 1972.04614 | 986.52671 | I | 343.19761 | 172.10244 | 3 |

| | | | | | | |
|---|---|---|---|---|---|---|
| 16 | 2069.09891 | 1035.05309 | P | 230.11354 | 115.56041 | 2 |
| 17 | | | N | 133.06077 | 67.03402 | 1 |

**Table S5.** HPLC purity and SMILES for the final compounds

| ID | Purity (%) | SMILES |
|---|---|---|
| WZ-06 | >95 | [H][C@@]12CC[C@]3(OO[C@@]14[C@@](CC[C@H]2C)([C@H]([C@@H](O[C@]4([H])O3)OC(CCC(N[C@@H](CC(NC(C)(C)C)=O)C(NCCNC(C5=CC(C6=CC=CC=C6)=CC=N5)=O)=O)=O)=O)C)[H])C |
| WZ-13 | >95 | [H][C@@]12CC[C@]3(OO[C@@]14[C@@](CC[C@H]2C)([C@H]([C@@H](O[C@]4([H])O3)OC(CCC(N[C@@H](CC(NC(C)(C)C)=O)C(NCCNC(C5=CC(C6=C(F)C=CC=C6)=CC=C5F)=O)=O)=O)=O)C)[H])C |
| WZ-20 | >95 | OC(CCC(N[C@@H](CC(NC(C)(C)C)=O)C(NCCNC(C1=CC(C2=C(F)C=CC=C2)=CC=C1F)=O)=O)=O)=O |
| PI01 | >95 | O=C(NCCNC([C@H](CC(NC(C)(C)C)=O)NC(CCCNC(OC(C)(C)C)=O)=O)=O)C1=C(F)C=CC(C2=C(F)C=CC=C2)=C1 |
| ART1 | >95 | C[C@]1(O[C@@]2([H])O[C@H](CC(N)=O)[C@@H]3C)CC[C@]4([H])[C@@]2(OO1)[C@@]3([H])CC[C@H]4C |
| ATZ1 | >95 | C[C@]1(O[C@@]2([H])O[C@H](CC(N[C@@H](CC(NC(C)(C)C)=O)C(NCCNC(C3=C(F)C=CC(C4=C(F)C=CC=C4)=C3)=O)=O)=O)[C@@H]5C)CC[C@]6([H])[C@@]2(OO1)[C@@]5([H])CC[C@H]6C |
| ATZ2 | >95 | C[C@]1(O[C@@]2([H])O[C@H](CC(NCC(N[C@@H](CC(NC(C)(C)C)=O)C(NCCNC(C3=C(F)C=CC(C4=C(F)C=CC=C4)=C3)=O)=O)=O)=O)[C@@H]5C)CC[C@]6([H])[C@@]2(OO1)[C@@]5([H])CC[C@H]6C |
| ATZ3 | >95 | C[C@]1(O[C@@]2([H])O[C@H](CC(NCCC(N[C@@H](CC(NC(C)(C)C)=O)C(NCCNC(C3=C(F)C=CC(C4=C(F)C=CC=C4)=C3)=O)=O)=O)=O)[C@@H]5C)CC[C@]6([H])[C@@]2(OO1)[C@@]5([H])CC[C@H]6C |
| ATZ4 | >95 | O=C(NCCNC([C@@H](NC(CCCNC(C[C@H]([C@@H]1C)O[C@]2([H])O[C@@]3(C)CC[C@]4([H])[C@@]2(OO3)[C@@]1([H])CC[C@H]4C)=O)=O)CC(NC(C)(C)C)=O)=O)C5=C(F)C=CC(C6=C(F)C=CC=C6)=C5 |
| Deoxy-ATZ4 | >95 | C[C@@](O[C@@]1([H])O[C@H](CC(NCCCC(N[C@@H](CC(NC(C)(C)C)=O)C(NCCNC(C2=C(F)C=CC(C3=C(F)C=CC=C3)=C2)=O)=O)=O)=O)[C@@H]4C)(O5)CC[C@]6([H])[C@@]15[C@@]4([H])CC[C@H]6C |

#### Methods

##### Materials

The human constitutive proteasome (c-20S, Catalog No.: E-360), human 20S immunoproteasome (i-20S, Catalog No.: E-370), and recombinant human PA28 activator alpha subunit (Catalog N0.:

E-381) were purchased from Boston Biochem. The *P. falciparum* 20S proteasome (*Pf*20S) was purified as reported.(1) β-Casein (Catalog No.: C6905), bovine serum albumin (BSA, Catalog No.: 3117057001), hemin (Catalog No.:51280), sodium ascorbate (Catalog No.: PHR1279), artemisinin (ART, Catalog No.: 361593) and artesunate (ASU, Catalog No.: A3731) were purchased from Sigma-Aldrich. Trypsin (V528A) or chymotrypsin (V106A) were purchased from Promega. Proteasome β5 substrate suc-LLVY-AMC and β5i substrate Ac-ANW-AMC were purchased from Boston Biochem. Activity-based probe MV151 was synthesized as reported.(2) β2-specific inhibitor WLW-VS was prepared following the reported method.(3)

The following parasite strains were obtained through BEI Resources, NIAID, NIH: *Plasmodium falciparum*; strain IPC 5202, MRA-1240 contributed by Didier Ménard and strain Cam3.I$^{rev}$, MRA-1252, contributed by David A. Fidock. IPC 5202 is also referred to as Cam 3.1$^{R539T}$ and has shown resistance to artemisinin and harbors a K13 propeller mutation of R539T. Cam3.I$^{rev}$ is a K13-propeller revertant mutant of the original Cam 3.1 strain.

**Chemicals and Spectroscopy**. Unless otherwise stated, all commercially available materials were purchased from Aldrich, P3 BioSystems, Combi-Blocks or other vendors and were used as received. All reactions in aprotic solvents were performed under argon in oven-dried glassware. Reaction progress was monitored on a Waters Acquity Ultra Performance Liquid Chromatography (UPLC/MS). All HPLC purifications were performed on a Waters Autopure (mass directed purification system) equipped with a Prep C18 5μm OBD (19 X 150 mm) column. $^{1}$H- and $^{13}$C-NMR spectra were acquired on a Bruker DRX-500 spectrometer. Chemical shift δ is expressed in parts per million, with the solvent resonance as an internal standard ($CDCl_3$, $^{1}$H: 7.26; $^{13}$C: 77.16 ppm; DMSO-*d6*, $^{1}$H: 2.50 ppm; $^{13}$C: 39.52 ppm). NMR data are reported as follows: chemical shift, multiplicity (br = broad, d = doublet, q = quartet, m = multiplet, s = singlet, t = triplet), coupling constant, and integration. High Resolution Mass Spectra (HRMS) of final products were collected on a PE SCIEX API 100.

**General Procedure for HATU mediated amide bond formation**: The solution of carboxylic acid (1.1 equivalent) and *O*-(7-azabenzotriazole-1-yl)-*N*,*N*,*N*,*N*'-tetramethyluronium hexafluorophosphate (HATU, 1.5 equivalent) in anhydrous DMF was cooled to 0 °C on ice bath prior to addition of  amine (1 equivalent) and Hünig base (2 - 3 equiv) sequentially to the reaction mixture at 0 °C. The reaction mixture was stirred at 0 °C and the reaction progress was monitored on an UPLC. The reactions usually completed in 2 – 3 h. After the completion of reaction, cold water was added to quench the reaction, and the mixture was stirred for 15 minutes. Mixture was extracted twice with ethyl acetate or dichloromethane, the combined organic layer was washed with 1N HCl, water, saturated $NaHCO_3$ solution and saturated brine solution. Organic layer was further dried over anhydrous $Na_2SO_4$ and evaporated under vacuum to give product, which was used directly used in next step without further purification or purified by flash column chromatography or preparative LCMS.

**General Procedure for Boc-Deprotection**. The solution of substrate in dichloromethane was cooled to 0 °C prior to addition of trifluoroacetic acid (20% v/v with respect to dichloromethane) dropwise at 0 °C with stirring. The mixture was allowed to warm to room temperature gradually over a period of 1 h and stirred until the completion of reaction (2 – 3 h; monitored on an UPLC). Excess trifluoroacetic acid and dichloromethane were evaporated and the residue was dried under vacuum.

*Preparation of benzyl $N^2$-(tert-butoxycarbonyl)-$N^4$-(tert-butyl)-L-asparaginate* ***1***

The title compound was synthesized by following the general procedure for HATU mediated coupling of Boc-Asp-OBn (3.55g, 11 mmol) and *tert*-butylamine (0.73 g, 10 mmol). The isolated off-white product (2.95g, 78%) was used in next step without further purification. $^1$H NMR (500 MHz, $CDCl_3$) δ 7.36 – 7.26 (m, 5H), 5.92 – 5.76 (m, 1H), 5.41 (br, 1H), 5.21 (d, *J* = 12.4 Hz, 1H), 5.14 (d, *J* = 12.4 Hz, 1H), 4.57 – 4.42 (m, 1H), 2.79 (dd, *J* = 15.8, 4.9 Hz, 1H), 2.62 (dd, *J* = 15.8, 4.2 Hz, 1H), 1.42 (s, 9H), 1.29 (s, 9H). $ES^+$ calc. for $C_{20}H_{31}N_2O_5$ $[M + H]^+$: 379.2. Found: 379.2.

*Preparation of $N^2$-(tert-butoxycarbonyl)-$N^4$-(tert-butyl)-L-asparagine* ***2***

**1** (1.89 g, 5 mmol) was dissolved in methanol, Palladium on carbon (10%) was added carefully. Residual air from the flask was removed and flask was flushed with hydrogen. The mixture was stirred at room temperature for 3 h under hydrogen atmosphere using a hydrogen balloon. After completion of reaction, the mixture was filtered through celite. Filtrate was evaporated and dried under vacuum to give **2** as a white powder (1.45g, quant.). $ES^+$ calc. for $C_{13}H_{23}N_2O_5$ $[M – H]^–$: 287.2. Found: 287.2.

*Preparation of N-(2-aminoethyl)-2',4-difluoro-[1,1'-biphenyl]-3-carboxamide trifluoroacetate salt* ***3***

The title compound was synthesized by two successive steps, one following the general procedure for HATU mediated coupling of 2-fluoro-5-(2-fluorophenyl)benzoic acid (141.6 mg, 605 μmol) and *tert*-butyl *N*-(2-aminoethyl)carbamate (88.1 mg, 550 μmol) and the other following the general procedure for Boc-deprotection of the product in first step. The isolated white product (172.0 mg, two step yield: 80%) was used in next step without further purification. $^1$H NMR (500 MHz, DMSO-*d6*) δ 8.61 – 8.50 (m, 1H), 8.04 – 7.76 (m, 4H), 7.76 – 7.67 (m, 1H), 7.61 – 7.51 (m, 1H), 7.49 – 7.39 (m, 2H), 7.38 – 7.30 (m, 2H), 3.59 – 3.46 (m, 2H), 3.07 – 2.93 (m, 2H). $ES^+$ calc. for $C_{15}H_{15}F_2N_2O$ $[M + H]^+$: 277.2. Found 277.2.

*Preparation of tert-butyl (S)-(4-(tert-butylamino)-1-((2-(2',4-difluoro-[1,1'-biphenyl]-3-carboxamido)ethyl)amino)-1,4-dioxobutan-2-yl)carbamate* ***5***

The title compound was synthesized by following the general procedure for HATU mediated coupling of **2** (63.5 mg, 220 μmol) and **3** (78.0 mg, 200 μmol). After completion of reaction, mixture was purified by flash column chromatography to give product (95.0 mg. 87%) as a white solid. $^1$H NMR (500 MHz, DMSO-*d6*) δ 8.44 – 8.33 (m, 1H), 7.98 (t, *J* = 5.6 Hz, 1H), 7.83 – 7.75 (m, 1H), 7.74 – 7.67 (m, 1H), 7.60 – 7.52 (m, 1H), 7.48 – 7.42 (m, 1H), 7.42 – 7.29 (m, 4H), 6.74 (d, *J* = 8.2 Hz, 1H), 4.24 – 4.13 (m, 1H), 3.40 – 3.24 (m, 3H), 3.24 – 3.15 (m, 1H), 2.38 (dd, *J* =

14.3, 5.4 Hz, 1H), 2.30 (dd, *J* = 14.3, 8.4 Hz, 1H), 1.34 (s, 9H), 1.19 (s, 9H). HRMS calc. for $C_{28}H_{36}F_2N_4O_5Na$ [M + Na]$^+$: 569.2551. Found: 569.2564.

*Preparation of (S)-2-amino-N4-(tert-butyl)-N1-(2-(2',4-difluoro-[1,1'-biphenyl]-3-carboxamido)ethyl)succinamide trifluoroacetate salt* ***7***

The title compound was synthesized by following the general procedure for Boc-deprotection of ***5*** (42.0 mg, 76 μmol). Isolated crude was purified by preparative LCMS to give product (40.2 mg, 90%) as a colorless gum. $^1$H NMR (500 MHz, DMSO-*d6*) δ 8.49 (t, *J* = 5.2 Hz, 1H), 8.46 – 8.41 (m, 1H), 8.10 (d, *J* = 4.8 Hz, 3H), 7.82 – 7.78 (m, 2H), 7.73 – 7.69 (m, 1H), 7.59 – 7.54 (m, 1H), 7.49 – 7.39 (m, 2H), 7.36 – 7.30 (m, 2H), 4.00 – 3.94 (m, 1H), 3.43 – 3.29 (m, 3H), 3.29 – 3.19 (m, 1H), 2.65 (dd, *J* = 16.5, 5.1 Hz, 1H), 2.55 (dd, *J* = 16.5, 7.8 Hz, 1H), 1.22 (s, 9H). ES$^+$ calc. for $C_{23}H_{29}F_2N_4O_3$ [M + H]$^+$: 447.2. Found: 447.3.

*Preparation of (3R,5aS,6R,8aS,9R,10S,12R,12aR)-3,6,9-trimethyldecahydro-12H-3,12-epoxy[1,2]dioxepino[4,3-i]isochromen-10-yl 4-(((S)-4-(tert-butylamino)-1-((2-(2',4-difluoro-[1,1'-biphenyl]-3-carboxamido)ethyl)amino)-1,4-dioxobutan-2-yl)amino)-4-oxobutanoate* ***WZ-13***

The title compound was synthesized by following the general procedure for HATU mediated coupling of artesunate (21.0 mg, 55 μmol) and **7** (28.0 mg, 50 μmol). After completion of reaction, mixture was purified by preparative LCMS to give product (30.5 mg, 75%) as a white powder. $^1$H NMR (500 MHz, $CDCl_3$) δ 8.14 (d, *J* = 6.7 Hz, 1H), 7.65 (t, *J* = 7.1 Hz, 2H), 7.55 (t, *J* = 5.0 Hz, 1H), 7.42 (t, *J* = 7.6 Hz, 2H), 7.32 (dd, *J* = 13.3, 6.8 Hz, 1H), 7.23 – 7.10 (m, 3H), 5.86 (s, 1H), 5.62 (d, *J* = 9.8 Hz, 1H), 5.39 (d, *J* = 11.1 Hz, 1H), 4.72 – 4.65 (m, 1H), 3.71 – 3.40 (m, 4H), 3.01 – 2.58 (m, 2H), 2.54 – 2.27 (m, 4H), 2.06 – 1.70 (m, 3H), 1.65 – 1.48 (m, 3H), 1.45 – 1.13 (m, 16H), 0.98 – 0.86 (m, 2H), 0.83 (d, *J* = 6.2 Hz, 3H), 0.74 (d, *J* = 7.1 Hz, 3H). $^{13}$C NMR (125 MHz, $CDCl_3$) δ 173.04, 171.59, 171.55, 170.96, 163.77, 163.75, 160.72, 159.15, 158.75, 133.72, 133.67, 132.54, 132.52, 132.21, 130.73, 130.71, 129.71, 129.64, 127.37, 127.26, 124.66, 124.63, 122.00, 121.90, 116.41, 116.36, 116.22, 116.19, 104.52, 104.19, 103.48, 92.80, 91.52, 87.89, 80.23, 51.67, 51.66, 50.59, 45.26, 39.66, 39.51, 37.61, 37.12, 36.33, 34.16, 31.72, 31.18, 29.81, 28.64, 25.92, 24.58, 21.88, 20.19, 12.09. HRMS calc. for $C_{42}H_{55}F_2N_4O_{10}$ [M + H]$^+$: 813.3881. Found: 813.3907.

*Preparation of (3R,5aS,6R,8aS,9R,10S,12R,12aR)-3,6,9-trimethyldecahydro-12H-3,12-epoxy[1,2]dioxepino[4,3-i]isochromen-10-yl 4-(((S)-4-(tert-butylamino)-1,4-dioxo-1-((2-(4-phenylpicolinamido)ethyl)amino)butan-2-yl)amino)-4-oxobutanoate **WZ-06***

The title compound was synthesized by following the general procedure for HATU mediated coupling of artesunate (21.0 mg, 55 μmol) and **8** (26.3 mg, 50 μmol). After completion of reaction, mixture was purified by preparative LCMS to give product (31.5 mg, 81%) as a white powder. $^{1}$H NMR (500 MHz, $CDCl_3$) δ 8.81 (d, *J* = 5.5 Hz, 1H), 8.72 (t, *J* = 5.7 Hz, 1H), 8.59 (s, 1H), 7.99 (d, *J* = 7.7 Hz, 1H), 7.94 (d, *J* = 5.3 Hz, 1H), 7.84 – 7.75 (m, 3H), 7.56 (dd, *J* = 9.9, 5.2 Hz, 3H), 6.75 (s, 1H), 5.65 (d, *J* = 9.9 Hz, 1H), 5.43 (s, 1H), 4.72 (dd, *J* = 12.2, 6.6 Hz, 1H), 3.83 (dd, *J* = 12.8, 5.7 Hz, 1H), 3.64 – 3.47 (m, 2H), 3.39 (dd, *J* = 12.5, 7.1 Hz, 1H), 2.80 (dd, *J* = 14.7, 6.6 Hz, 1H), 2.73 – 2.52 (m, 4H), 2.50 – 2.29 (m, 3H), 1.99 (d, *J* = 14.4 Hz, 1H), 1.86 (dd, *J* = 8.9, 4.5 Hz, 1H), 1.75 – 1.54 (m, 3H), 1.48 – 1.22 (m, 16H), 1.02 – 0.86 (m, 4H), 0.76 (d, *J* = 7.1 Hz, 3H). $^{13}$C NMR (125 MHz, $CDCl_3$) δ 173.08, 172.70, 172.15, 171.26, 162.84, 154.12, 147.34, 146.24, 135.98, 131.00, 129.71, 127.75, 124.82, 121.35, 104.83, 93.12, 91.81, 80.32, 52.22, 51.64, 51.31, 45.22, 39.70, 38.97, 37.74, 37.28, 36.31, 34.13, 31.69, 30.51, 29.58, 28.47, 25.80, 24.58, 21.95, 20.26, 12.07. HRMS calc. for $C_{41}H_{56}N_5O_{10}$ $[M + H]^+$: 778.4022. Found: 778.4001.

*Preparation of (S)-4-((4-(tert-butylamino)-1-((2-(2',4-difluoro-[1,1'-biphenyl]-3-carboxamido)ethyl)amino)-1,4-dioxobutan-2-yl)amino)-4-oxobutanoic acid **WZ-20***

To a stirred solution of **7** (28.0 mg, 50 μmol) and succinic anhydride (5.5 mg, 55 μmol) in dry DMF (1 mL) was added Hünig base (35μL, 200 μmol) at 0 °C. The reaction mixture was allowed to stir at r.t. for 4 h. After completion of reaction, mixture was purified by preparative LCMS to give product (24.5 mg, 90%) as a white solid. $^{1}$H NMR (500 MHz, $CD_3OD$) δ 8.11 – 7.78 (m, 1H), 7.70 (s, 1H), 7.55 – 7.10 (m, 6H), 4.66 – 4.54 (m, 1H), 3.62 – 3.34 (m, 4H), 2.69 – 2.32 (m, 5H), 1.27 (s, 9H). $^{13}$C NMR (125 MHz, $CD_3OD$) δ 174.73, 173.82, 171.64, 171.56, 166.85, 161.99, 161.92, 160.03, 159.92, 134.55, 133.67, 132.03, 131.82, 130.95, 130.89, 128.48, 128.38, 125.91, 125.88, 124.37, 124.26, 117.52, 117.34, 117.21, 117.03, 52.13, 52.09, 52.02, 40.51, 40.28, 39.08, 39.03, 28.88. HRMS calc. for $C_{27}H_{33}F_2N_4O_6$ $[M + H]^+$: 547.2363. Found: 547.2355.

*Preparation of 2-((3R,5aS,6R,8aS,9R,10R,12R,12aR)-3,6,9-trimethyldecahydro-12H-3,12-epoxy[1,2]dioxepino[4,3-i]isochromen-10-yl)acetic acid **9***

Using artemisinin as the starting material, ART-AcOH **9** was synthesized in four steps according to the literature procedures,(4) $^{1}H$ NMR (500 MHz, $CDCl_3$) δ 5.34 (s, 1H), 4.83 (ddd, $J$ = 9.9, 6.0, 3.7 Hz, 1H), 2.75 – 2.62 (m, 2H), 2.49 (dd, $J$ = 15.6, 3.5 Hz, 1H), 2.31 (td, $J$ = 14.0, 3.7 Hz, 1H), 2.06 – 1.88 (m, 3H), 1.81 – 1.75 (m, 1H), 1.70 – 1.62 (m, 2H), 1.40 (s, 3H), 1.31 – 1.21 (m, 4H), 0.95 (d, $J$ = 5.9 Hz, 3H), 0.86 (d, $J$ = 7.6 Hz, 3H). $^{13}C$ NMR (125 MHz, $CDCl_3$) δ 176.45, 103.39, 89.43, 80.95, 71.15, 52.22, 44.03, 37.55, 36.56, 35.94, 34.47, 29.84, 25.93, 24.82, 24.77, 20.21, 12.89. $ES^+$ calc. for $C_{17}H_{26}O_4$ $[M - O_2]^+$: 294.

*Preparation of 2-((3R,5aS,6R,8aS,9R,10R,12R,12aR)-3,6,9-trimethyldecahydro-12H-3,12-epoxy[1,2]dioxepino[4,3-i]isochromen-10-yl)acetamide* ***ART1***

The title compound was synthesized by following the general procedure for HATU mediated coupling of ***9*** (32.6 mg, 100 μmol) and 2 M Ammonia solution in ethanol (1 mL, 2 mmol). After completion of reaction, mixture was purified by preparative LCMS to give product (31.0 mg, 95%) as a white solid. $^{1}H$ NMR (500 MHz, $CDCl_3$) δ 6.98 (s, 1H), 5.58 (s, 1H), 5.37 (s, 1H), 4.79 (dd, $J$ = 11.0, 6.1 Hz, 1H), 2.63 – 2.49 (m, 2H), 2.31 (dt, $J$ = 14.0, 7.3 Hz, 2H), 2.04 (d, $J$ = 14.4 Hz, 2H), 1.96 (dd, $J$ = 10.1, 4.4 Hz, 1H), 1.83 – 1.62 (m, 3H), 1.37 (s, 3H), 1.31 – 1.19 (m, 3H), 1.02 – 0.90 (m, 4H), 0.87 (d, $J$ = 7.5 Hz, 3H). $^{13}C$ NMR (125 MHz, $CDCl_3$) δ 174.65, 103.06, 90.28, 81.00, 69.79, 51.91, 43.47, 37.64, 37.31, 36.60, 34.34, 30.52, 25.84, 24.93, 24.92, 20.11, 12.11. HRMS calc. for $C_{17}H_{27}NNaO_5$ $[M + Na]^+$: 348.1781. Found: 348.1770.

*Preparation of* tert-butyl (S)-(2-((4-(tert-butylamino)-1-((2-(2',4-difluoro-[1,1'-biphenyl]-3-carboxamido)ethyl)amino)-1,4-dioxobutan-2-yl)amino)-2-oxoethyl)carbamate ***WZ-0917***

The title compound was synthesized by following the general procedure for HATU mediated coupling of ***Boc-Gly-OH*** (19.3 mg, 110 μmol) and **7** (56.0 mg, 100 μmol). After completion of reaction, mixture was purified by preparative LCMS to give product (54.2 mg, 90%) as a white solid. $^{1}H$ NMR (500 MHz, $CDCl_3$) δ 8.22 – 7.98 (m, 2H), 7.76 (s, 1H), 7.63 – 7.47 (m, 2H), 7.40 (dd, $J$ = 11.1, 4.2 Hz, 1H), 7.33 – 7.27 (m, 1H), 7.20 – 7.08 (m, 3H), 6.30 (s, 1H), 5.74 (s, 1H),

4.69 (d, $J$ = 6.4 Hz, 1H), 3.74 (qd, $J$ = 17.0, 5.5 Hz, 2H), 3.58 (s, 2H), 3.51 – 3.32 (m, 2H), 2.78 (dd, $J$ = 14.6, 3.7 Hz, 1H), 2.44 (dd, $J$ = 14.7, 4.8 Hz, 1H), 1.36 (s, 9H), 1.20 (s, 9H). $^{13}$C NMR (125 MHz, $CDCl_3$) δ 174.09, 171.76, 170.41, 170.05, 164.30, 163.23, 160.98, 160.64, 158.99, 158.66, 156.75, 133.75, 133.70, 132.49, 132.47, 131.98, 130.69, 130.67, 129.62, 129.56, 127.24, 127.14, 124.62, 124.59, 121.80, 121.70, 116.36, 116.28, 116.16, 116.10, 80.53, 51.50, 50.62, 44.69, 39.98, 39.91, 37.60, 28.52, 28.30. ES$^+$ calc. for $C_{30}H_{40}F_2N_5O_6$ [M + H]$^+$: 604.3. Found: 604.3.

*Preparation of tert-butyl (S)-(3-((4-(tert-butylamino)-1-((2-(2',4-difluoro-[1,1'-biphenyl]-3-carboxamido)ethyl)amino)-1,4-dioxobutan-2-yl)amino)-3-oxopropyl)carbamate* ***WZ-0918***

The title compound was synthesized by following the general procedure for HATU mediated coupling of ***Boc-β-Ala-OH*** (20.8 mg, 110 μmol) and **7** (56.0 mg, 100 μmol). After completion of reaction, mixture was purified by preparative LCMS to give product (50.7 mg, 82%) as a white solid. $^1$H NMR (500 MHz, $CDCl_3$) δ 8.12 (d, $J$ = 6.4 Hz, 1H), 7.78 (s, 1H), 7.61 (dd, $J$ = 4.4, 1.6 Hz, 2H), 7.42 (t, $J$ = 7.4 Hz, 2H), 7.31 (dd, $J$ = 12.8, 6.7 Hz, 1H), 7.22 – 7.11 (m, 3H), 6.24 (s, 1H), 5.40 (s, 1H), 4.70 (s, 1H), 3.62 (d, $J$ = 4.3 Hz, 2H), 3.55 – 3.41 (m, 2H), 2.73 (d, $J$ = 13.1 Hz, 1H), 2.56 – 2.37 (m, 4H), 1.37 (s, 9H), 1.24 (s, 9H). ES$^+$ calc. for $C_{31}H_{42}F_2N_5O_6$ [M + H]$^+$: 618.3. Found: 618.3.

*Preparation of tert-butyl (S)-(4-((4-(tert-butylamino)-1-((2-(2',4-difluoro-[1,1'-biphenyl]-3-carboxamido)ethyl)amino)-1,4-dioxobutan-2-yl)amino)-4-oxobutyl)carbamate* ***PI01***

The title compound was synthesized by following the general procedure for HATU mediated coupling of ***Boc-GABA-OH*** (22.5 mg, 110 μmol) and **7** (56.0 mg, 100 μmol). After completion of reaction, mixture was purified by preparative LCMS to give product (56.2 mg, 89%) as a white solid. $^1$H NMR (500 MHz, $CDCl_3$) δ 8.14 (d, $J$ = 6.2 Hz, 1H), 8.02 (s, 1H), 7.67 – 7.61 (m, 1H),

7.54 (d, *J* = 6.1 Hz, 1H), 7.44 (t, *J* = 7.3 Hz, 2H), 7.33 (dd, *J* = 13.6, 6.8 Hz, 1H), 7.18 (ddd, *J* = 22.0, 17.3, 9.2 Hz, 3H), 6.01 (s, 1H), 4.71 (s, 2H), 3.68 – 3.41 (m, 4H), 3.15 (d, *J* = 4.8 Hz, 1H), 2.98 (s, 1H), 2.79 (d, *J* = 13.2 Hz, 1H), 2.46 (d, *J* = 10.0 Hz, 1H), 2.29 (t, *J* = 6.3 Hz, 2H), 1.89 (s, 1H), 1.63 (s, 1H), 1.41 (s, 9H), 1.25 (s, 9H). $^{13}$C NMR (125 MHz, $CDCl_3$) δ 172.69, 172.17, 170.89, 164.02, 162.80, 161.10, 160.75, 159.12, 158.78, 156.78, 133.80, 133.76, 132.64, 132.61, 132.28, 130.80, 130.78, 129.70, 129.64, 127.40, 127.29, 124.69, 124.66, 121.88, 121.78, 116.38, 116.20, 79.67, 51.92, 50.77, 40.55, 39.84, 39.04, 38.14, 32.66, 28.60, 28.52, 26.10. HRMS calc. for $C_{32}H_{43}F_2N_5NaO_6$ [M + Na]$^+$: 654.3074. Found: 654.3058.

*Preparation of (S)-N4-(tert-butyl)-N1-(2-(2',4-difluoro-[1,1'-biphenyl]-3-carboxamido)ethyl)-2-(2-((3R,5aS,6R,8aS,9R,10R,12R,12aR)-3,6,9-trimethyldecahydro-12H-3,12-epoxy[1,2]dioxepino[4,3-i]isochromen-10-yl)acetamido)succinamide* ***ATZ1***

The title compound was synthesized by following the general procedure for HATU mediated coupling of ***9*** (18.0 mg, 55 μmol) and **7** (28.0 mg, 50 μmol). After completion of reaction, mixture was purified by preparative LCMS to give product (29.0 mg, 77%) as a white powder. $^1$H NMR (500 MHz, $CDCl_3$) δ 8.24 (dd, *J* = 8.3, 6.7 Hz, 1H), 7.73 – 7.59 (m, 2H), 7.53 (d, *J* = 8.7 Hz, 1H), 7.47 (dd, *J* = 15.8, 7.8 Hz, 2H), 7.34 (dd, *J* = 12.8, 6.2 Hz, 1H), 7.24 – 7.13 (m, 3H), 5.65 (s, 1H), 5.15 (s, 1H), 4.86 – 4.78 (m, 2H), 3.73 (dd, *J* = 18.2, 13.5 Hz, 1H), 3.62 – 3.55 (m, 1H), 3.52 – 3.37 (m, 2H), 3.00 (dd, *J* = 15.4, 4.4 Hz, 1H), 2.51 – 2.42 (m, 2H), 2.41 – 2.28 (m, 2H), 2.22 (td, *J* = 14.1, 3.7 Hz, 1H), 1.96 (d, *J* = 14.3 Hz, 1H), 1.87 – 1.80 (m, 1H), 1.62 – 1.55 (m, 1H), 1.54 – 1.47 (m, 1H), 1.38 – 1.19 (m, 14H), 1.12 (td, *J* = 11.2, 6.2 Hz, 1H), 1.06 – 0.96 (m, 1H), 0.80 (d, *J* = 6.2 Hz, 3H), 0.78 – 0.62 (m, 5H). $^{13}$C NMR (125 MHz, $CDCl_3$) δ 172.04, 171.77, 171.29, 170.11, 163.44, 163.42, 162.52, 161.37, 160.79, 159.37, 158.82, 133.74, 133.70, 133.67, 133.63,

132.56, 132.21, 130.73, 130.71, 129.75, 129.68, 127.25, 127.15, 124.72, 124.69, 122.12, 122.02, 116.48, 116.30, 116.26, 103.00, 90.05, 80.72, 71.69, 51.83, 51.81, 50.13, 43.47, 39.77, 39.62, 39.16, 37.91, 37.21, 36.52, 34.23, 30.24, 28.73, 25.59, 24.74, 24.47, 20.02, 12.46. HRMS calc. for $C_{40}H_{52}F_2N_4NaO_8$ $[M + Na]^+$: 777.3645. Found: 777.3657.

*Preparation of (S)-N4-(tert-butyl)-N1-(2-(2',4-difluoro-[1,1'-biphenyl]-3-carboxamido)ethyl)-2-(2-(2-((3R,5aS,6R,8aS,9R,10R,12R,12aR)-3,6,9-trimethyldecahydro-12H-3,12-epoxy[1,2]dioxepino[4,3-i]isochromen-10-yl)acetamido)acetamido)succinamide* ***ATZ2***

The title compound was synthesized by two successive steps, one following the general procedure for Boc-deprotection of **WZ-0917** (30.2 mg, 50 μmol) and the other following the general procedure for HATU mediated coupling of ***9*** (18.0 mg, 55 μmol) with all of the product in first step. After completion of reaction, mixture in the second step was purified by preparative LCMS to give product (33.8 mg, two step yield: 83%) as a white powder. $^1$H NMR (500 MHz, $CDCl_3$) δ 8.26 (d, $J$ = 7.8 Hz, 1H), 8.11 (d, $J$ = 5.9 Hz, 1H), 7.85 (t, $J$ = 4.7 Hz, 1H), 7.76 (d, $J$ = 6.7 Hz, 1H), 7.65 – 7.56 (m, 2H), 7.44 (t, $J$ = 7.4 Hz, 1H), 7.32 (dd, $J$ = 12.9, 6.4 Hz, 1H), 7.23 – 7.10 (m, 3H), 5.89 (s, 1H), 5.37 (s, 1H), 4.85 (dd, $J$ = 10.8, 6.1 Hz, 1H), 4.64 – 4.57 (m, 1H), 3.81 (d, $J$ = 5.1 Hz, 2H), 3.69 – 3.58 (m, 2H), 3.47 (d, $J$ = 5.1 Hz, 2H), 2.78 (dd, $J$ = 14.7, 3.8 Hz, 1H), 2.60 – 2.50 (m, 2H), 2.39 (dd, $J$ = 14.7, 5.1 Hz, 1H), 2.31 (dd, $J$ = 22.6, 9.5 Hz, 2H), 2.06 – 1.99 (m, 1H), 1.94 (d, $J$ = 11.7 Hz, 1H), 1.79 – 1.71 (m, 1H), 1.71 – 1.60 (m, 2H), 1.35 (s, 3H), 1.32 – 1.11 (m, 13H), 1.04 – 0.87 (m, 4H), 0.80 (d, $J$ = 7.4 Hz, 3H). $^{13}$C NMR (125 MHz, $CDCl_3$) δ 174.11, 171.41, 170.56, 168.98, 164.08, 164.06, 162.91, 160.99, 160.74, 159.00, 158.77, 133.60, 133.57, 133.53, 133.50, 132.48, 132.46, 132.16, 130.81, 130.79, 129.64, 129.57, 127.40, 127.29, 124.67, 124.64, 122.43, 122.33, 116.35, 116.33, 116.17, 116.13, 103.25, 89.94, 80.93, 70.32, 52.03, 51.68, 50.96,

44.36, 43.64, 40.06, 40.04, 37.57, 37.23, 36.76, 36.55, 34.33, 30.28, 28.55, 25.91, 24.90, 24.85, 20.14, 12.27. HRMS calc. for $C_{42}H_{55}F_2N_5NaO_9$ [M + Na]$^+$: 834.3860. Found: 834.3876.

*Preparation of (S)-N4-(tert-butyl)-N1-(2-(2',4-difluoro-[1,1'-biphenyl]-3-carboxamido)ethyl)-2-(3-(2-((3R,5aS,6R,8aS,9R,10R,12R,12aR)-3,6,9-trimethyldecahydro-12H-3,12-epoxy[1,2]dioxepino[4,3-i]isochromen-10-yl)acetamido)propanamido)succinamide* ***ATZ3***

The title compound was synthesized by two successive steps, one following the general procedure for Boc-deprotection of **WZ-0918** (30.9 mg, 50 μmol) and the other following the general procedure for HATU mediated coupling of ***9*** (18.0 mg, 55 μmol) with all of the product in first step. After completion of reaction, mixture in the second step was purified by preparative LCMS to give product (37.5 mg, two step yield: 91%) as a white powder. $^1$H NMR (500 MHz, $CDCl_3$) δ 8.16 (d, *J* = 5.8 Hz, 1H), 7.73 – 7.63 (m, 2H), 7.52 – 7.40 (m, 4H), 7.36 – 7.30 (m, 1H), 7.24 – 7.12 (m, 3H), 5.84 (s, 1H), 5.37 (s, 1H), 4.76 – 4.66 (m, 2H), 3.68 – 3.41 (m, 6H), 2.74 (dd, *J* = 14.7, 3.7 Hz, 1H), 2.60 – 2.40 (m, 5H), 2.29 (dt, *J* = 23.4, 8.7 Hz, 2H), 2.01 (dd, *J* = 11.0, 3.2 Hz, 1H), 1.92 (d, *J* = 12.8 Hz, 1H), 1.76 – 1.72 (m, 1H), 1.65 (dd, *J* = 15.9, 9.6 Hz, 2H), 1.37 (s, 3H), 1.28 – 1.19 (m, 13H), 0.99 – 0.87 (m, 4H), 0.81 (d, *J* = 7.4 Hz, 3H). $^{13}$C NMR (125 MHz, $CDCl_3$) δ 172.51, 172.13, 170.80, 164.13, 161.11, 160.76, 159.13, 158.79, 133.96, 133.92, 133.88, 133.85, 132.74, 132.71, 132.30, 130.81, 130.78, 129.74, 129.67, 127.34, 127.23, 124.72, 124.69, 121.77, 121.67, 116.44, 116.41, 116.23, 103.29, 89.80, 81.07, 71.00, 52.10, 51.72, 50.76, 43.83, 40.13, 40.00, 38.37, 37.57, 37.49, 37.22, 36.60, 36.23, 34.41, 30.35, 28.66, 25.97, 24.89, 24.84, 20.17, 12.50. HRMS calc. for $C_{43}H_{57}F_2N_5NaO_9$ [M + Na]$^+$: 848.4017. Found: 848.4001.

*Preparation of (S)-N4-(tert-butyl)-N1-(2-(2',4-difluoro-[1,1'-biphenyl]-3-carboxamido)ethyl)-2-(4-(2-((3R,5aS,6R,8aS,9R,10R,12R,12aR)-3,6,9-trimethyldecahydro-12H-3,12-epoxy[1,2]dioxepino[4,3-i]isochromen-10-yl)acetamido)butanamido)succinamide* ***ATZ4***

The title compound was synthesized by two successive steps, one following the general procedure for Boc-deprotection of **PI01** (31.5 mg, 50 μmol) and the other following the general procedure for HATU mediated coupling of ***9*** (18.0 mg, 55 μmol) with all of the product in first step. After completion of reaction, mixture in the second step was purified by preparative LCMS to give product (31.5 mg, two step yield: 75%) as a white powder. $^{1}$H NMR (500 MHz, $CDCl_3$) δ 8.13 (d, *J* = 6.3 Hz, 1H), 8.05 (s, 1H), 7.65 (s, 2H), 7.55 – 7.48 (m, 1H), 7.43 (t, *J* = 7.3 Hz, 1H), 7.40 – 7.30 (m, 2H), 7.24 – 7.11 (m, 3H), 6.25 (s, 1H), 5.44 (s, 1H), 4.74 (s, 1H), 4.64 (dd, *J* = 11.0, 6.1 Hz, 1H), 3.63 (d, *J* = 4.4 Hz, 2H), 3.50 (d, *J* = 4.5 Hz, 2H), 3.33 (dt, *J* = 22.8, 11.6 Hz, 1H), 3.09 (dd, *J* = 12.4, 5.8 Hz, 1H), 2.71 (dd, *J* = 14.2, 5.2 Hz, 1H), 2.63 – 2.46 (m, 3H), 2.39 – 2.25 (m, 4H), 2.03 (t, *J* = 12.5 Hz, 1H), 1.94 (d, *J* = 10.9 Hz, 2H), 1.79 – 1.60 (m, 4H), 1.36 (s, 3H), 1.30 – 1.11 (m, 13H), 1.00 – 0.87 (m, 4H), 0.81 (d, *J* = 7.4 Hz, 3H). $^{13}$C NMR (125 MHz, $CDCl_3$) δ 173.75, 173.47, 172.24, 170.85, 164.46, 161.05, 160.73, 159.06, 158.76, 134.02, 133.99, 133.94, 133.91, 132.72, 132.69, 132.14, 130.75, 130.72, 129.78, 129.72, 127.23, 127.13, 124.73, 124.70, 121.65, 121.55, 116.46, 116.40, 116.26, 116.22, 103.50, 89.81, 81.00, 70.87, 52.01, 51.92, 50.95, 43.78, 40.02, 38.38, 37.94, 37.60, 36.89, 36.52, 34.33, 32.23, 30.23, 28.53, 25.86, 25.05, 24.93, 24.80, 20.13, 12.43. HRMS calc. for $C_{44}H_{59}F_2N_5NaO_9$ $[M + Na]^+$: 862.4173. Found: 862.4147.

DeoxyART-AcOH **10** was prepared from ***9*** by an activated zinc mediated reduction according to the literature procedures,[(5)] $^{1}$H NMR (500 MHz, $CDCl_3$) δ 5.26 (s, 1H), 4.61 (q, *J* = 7.3 Hz, 1H), 2.57 – 2.44 (m, 2H), 2.33 (h, *J* = 8.0 Hz, 1H), 1.96 (ddd, *J* = 13.1, 8.4, 4.0 Hz, 1H), 1.85 (dt, *J* =

13.3, 3.8 Hz, 1H), 1.77 (dt, *J* = 13.1, 3.8 Hz, 1H), 1.73 – 1.66 (m, 2H), 1.58 (dq, *J* = 12.8, 5.9 Hz, 2H), 1.25 (q, *J* = 6.0, 4.9 Hz, 3H), 1.23 – 1.18 (m, 1H), 1.18 – 1.12 (m, 2H), 0.99 –0.78 (m, 8H). $^{13}$C NMR (125 MHz, $CDCl_3$) δ 176.68, 107.56, 97.04, 82.65, 65.76, 45.39, 40.34, 37.31, 35.68, 34.62, 34.55, 29.09, 25.16, 23.69, 22.30, 18.87, 12.45. HRMS calc. for $C_{17}H_{26}NaO_5$ [M + Na]$^+$: 333.1672. Found: 333.1681.

*Preparation of (S)-N4-(tert-butyl)-N1-(2-(2',4-difluoro-[1,1'-biphenyl]-3-carboxamido)ethyl)-2-(4-(2-((2R,3R,3aS,3a1R,6R,6aS,9S,10aR)-3,6,9-trimethyldecahydro-10aH-3a1,9-epoxyoxepino [4,3,2-ij]isochromen-2-yl)acetamido)butanamido)succinamide* ***Deoxy-ATZ4***

The title compound was synthesized by two successive steps, one following the general procedure for Boc-deprotection of **PI01** (15.7 mg, 25 μmol) and the other following the general procedure for HATU mediated coupling of ***10*** (9.3 mg, 30 μmol) with all of the product in first step. After completion of reaction, mixture in the second step was purified by preparative LCMS to give product (18.0 mg, two step yield: 89%) as a white powder. $^1$H NMR (500 MHz, $CDCl_3$) δ 8.17 – 8.09 (m, 2H), 7.63 (tq, *J* = 4.5, 3.0, 2.2 Hz, 2H), 7.45 (td, *J* = 7.8, 1.8 Hz, 1H), 7.38 – 7.29 (m, 1H), 7.32 – 7.27 (m, 1H), 7.21 (td, *J* = 7.5, 1.3 Hz, 1H), 7.19 – 7.10 (m, 2H), 6.64 (dd, *J* = 8.6, 4.4 Hz, 1H), 5.77 (s, 1H), 5.27 (s, 1H), 4.80 (dt, *J* = 8.2, 5.1 Hz, 1H), 4.43 (ddd, *J* = 10.4, 7.1, 2.7 Hz, 1H), 3.63 (q, *J* = 5.3 Hz, 2H), 3.57 – 3.46 (m, 3H), 2.89 (dq, *J* = 13.8, 4.5 Hz, 1H), 2.72 (dd, *J* = 14.9, 5.1 Hz, 1H), 2.51 (dd, *J* = 14.9, 5.3 Hz, 1H), 2.28 – 2.10 (m, 5H), 2.08 – 1.82 (m, 3H), 1.81 – 1.66 (m, 2H), 1.64 – 1.51 (m, 1H), 1.25 (s, 10H), 1.23 – 1.10 (m, 4H), 0.96 (td, *J* = 12.4, 11.5, 3.2 Hz, 1H), 0.89 (d, *J* = 5.6 Hz, 3H), 0.76 (d, *J* = 7.6 Hz, 3H). $^{13}$C NMR (126 MHz, $CDCl_3$) δ 172.72, 172.50, 172.02, 170.69, 170.58, 163.94, 163.92, 160.98, 160.77, 158.99, 158.80, 133.52, 133.49, 133.45, 133.41, 132.50, 132.48, 132.16, 132.14, 132.12, 130.84, 130.82, 129.65, 129.59, 127.44, 127.34, 124.68, 124.65, 122.62, 122.51, 116.37, 116.31, 116.19, 116.12, 107.88, 97.25, 82.64, 65.70, 51.57, 50.50, 45.28, 40.31, 40.00, 39.72, 38.78, 38.39, 37.31, 35.73, 34.55, 34.50,

32.28, 29.54, 28.69, 28.66, 26.11, 25.30, 23.77, 22.26, 18.87, 12.03. HRMS calc. for $C_{44}H_{60}F_2N_5O_8$ $[M + H]^+$: 824.4404. Found: 824.4435.

**NMR**

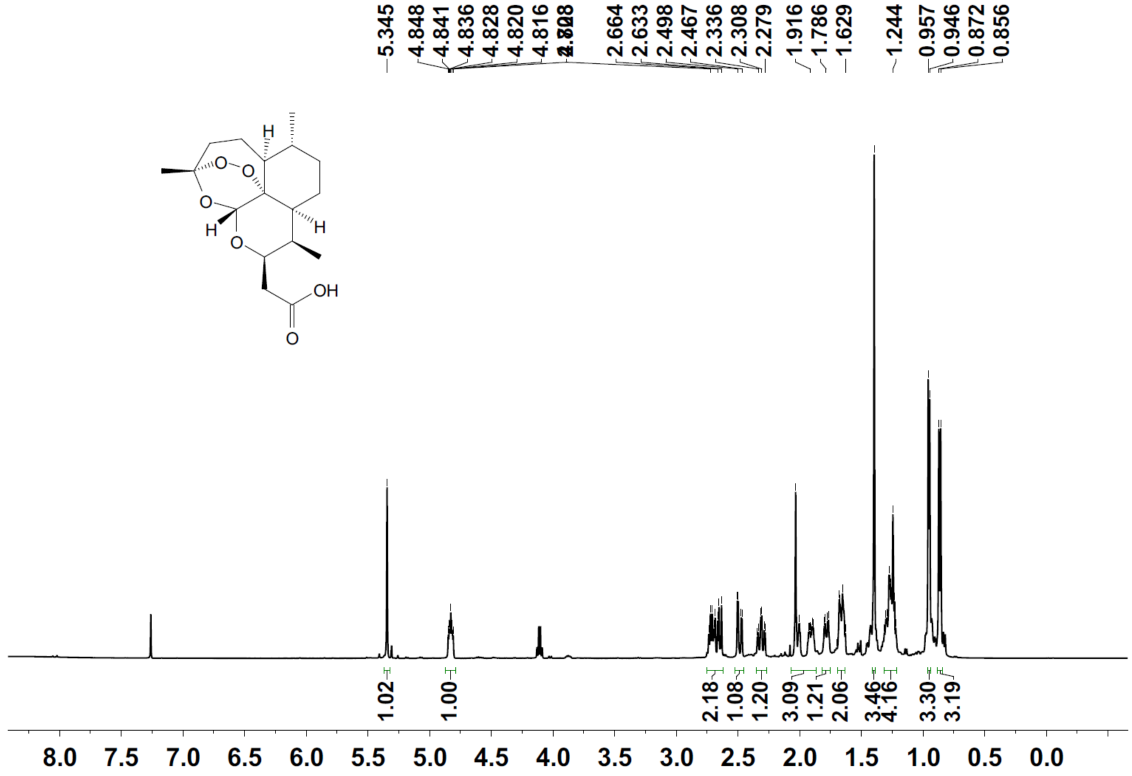
5.345
4.848
4.841
4.836
4.828
4.820
4.816
2.664
2.633
2.498
2.467
2.336
2.308
2.279
1.916
1.786
1.629
1.244
0.957
0.946
0.872
0.856
OH
O
H
1.02
1.00
2.18
1.08
1.20
3.09
1.21
2.06
3.46
4.16
3.30
3.19
8.0 7.5 7.0 6.5 6.0 5.5 5.0 4.5 4.0 3.5 3.0 2.5 2.0 1.5 1.0 0.5 0.0

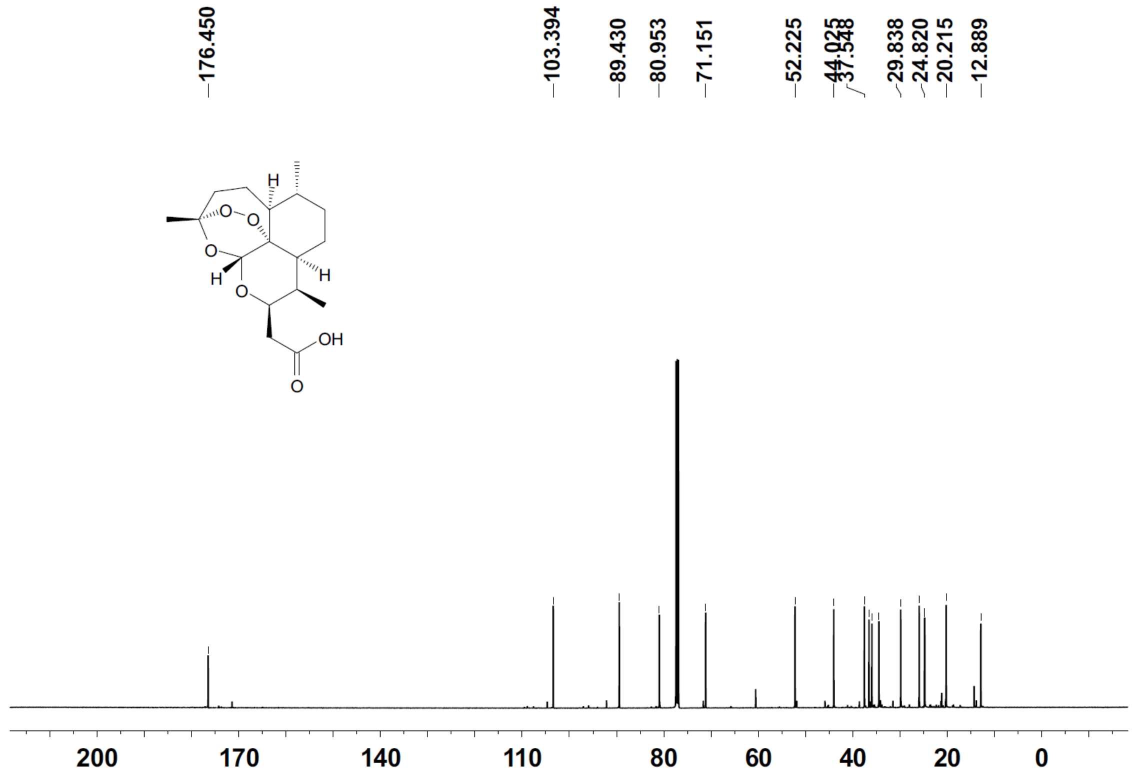
176.450
103.394
89.430
80.953
71.151
52.225
44.025
29.838
24.820
20.215
12.889
OH
O
H
200 170 140 110 80 60 40 20 0

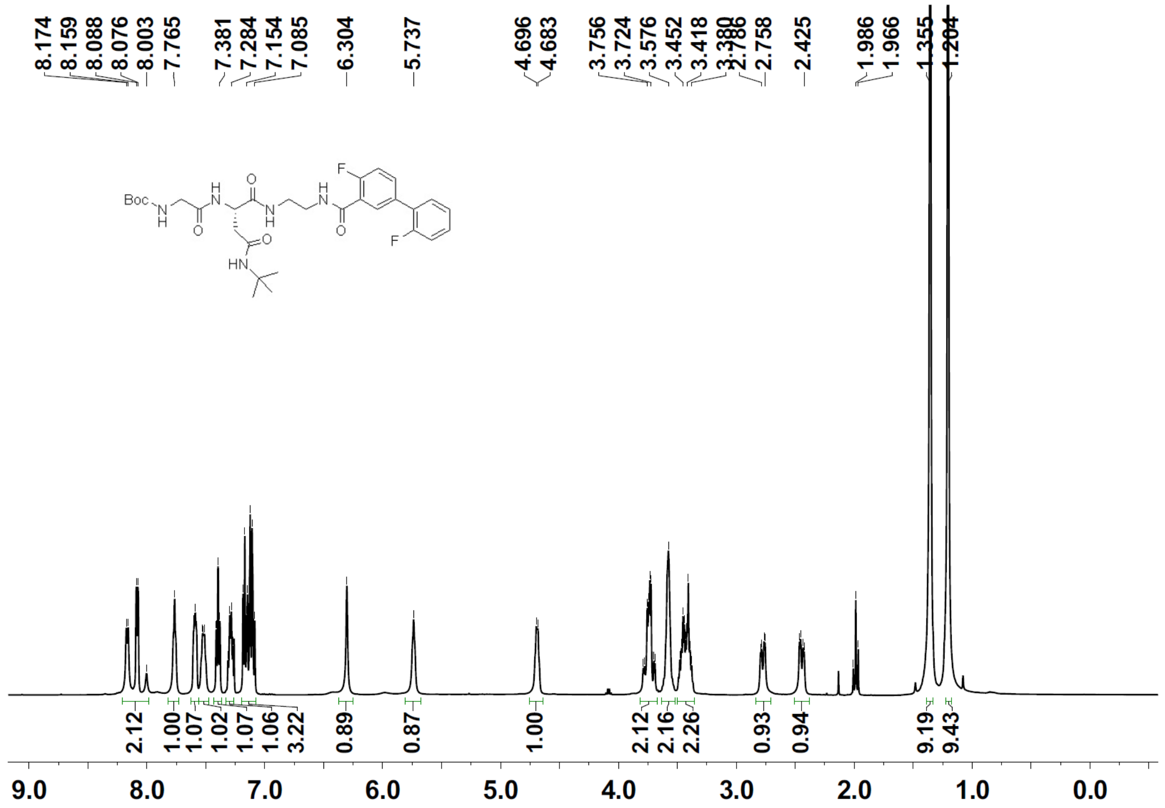

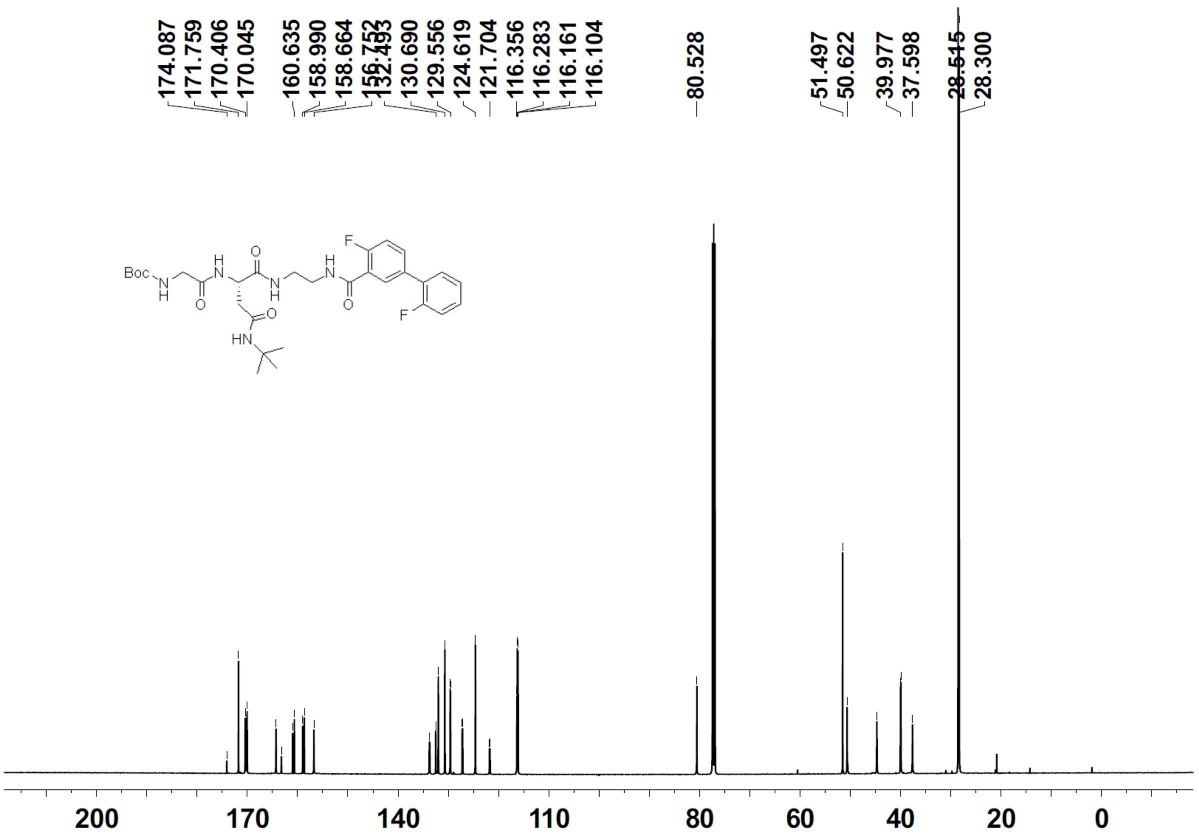

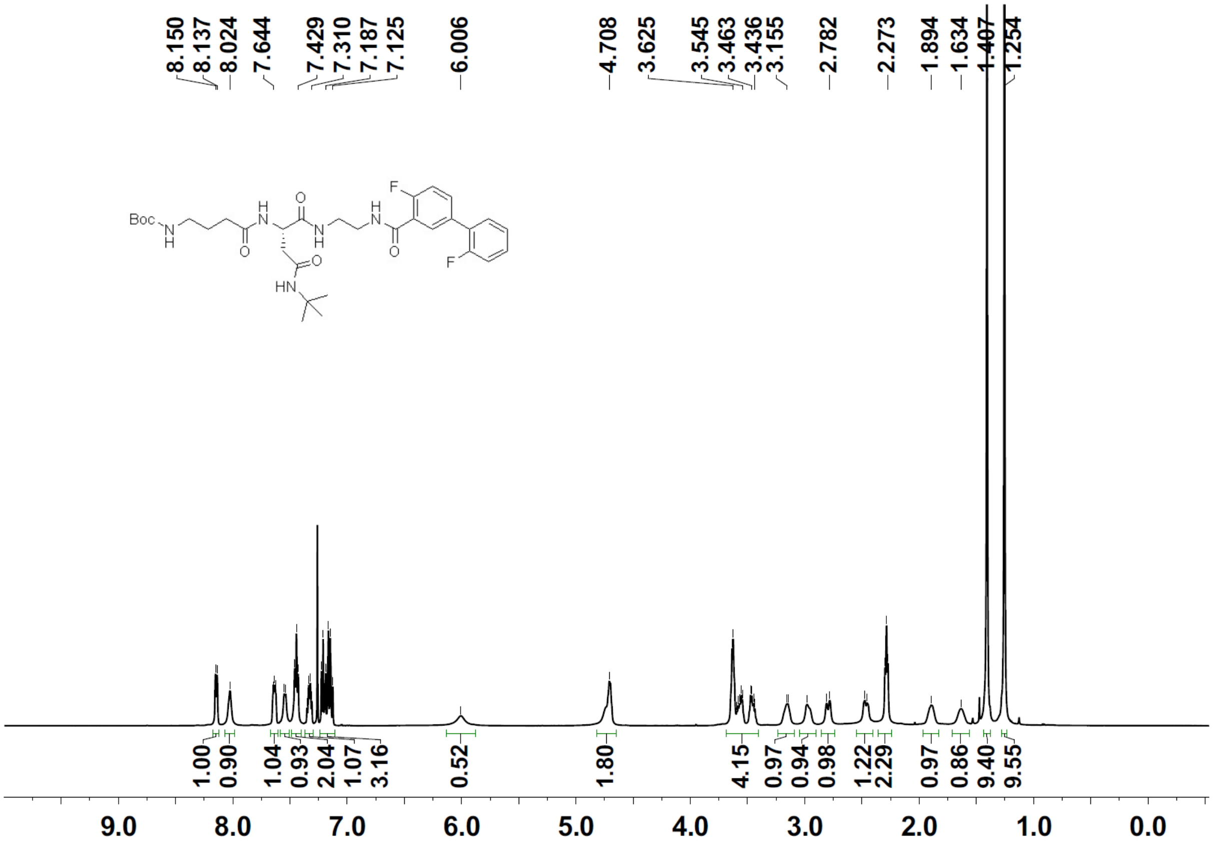
8.150
8.137
8.024
7.644
7.429
7.310
7.187
7.125
6.006
4.708
3.625
3.545
3.463
3.436
3.155
2.782
2.273
1.894
1.634
1.407
1.254
Boc
N
H
O
F
HN
1.00
0.90
1.04
0.93
2.04
1.07
3.16
0.52
1.80
4.15
0.97
0.94
0.98
1.22
2.29
0.97
0.86
9.40
9.55
9.0
8.0
7.0
6.0
5.0
4.0
3.0
2.0
1.0
0.0

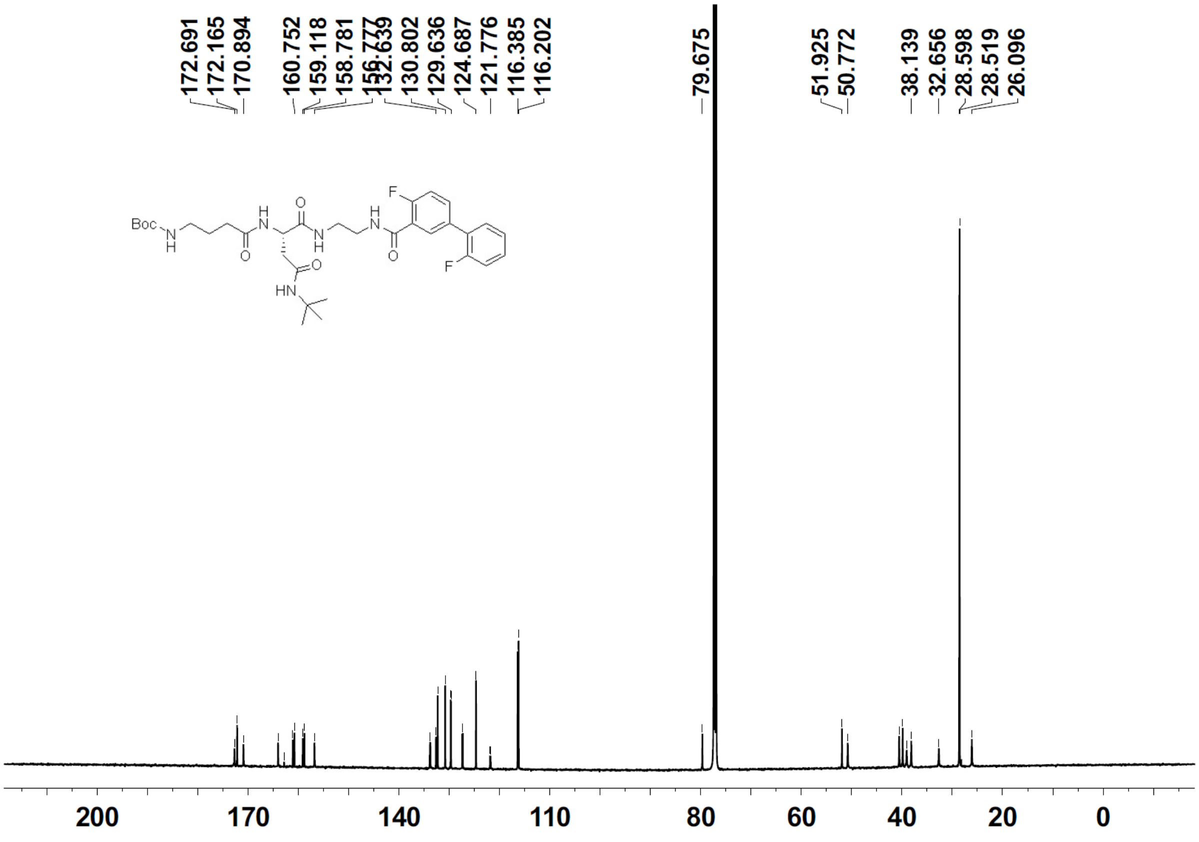
172.691
172.165
170.894
160.752
159.118
158.781
132.639
130.802
129.636
124.687
121.776
116.385
116.202
79.675
51.925
50.772
38.139
32.656
28.598
28.519
26.096
Boc
N
H
O
F
HN
200
170
140
110
80
60
40
20
0

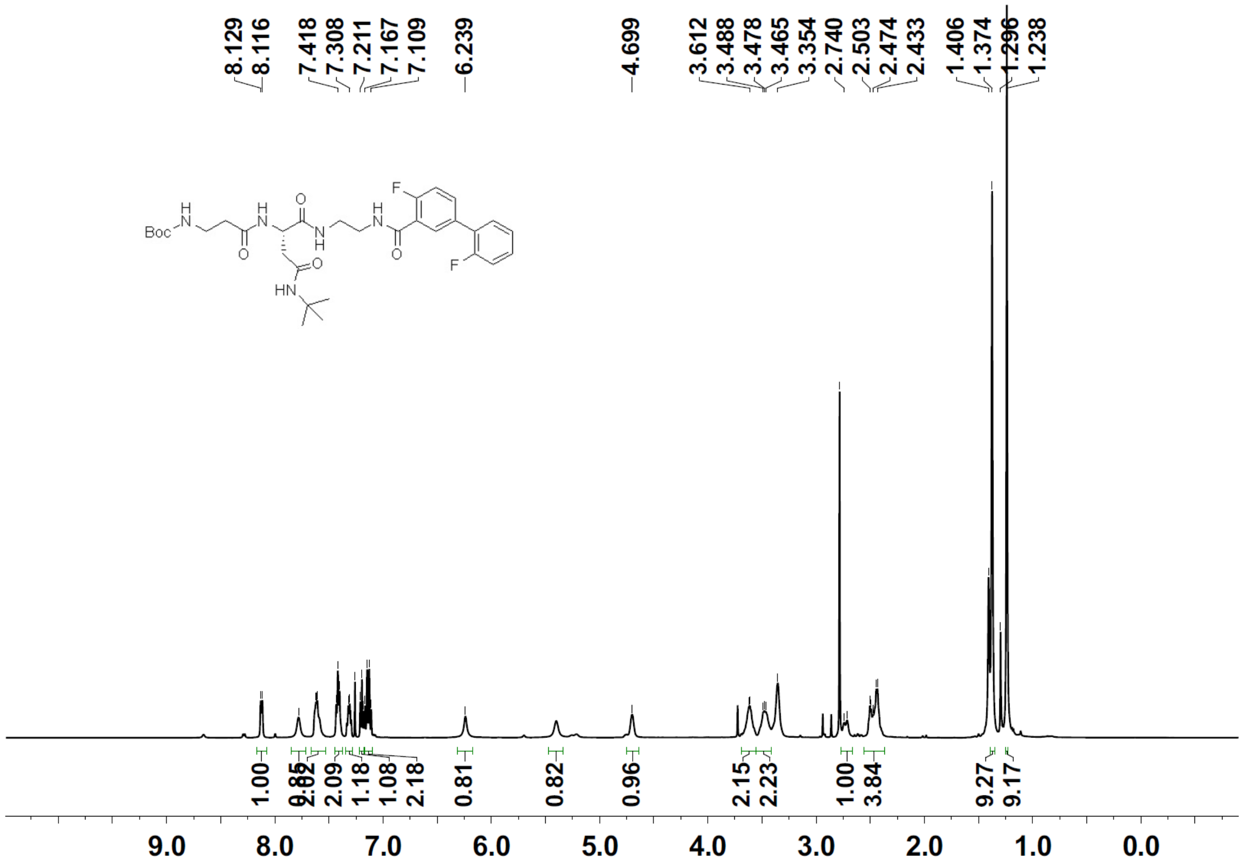
8.129
8.116
7.418
7.308
7.211
7.167
7.109
6.239
4.699
3.612
3.488
3.478
3.465
3.354
2.740
2.503
2.474
2.433
1.406
1.374
1.296
1.238
Boc
F
HN
1.00
0.85
2.02
2.09
1.18
1.08
2.18
0.81
0.82
0.96
2.15
2.23
1.00
3.84
9.27
9.17
9.0
8.0
7.0
6.0
5.0
4.0
3.0
2.0
1.0
0.0

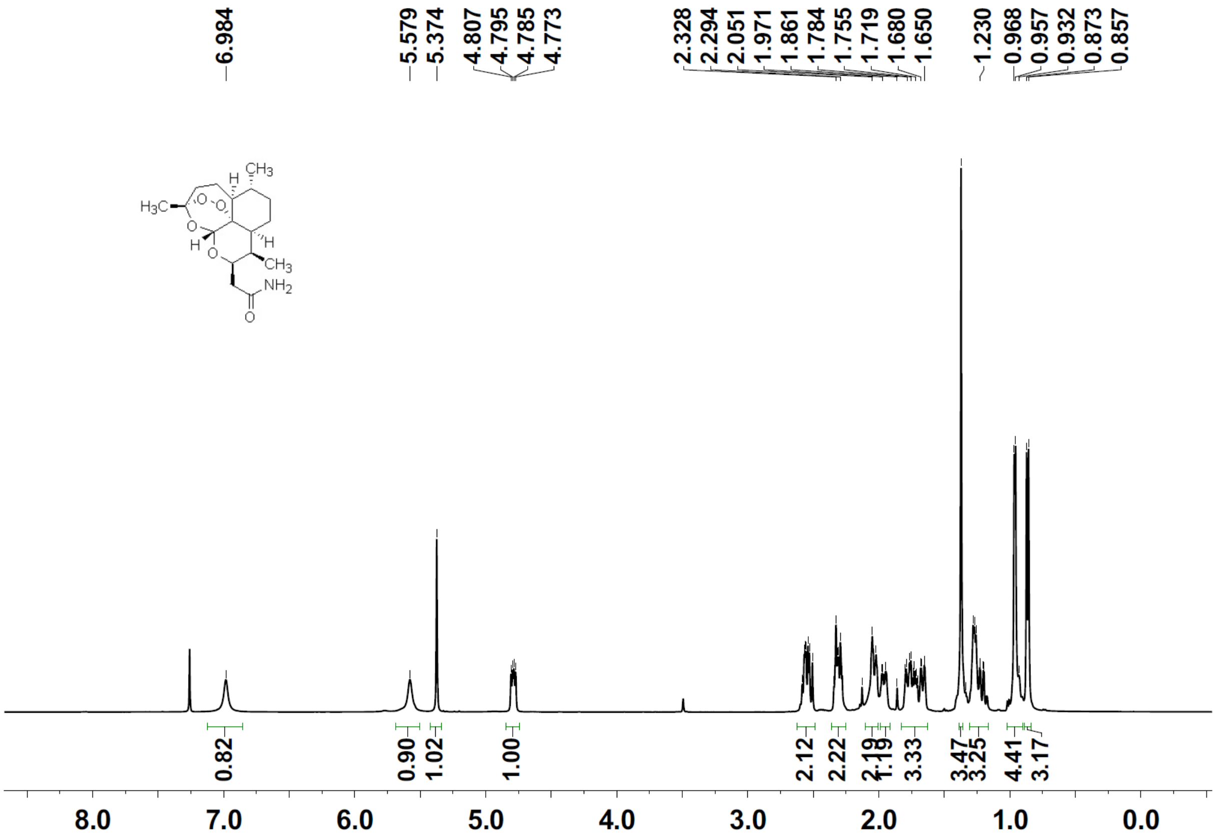
6.984
5.579
5.374
4.807
4.795
4.785
4.773
2.328
2.294
2.051
1.971
1.861
1.784
1.755
1.719
1.680
1.650
1.230
0.968
0.957
0.932
0.873
0.857
0.82
0.90
1.02
1.00
2.12
2.22
2.19
1.19
3.33
3.47
3.25
4.41
3.17
8.0
7.0
6.0
5.0
4.0
3.0
2.0
1.0
0.0

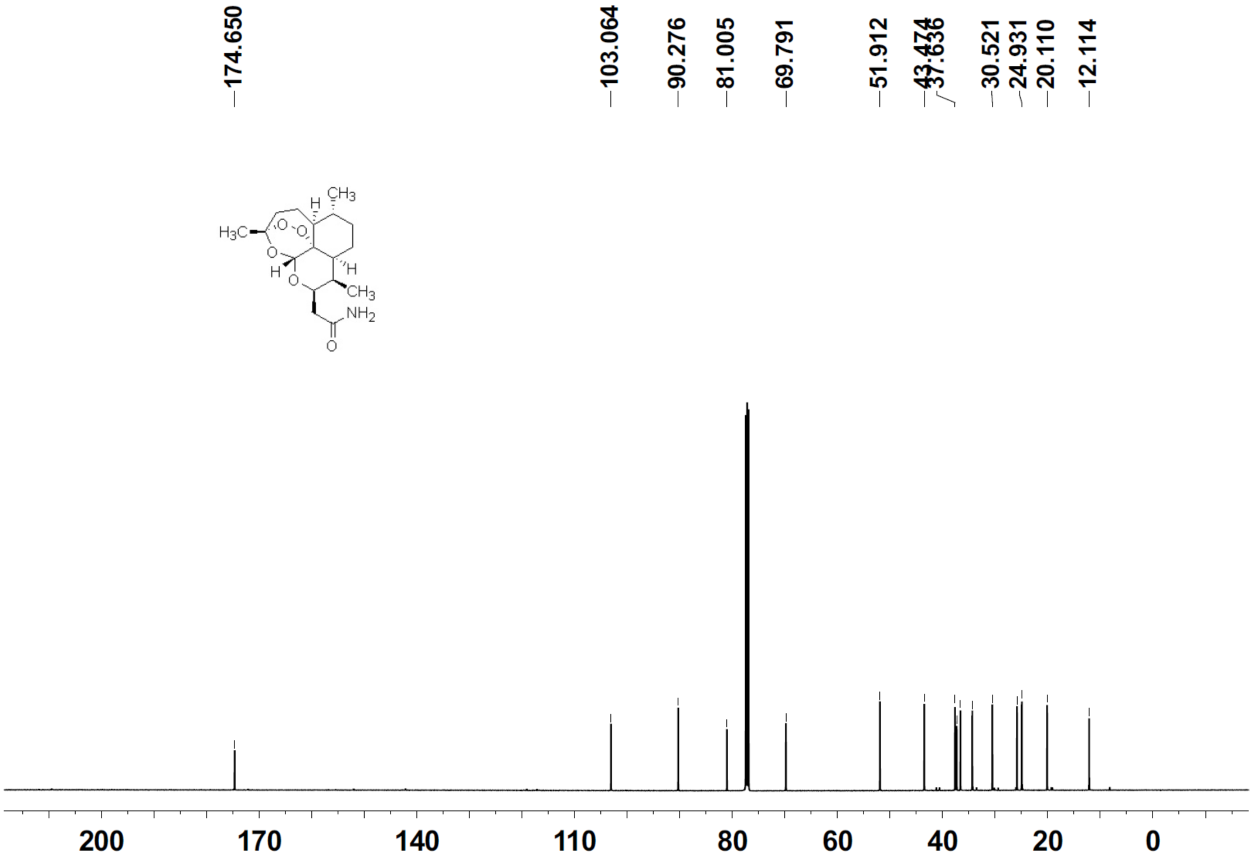
174.650
103.064
90.276
81.005
69.791
51.912
43.474
37.636
30.521
24.931
20.110
12.114
200
170
140
110
80
60
40
20
0

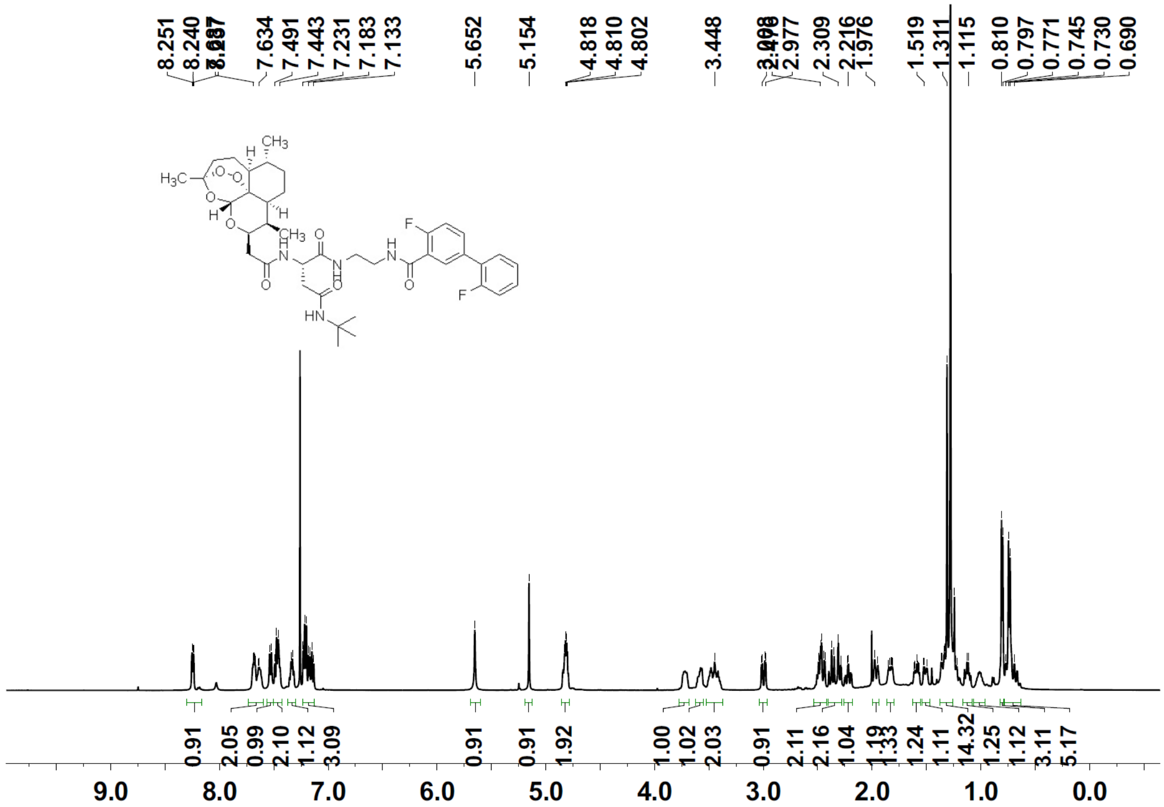
8.251
8.240
7.634
7.491
7.443
7.231
7.183
7.133
5.652
5.154
4.818
4.810
4.802
3.448
2.977
2.309
2.216
1.976
1.519
1.311
1.115
0.810
0.797
0.771
0.745
0.730
0.690
H
CH3
H3C
O
H
F
HN
0.91
2.05
0.99
2.10
1.12
3.09
0.91
0.91
1.92
1.00
1.02
2.03
0.91
2.11
2.16
1.04
1.19
1.33
1.24
1.11
14.32
1.25
1.12
3.11
5.17
9.0
8.0
7.0
6.0
5.0
4.0
3.0
2.0
1.0
0.0

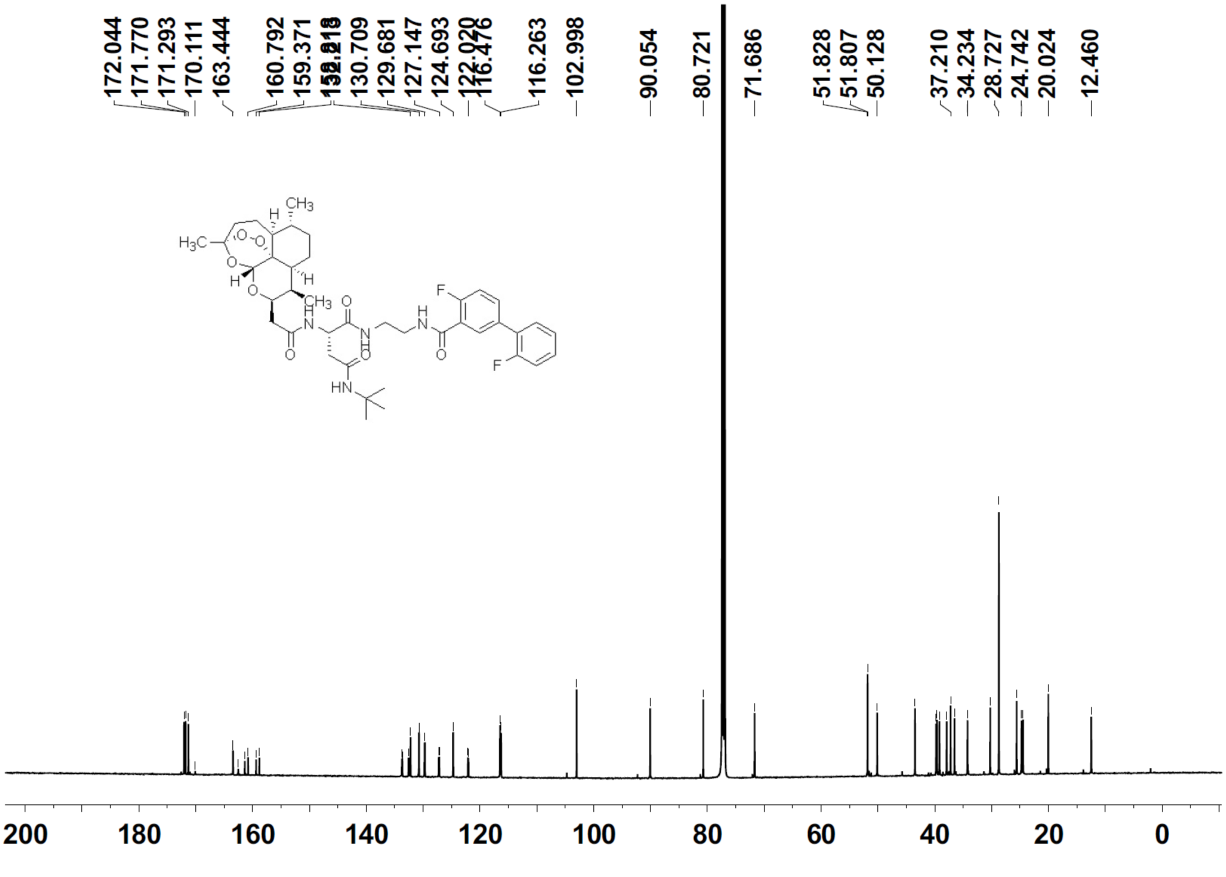
172.044
171.770
171.293
170.111
163.444
160.792
159.371
130.709
129.681
127.147
124.693
122.020
116.476
116.263
102.998
90.054
80.721
71.686
51.828
51.807
50.128
37.210
34.234
28.727
24.742
20.024
12.460
200
180
160
140
120
100
80
60
40
20
0

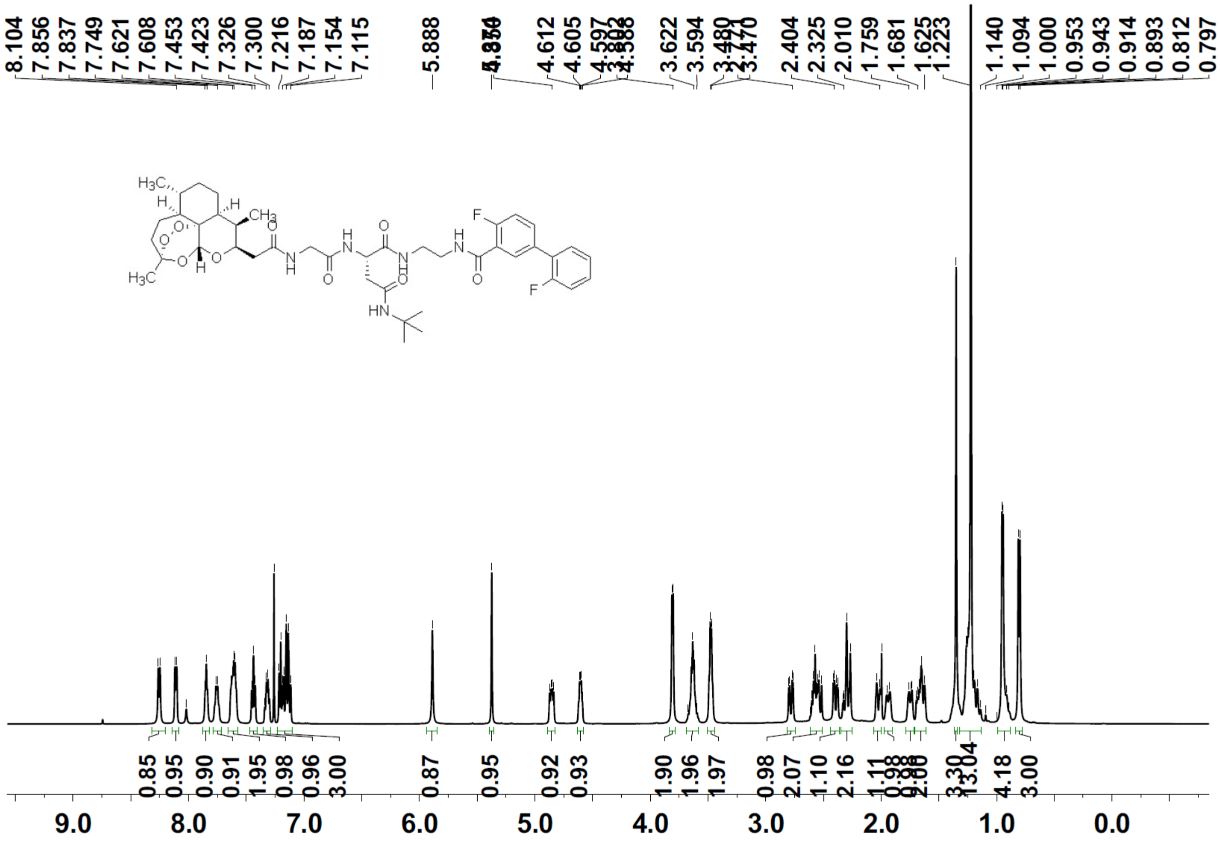

8.104
7.856
7.837
7.749
7.621
7.608
7.453
7.423
7.326
7.300
7.216
7.187
7.154
7.115
5.888
4.612
4.605
4.597
3.622
3.594
3.470
2.404
2.325
2.010
1.759
1.681
1.625
1.223
1.140
1.094
1.000
0.953
0.943
0.914
0.893
0.812
0.797
0.85
0.95
0.90
0.91
1.95
0.98
0.96
3.00
0.87
0.95
0.92
0.93
1.90
1.96
1.97
0.98
2.07
1.10
2.16
1.11
4.18
3.00
9.0
8.0
7.0
6.0
5.0
4.0
3.0
2.0
1.0
0.0

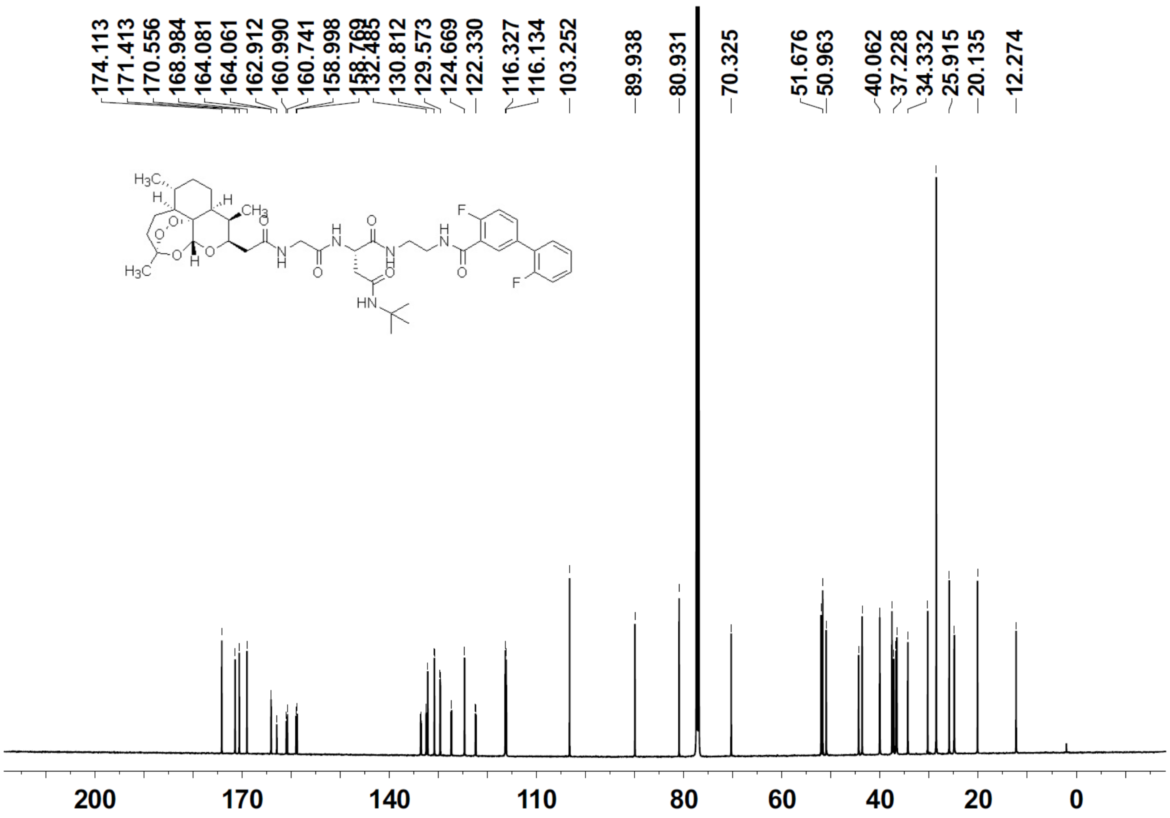

174.113
171.413
170.556
168.984
164.081
164.061
162.912
160.990
160.741
158.998
132.485
130.812
129.573
124.669
122.330
116.327
116.134
103.252
89.938
80.931
70.325
51.676
50.963
40.062
37.228
34.332
25.915
20.135
12.274
200
170
140
110
80
60
40
20
0

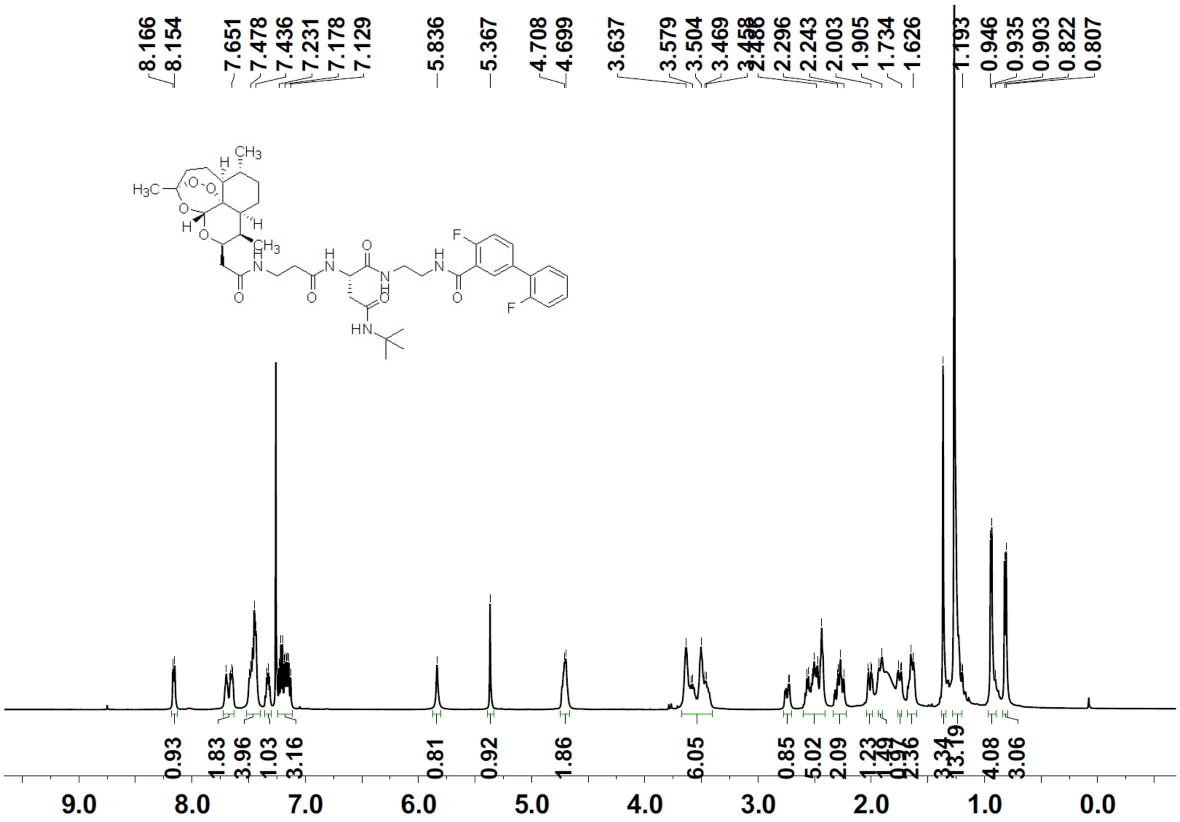
8.166
8.154
7.651
7.478
7.436
7.231
7.178
7.129
5.836
5.367
4.708
4.699
3.637
3.579
3.504
3.469
2.296
2.243
2.003
1.905
1.734
1.626
1.193
0.946
0.935
0.903
0.822
0.807
0.93
1.83
3.96
1.03
3.16
0.81
0.92
1.86
6.05
0.85
5.02
2.09
4.08
3.06
9.0
8.0
7.0
6.0
5.0
4.0
3.0
2.0
1.0
0.0

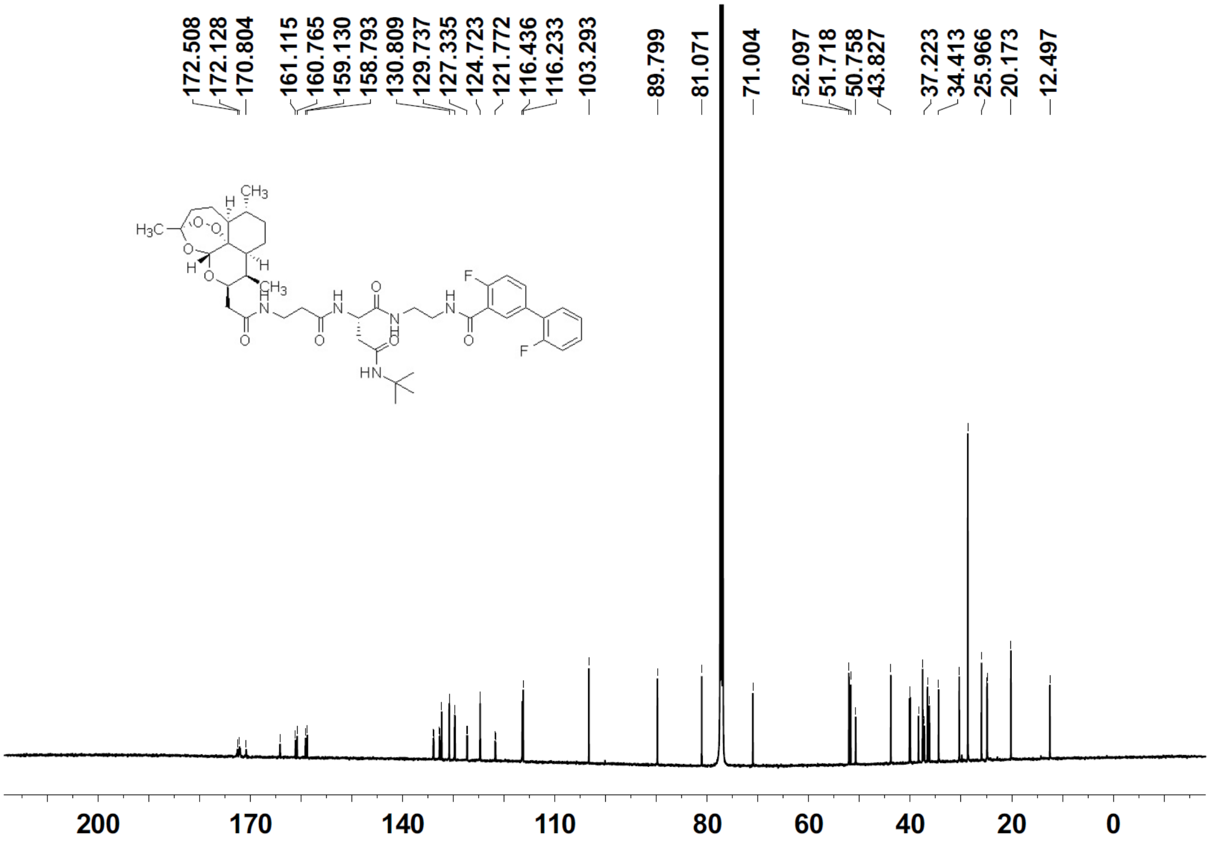
172.508
172.128
170.804
161.115
160.765
159.130
158.793
130.809
129.737
127.335
124.723
121.772
116.436
116.233
103.293
89.799
81.071
71.004
52.097
51.718
50.758
43.827
37.223
34.413
25.966
20.173
12.497
200
170
140
110
80
60
40
20
0

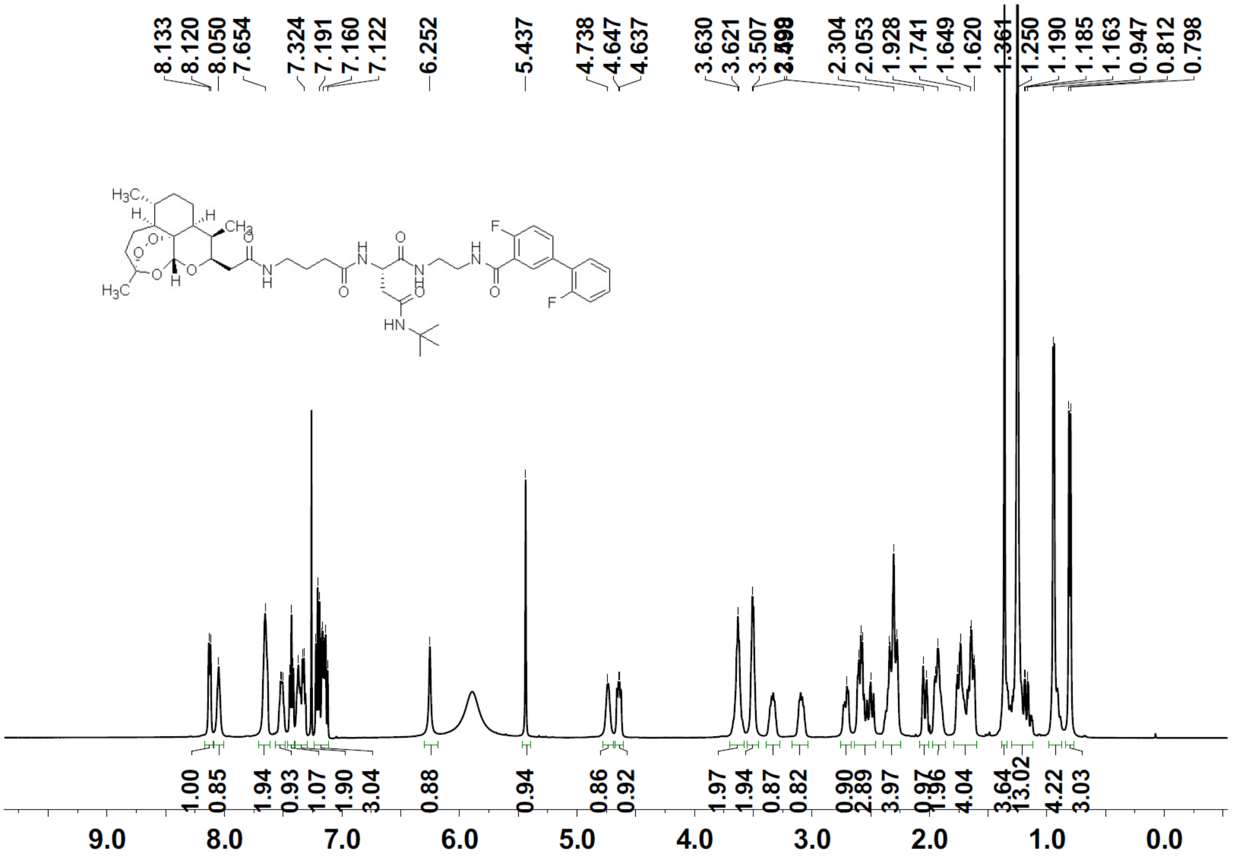
8.133
8.120
8.050
7.654
7.324
7.191
7.160
7.122
6.252
5.437
4.738
4.647
4.637
3.630
3.621
3.507
2.304
2.053
1.928
1.741
1.649
1.620
1.361
1.250
1.190
1.185
1.163
0.947
0.812
0.798
1.00
0.85
1.94
0.93
1.07
1.90
3.04
0.88
0.94
0.86
0.92
1.97
1.94
0.87
0.82
0.90
2.89
3.97
4.04
4.22
3.03
9.0
8.0
7.0
6.0
5.0
4.0
3.0
2.0
1.0
0.0

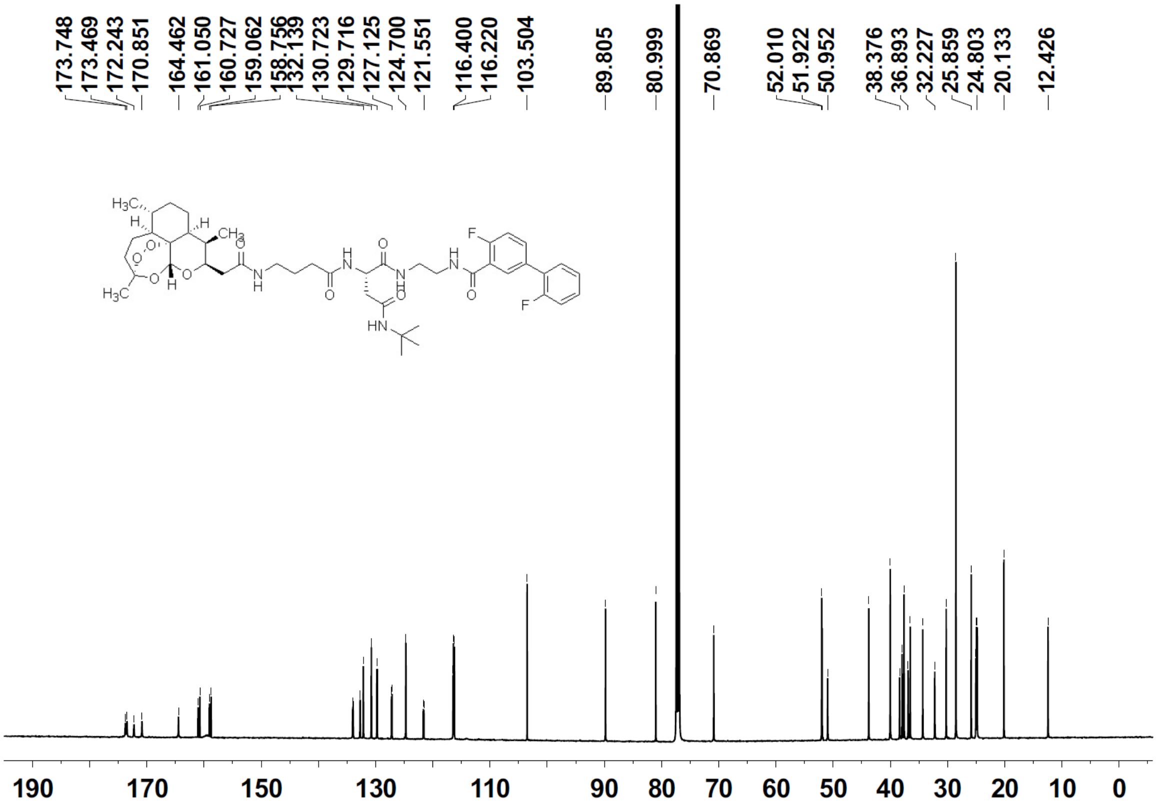
173.748
173.469
172.243
170.851
164.462
161.050
160.727
159.062
158.756
132.139
130.723
129.716
127.125
124.700
121.551
116.400
116.220
103.504
89.805
80.999
70.869
52.010
51.922
50.952
38.376
36.893
32.227
25.859
24.803
20.133
12.426
190
170
150
130
110
90
80
70
60
50
40
30
20
10
0

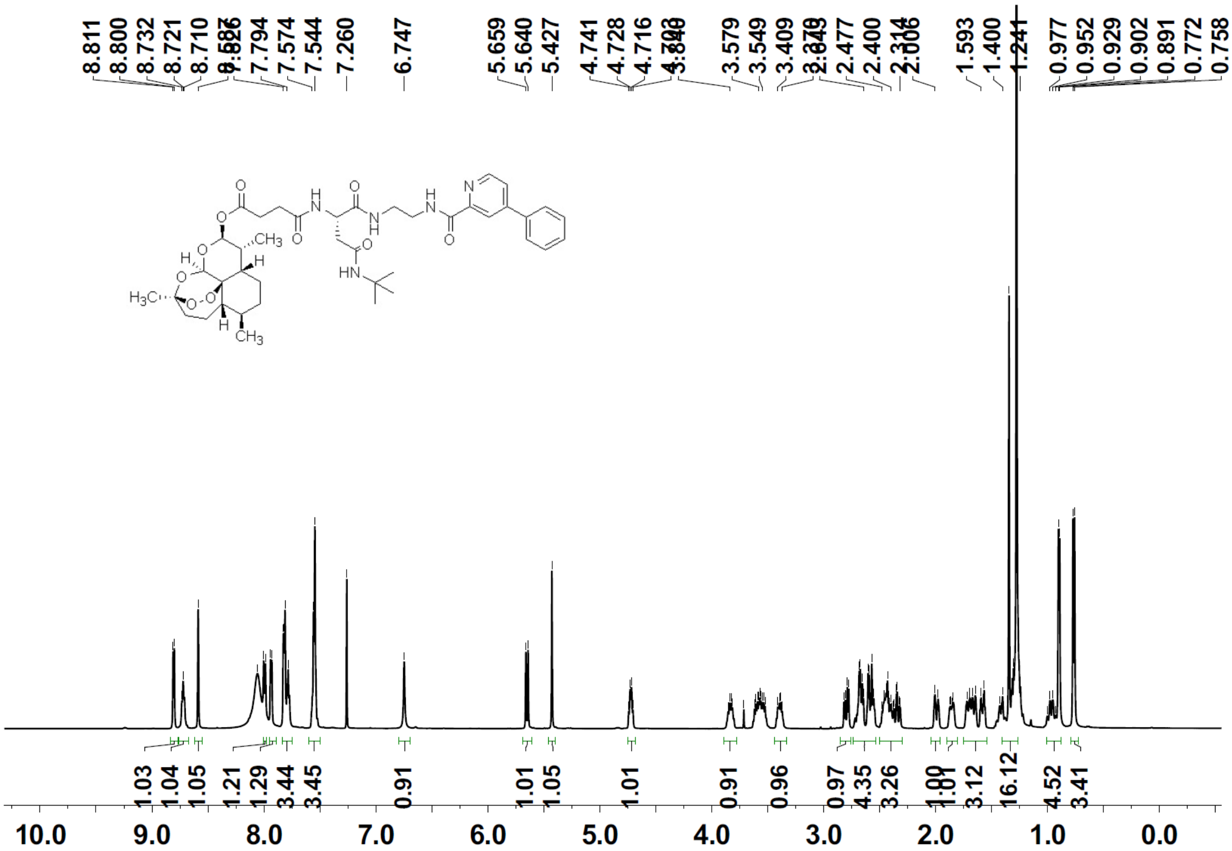
8.811
8.800
8.732
8.721
8.710
7.794
7.574
7.544
7.260
6.747
5.659
5.640
5.427
4.741
4.728
4.716
3.579
3.549
3.409
2.477
2.400
1.593
1.400
0.977
0.952
0.929
0.902
0.891
0.772
0.758
1.03
1.04
1.05
1.21
1.29
3.44
3.45
0.91
1.01
1.05
1.01
0.91
0.96
0.97
4.35
3.26
3.12
16.12
4.52
3.41
10.0
9.0
8.0
7.0
6.0
5.0
4.0
3.0
2.0
1.0
0.0

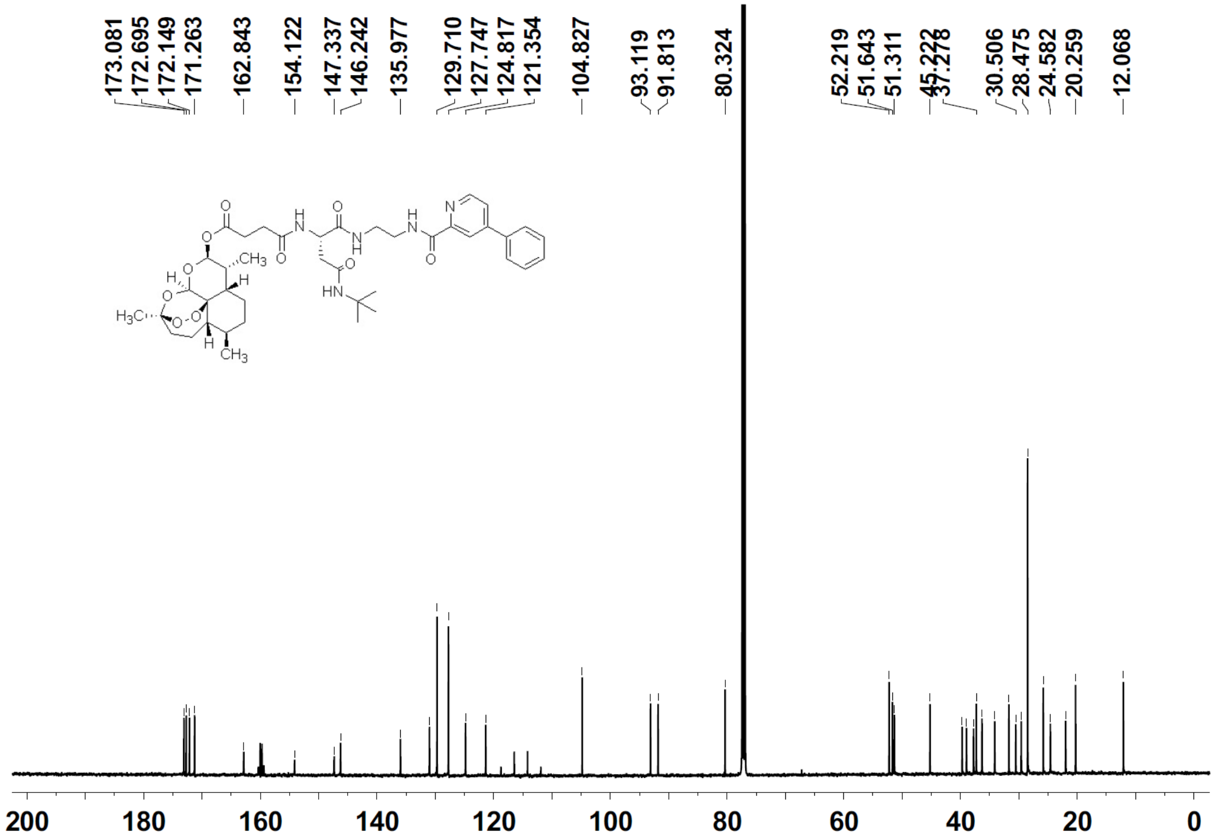
173.081
172.695
172.149
171.263
162.843
154.122
147.337
146.242
135.977
129.710
127.747
124.817
121.354
104.827
93.119
91.813
80.324
52.219
51.643
51.311
30.506
28.475
24.582
20.259
12.068
200
180
160
140
120
100
80
60
40
20
0

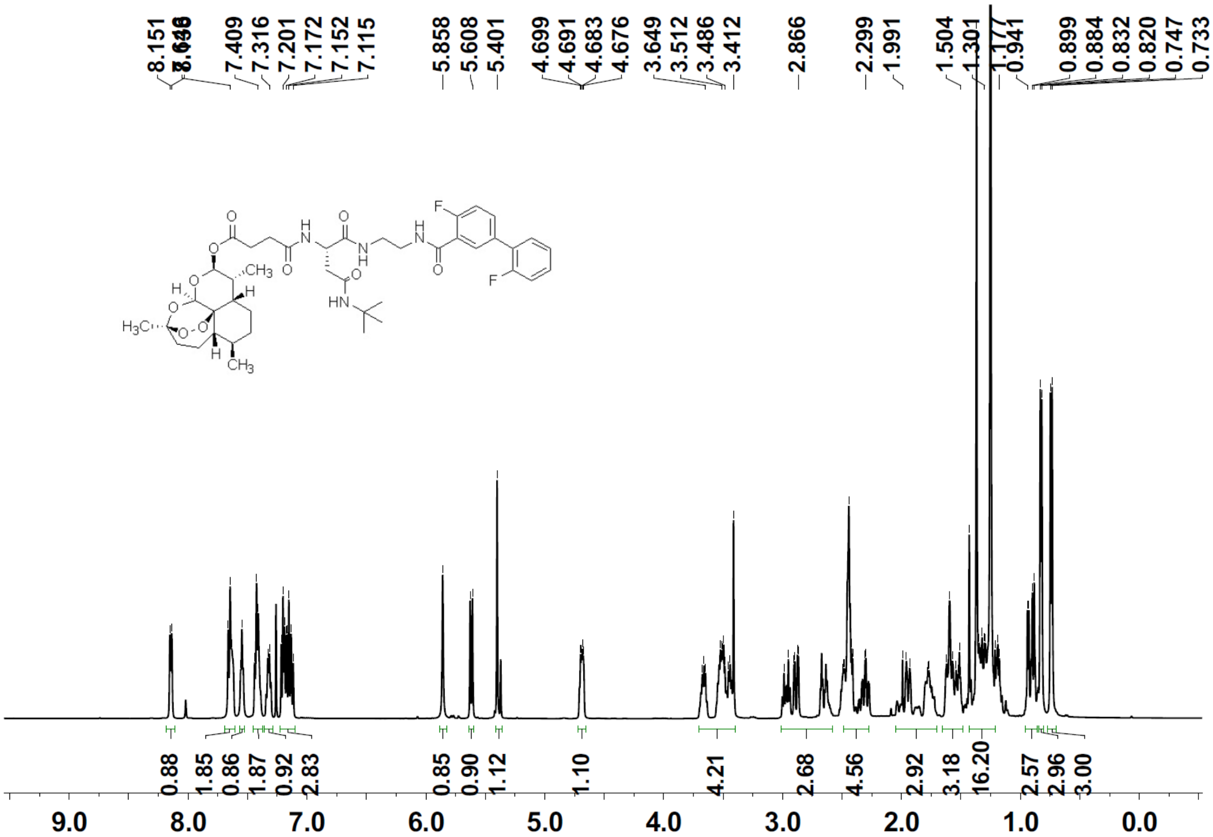
8.151
7.409
7.316
7.201
7.172
7.152
7.115
5.858
5.608
5.401
4.699
4.691
4.683
4.676
3.649
3.512
3.486
3.412
2.866
2.299
1.991
1.504
1.301
0.941
0.899
0.884
0.832
0.820
0.747
0.733
0.88
1.85
0.86
1.87
0.92
2.83
0.85
0.90
1.12
1.10
4.21
2.68
4.56
2.92
3.18
16.20
2.57
2.96
3.00
9.0
8.0
7.0
6.0
5.0
4.0
3.0
2.0
1.0
0.0

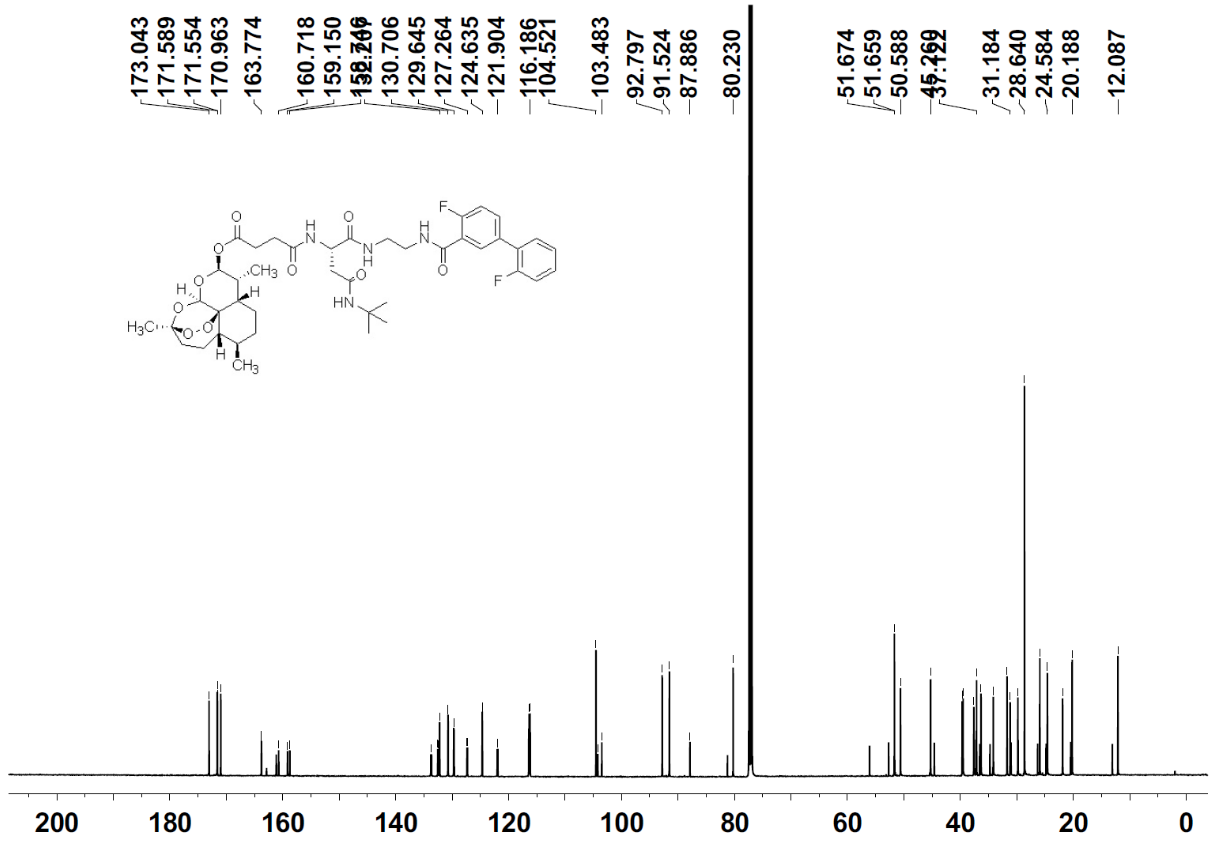
173.043
171.589
171.554
170.963
163.774
160.718
159.150
130.706
129.645
127.264
124.635
121.904
116.186
104.521
103.483
92.797
91.524
87.886
80.230
51.674
51.659
50.588
31.184
28.640
24.584
20.188
12.087
200
180
160
140
120
100
80
60
40
20
0

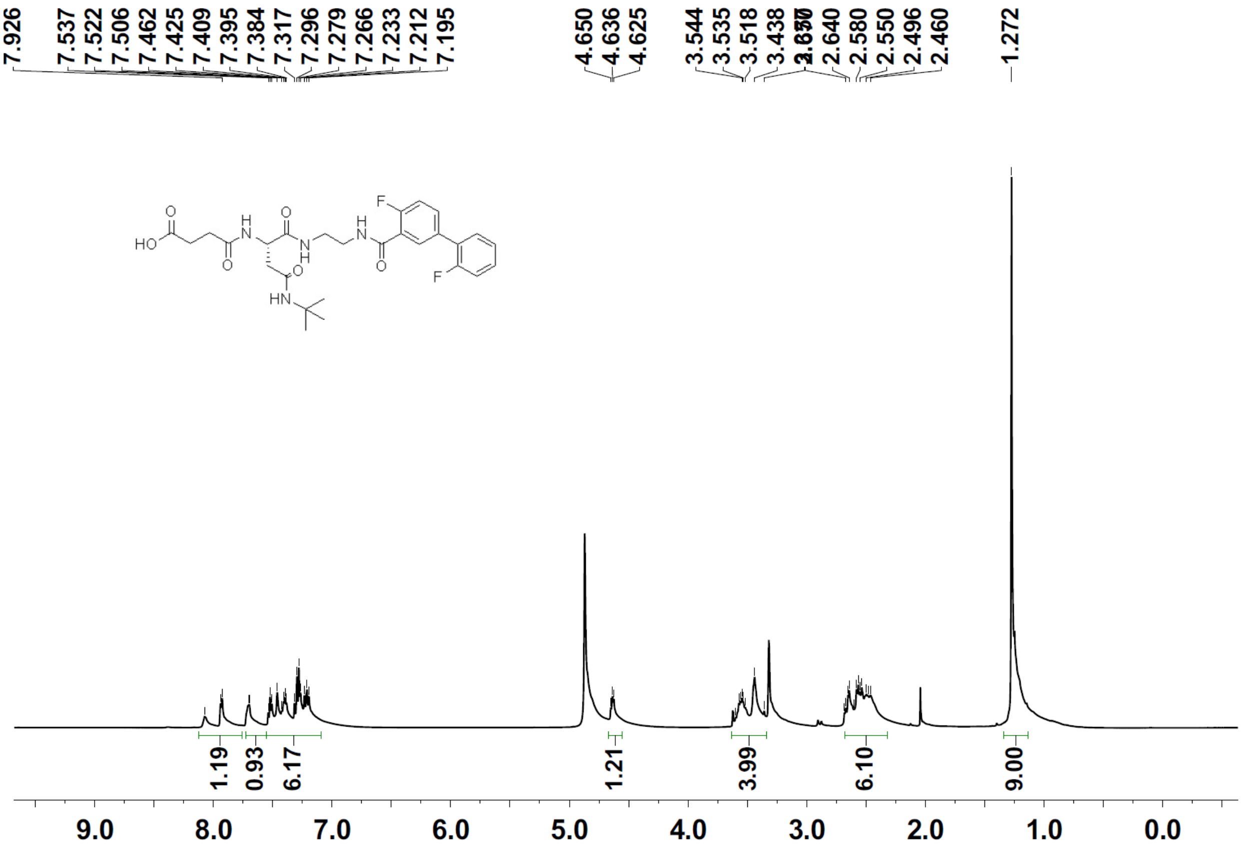
7.926
7.537
7.522
7.506
7.462
7.425
7.409
7.395
7.384
7.317
7.296
7.279
7.266
7.233
7.212
7.195
4.650
4.636
4.625
3.544
3.535
3.518
3.438
2.640
2.580
2.550
2.496
2.460
1.272
HO
O
H
N
O
O
N
H
H
N
O
F
F
HN
1.19
0.93
6.17
1.21
3.99
6.10
9.00
9.0
8.0
7.0
6.0
5.0
4.0
3.0
2.0
1.0
0.0

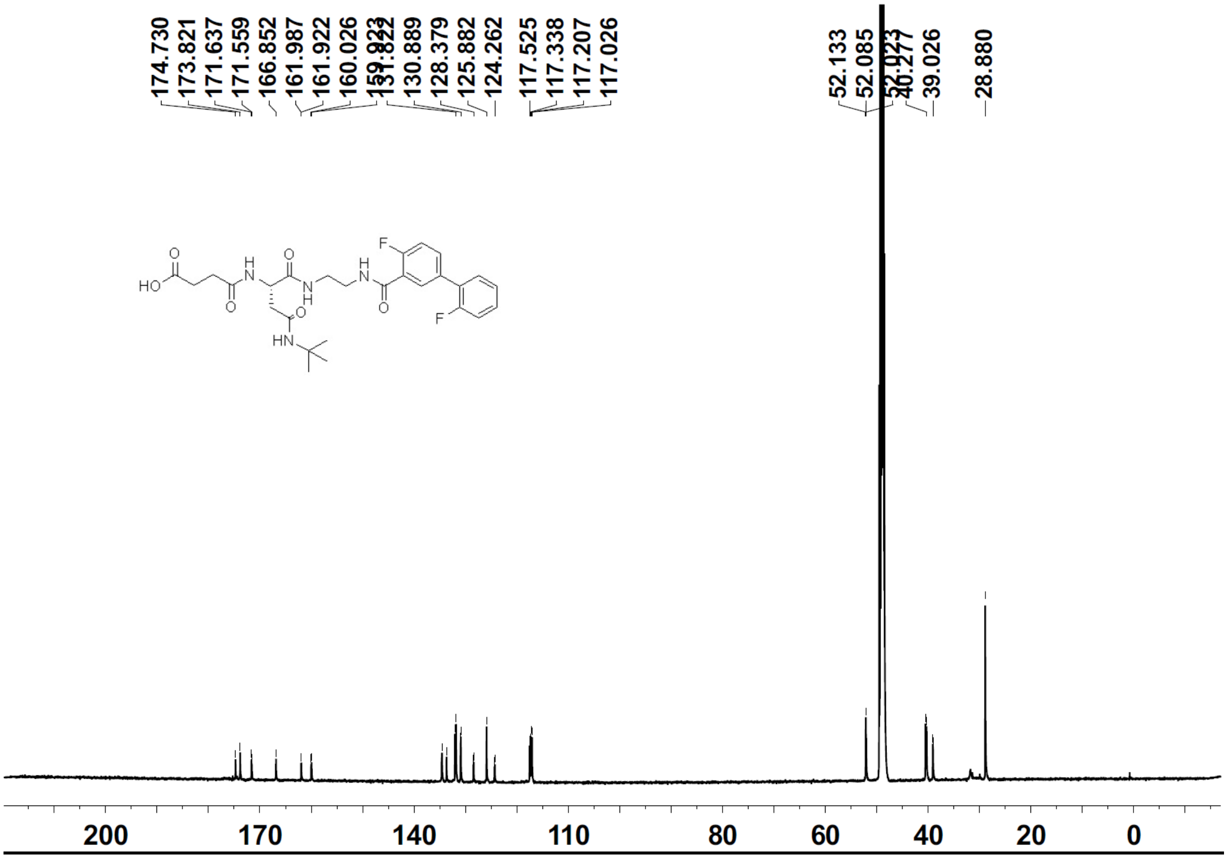
174.730
173.821
171.637
171.559
166.852
161.987
161.922
160.026
130.889
128.379
125.882
124.262
117.525
117.338
117.207
117.026
52.133
52.085
39.026
28.880
HO
O
H
N
O
O
N
H
H
N
O
F
F
HN
200
170
140
110
80
60
40
20
0

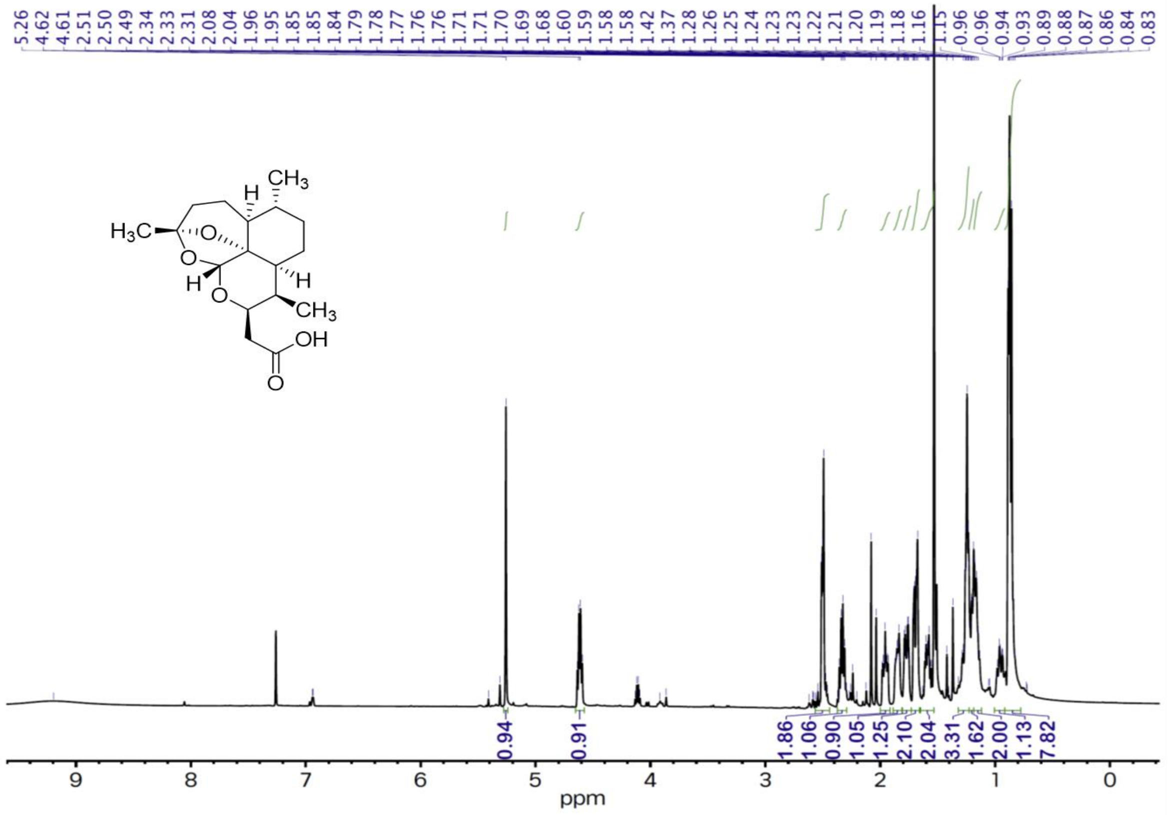
5.26 4.62 4.61 2.51 2.50 2.49 2.34 2.33 2.31 2.08 2.04 1.96 1.95 1.85 1.85 1.84 1.79 1.78 1.77 1.76 1.76 1.71 1.70 1.69 1.68 1.60 1.59 1.58 1.58 1.42 1.37 1.28 1.26 1.25 1.24 1.23 1.23 1.22 1.21 1.20 1.19 1.18 1.16 1.15 0.96 0.96 0.94 0.93 0.89 0.88 0.87 0.86 0.84 0.83
CH3
H
H3C
O
O
H
O
H
CH3
OH
O
0.94 0.91 1.86 1.06 0.90 1.05 1.25 2.10 2.04 3.31 1.62 2.00 1.13 7.82
9 8 7 6 5 4 3 2 1 0
ppm

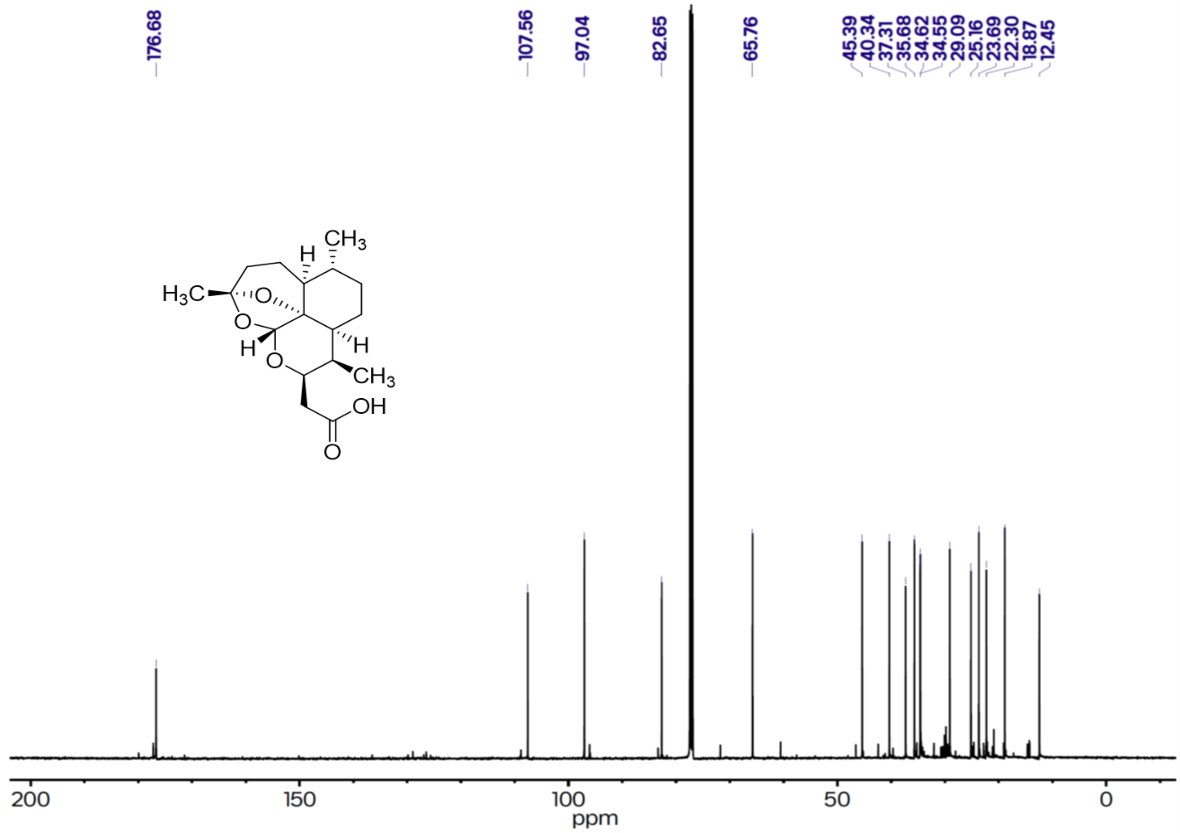
176.68
107.56
97.04
82.65
65.76
45.39 40.34 37.31 35.68 34.62 34.55 29.09 25.16 23.69 22.30 18.87 12.45
CH3
H
H3C
O
O
H
O
H
CH3
OH
O
200 150 100 50 0
ppm

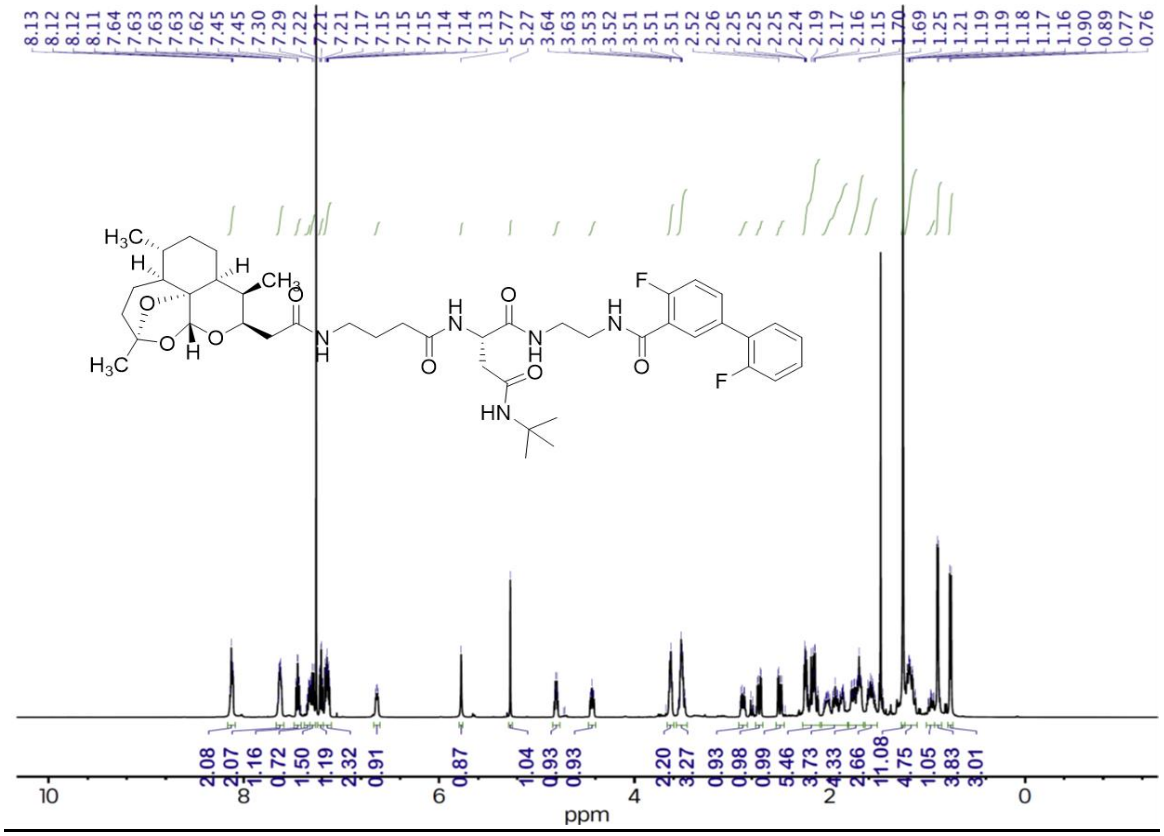
8.13 8.12 8.12 8.11 7.64 7.63 7.63 7.63 7.62 7.45 7.45 7.30 7.29 7.22 7.21 7.21 7.17 7.15 7.15 7.15 7.14 7.14 7.13 5.77 5.27 3.64 3.63 3.53 3.52 3.51 3.51 3.51 2.52 2.26 2.25 2.25 2.25 2.24 2.19 2.17 2.16 2.15 1.70 1.69 1.25 1.21 1.19 1.19 1.18 1.17 1.16 0.90 0.89 0.77 0.76
2.08 2.07 1.16 0.72 1.50 1.19 2.32 0.91 0.87 1.04 0.93 0.93 2.20 3.27 0.93 0.98 0.99 5.46 3.73 4.33 2.66 11.08 4.75 1.05 3.83 3.01
ppm

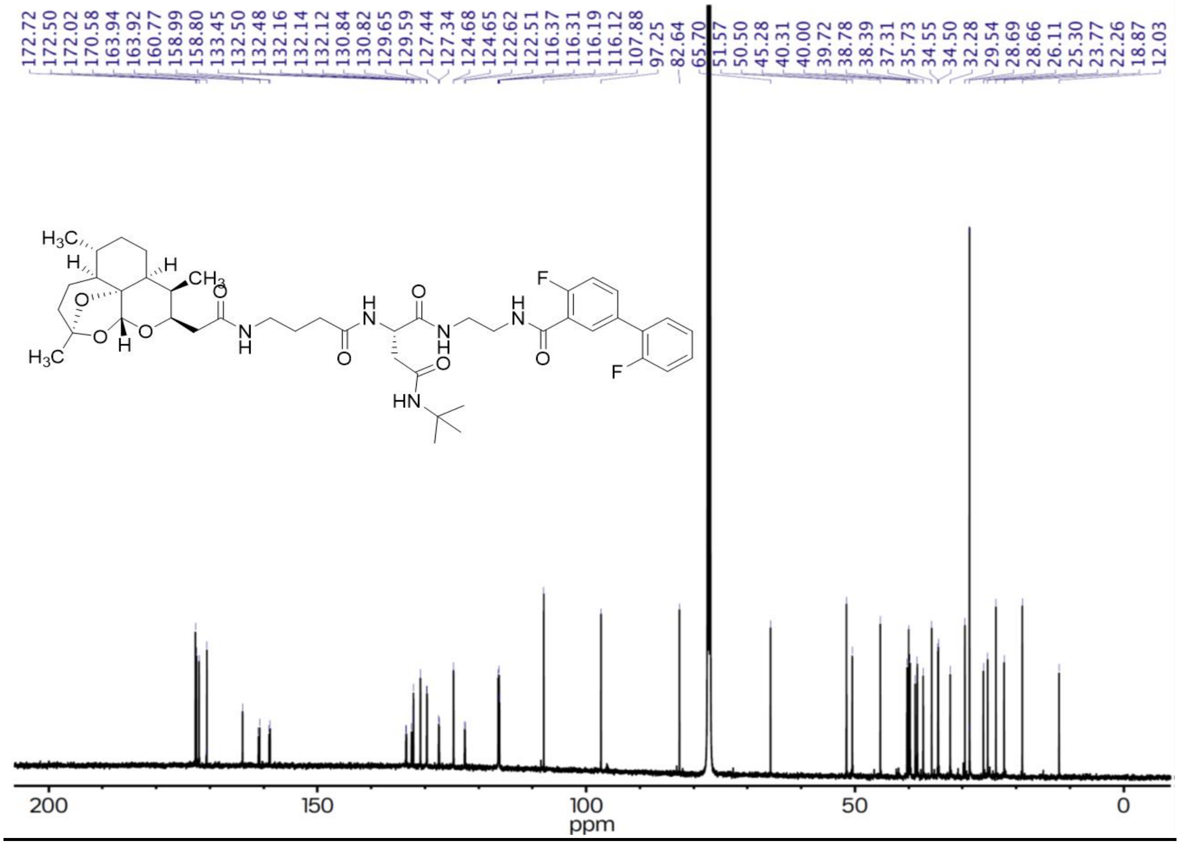
172.72 172.50 172.02 170.58 163.94 163.92 160.77 158.99 158.80 133.45 132.50 132.48 132.16 132.14 132.12 130.84 130.82 129.65 129.59 127.44 127.34 124.68 124.65 122.62 122.51 116.37 116.31 116.19 116.12 107.88 97.25 82.64 65.70 51.57 50.50 45.28 40.31 40.00 39.72 38.78 38.39 37.31 35.73 34.55 34.50 32.28 29.54 28.69 28.66 26.11 25.30 23.77 22.26 18.87 12.03
ppm

**LCMS**

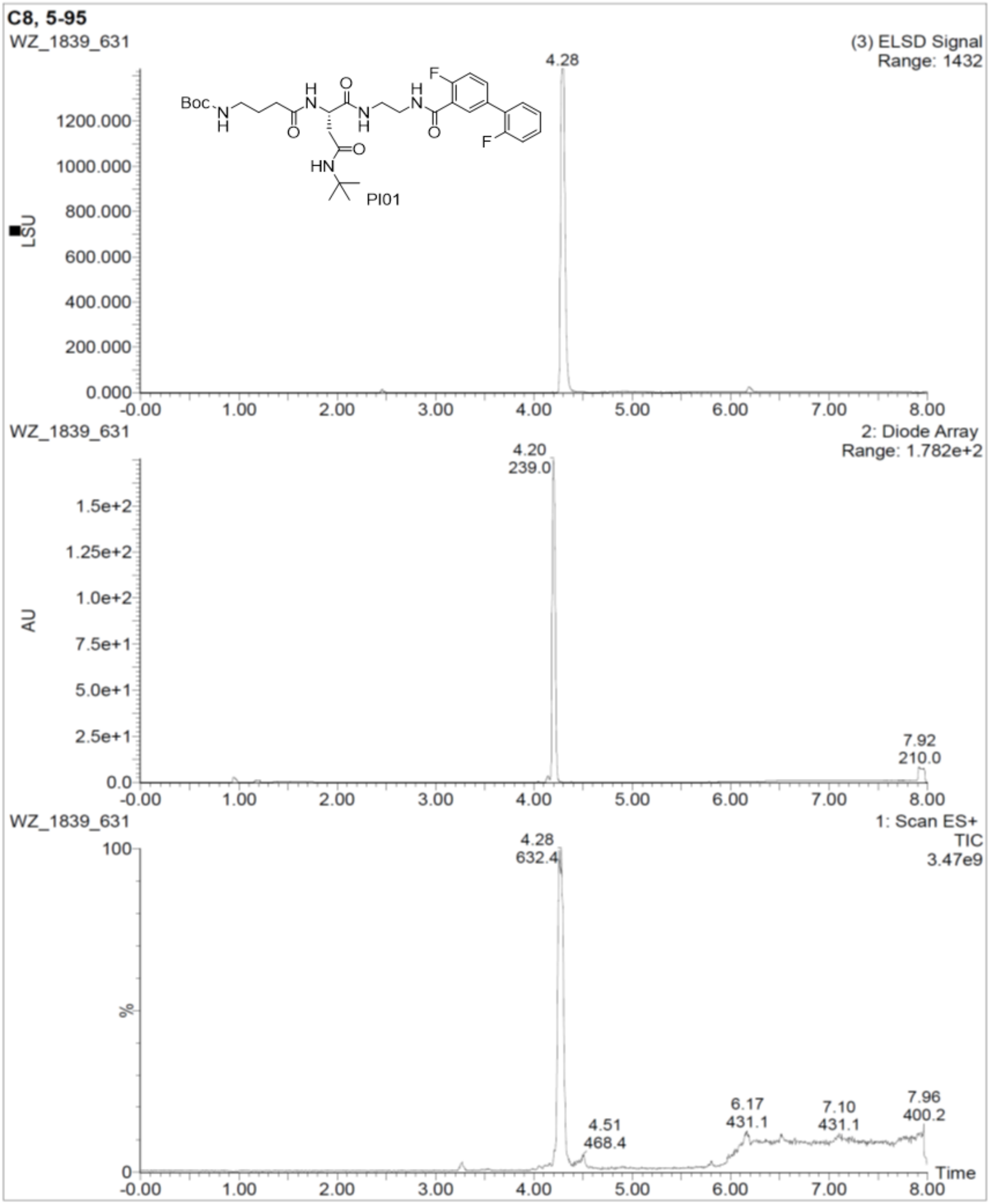
C8, 5-95
WZ_1839_631
(3) ELSD Signal
Range: 1432
4.28
PI01
LSU
2: Diode Array
Range: 1.782e+2
4.20
239.0
7.92
210.0
AU
1: Scan ES+
TIC
3.47e9
4.28
632.4
4.51
468.4
6.17
431.1
7.10
431.1
7.96
400.2
%
Time

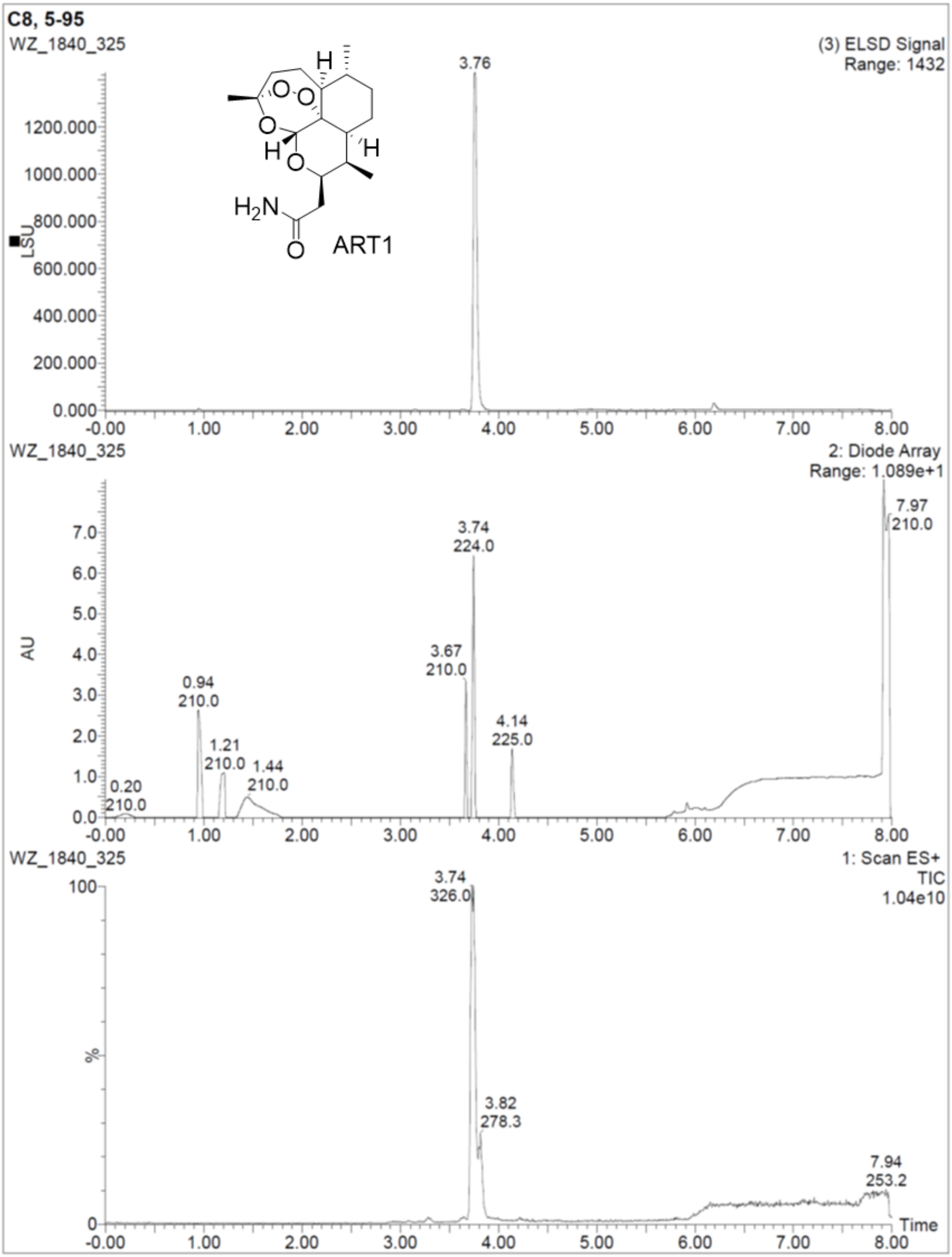
C8, 5-95
WZ_1840_325
(3) ELSD Signal
Range: 1432
3.76
ART1
LSU
1200.000
1000.000
800.000
600.000
400.000
200.000
0.000
-0.00
1.00
2.00
3.00
4.00
5.00
6.00
7.00
8.00
WZ_1840_325
2: Diode Array
Range: 1.089e+1
AU
7.97
210.0
3.74
224.0
3.67
210.0
0.94
210.0
4.14
225.0
1.21
210.0
1.44
210.0
0.20
210.0
WZ_1840_325
1: Scan ES+
TIC
1.04e10
3.74
326.0
3.82
278.3
7.94
253.2
%
Time

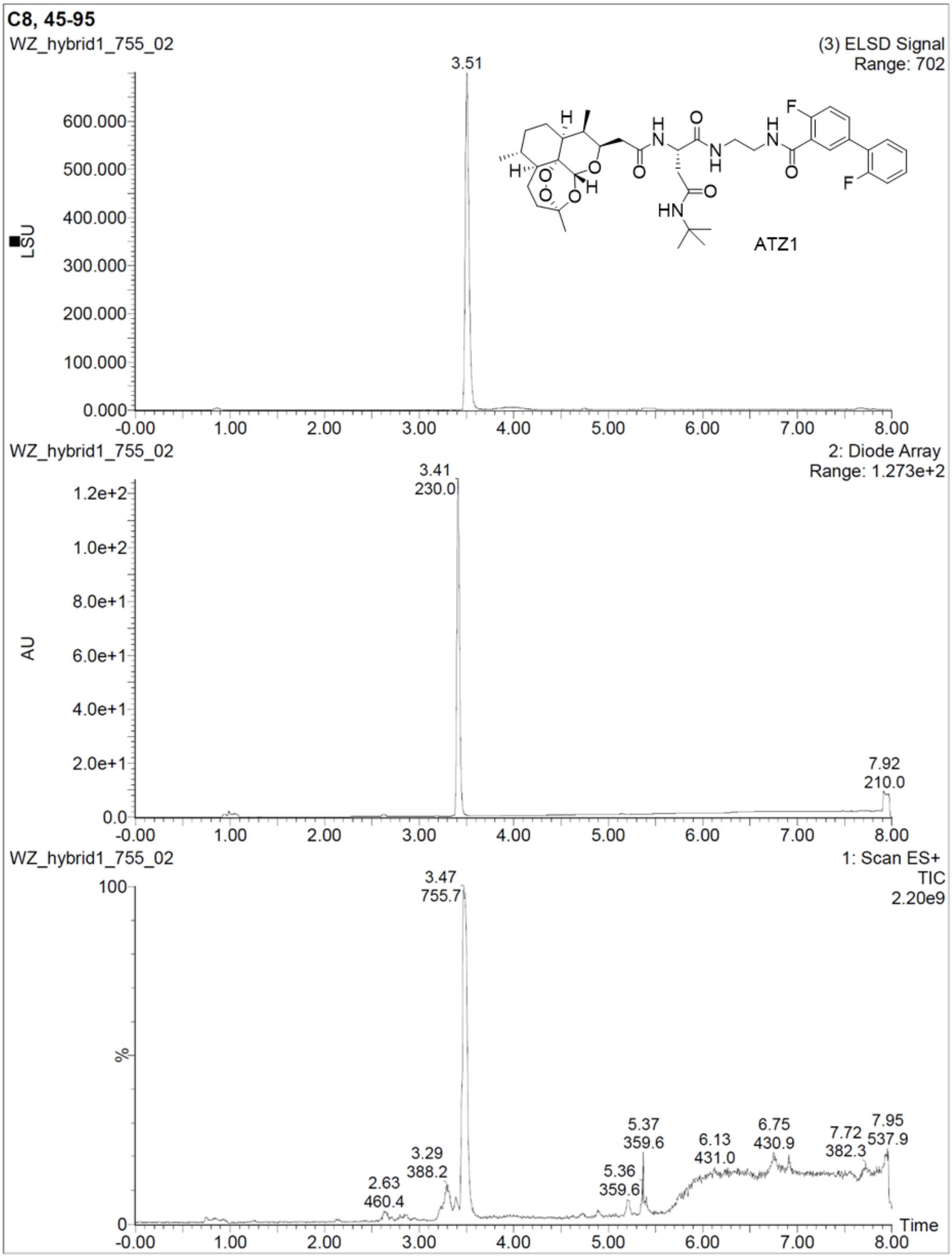
C8, 45-95
WZ_hybrid1_755_02
(3) ELSD Signal
Range: 702
3.51
LSU
ATZ1
WZ_hybrid1_755_02
2: Diode Array
Range: 1.273e+2
3.41
230.0
AU
7.92
210.0
WZ_hybrid1_755_02
1: Scan ES+
TIC
2.20e9
3.47
755.7
%
2.63
460.4
3.29
388.2
5.36
359.6
5.37
359.6
6.13
431.0
6.75
430.9
7.72
382.3
7.95
537.9
Time

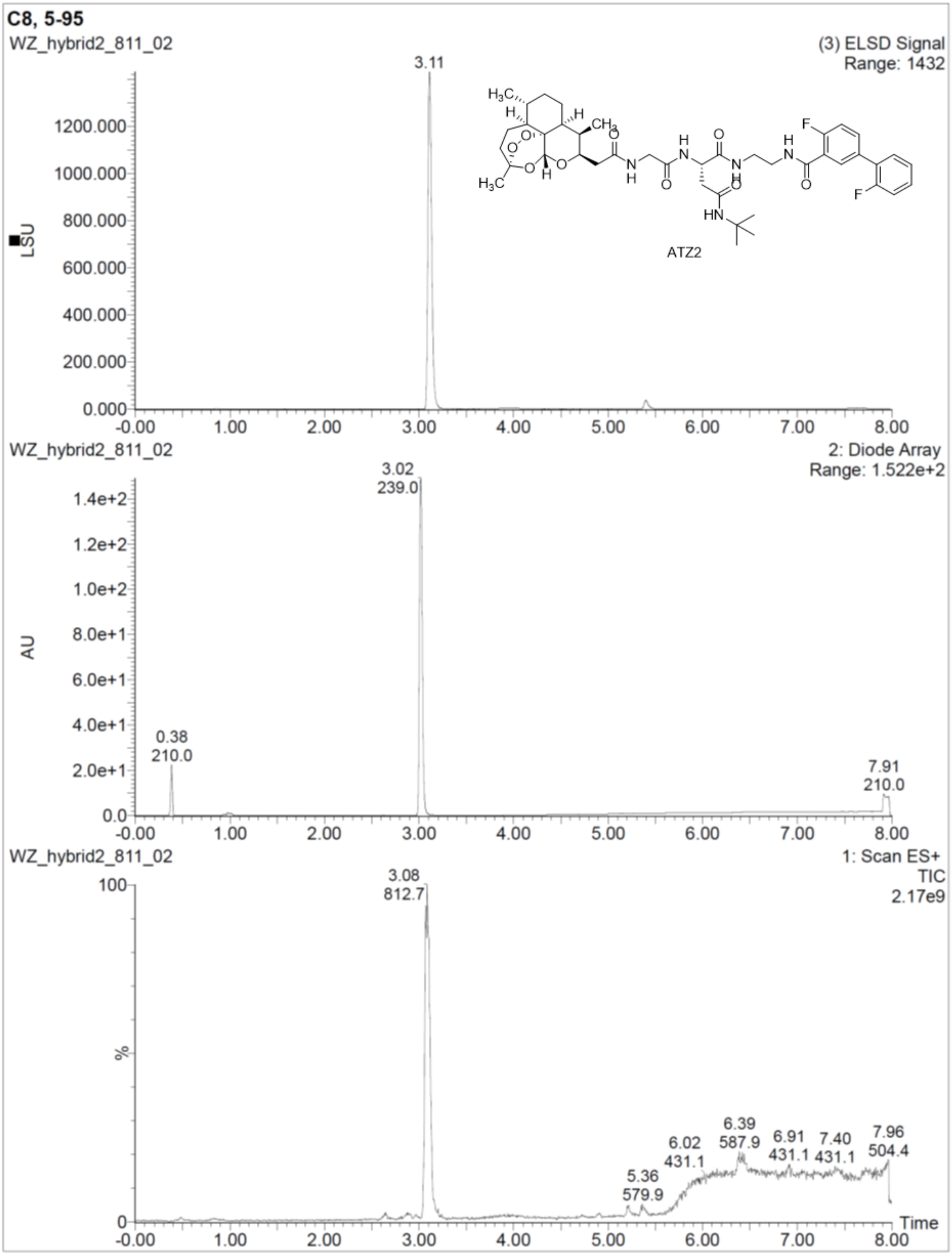
C8, 5-95
WZ_hybrid2_811_02
(3) ELSD Signal
Range: 1432
3.11
LSU
ATZ2
WZ_hybrid2_811_02
2: Diode Array
Range: 1.522e+2
3.02
239.0
AU
0.38
210.0
7.91
210.0
WZ_hybrid2_811_02
1: Scan ES+
TIC
2.17e9
3.08
812.7
%
5.36
579.9
6.02
431.1
6.39
587.9
6.91
431.1
7.40
431.1
7.96
504.4
Time

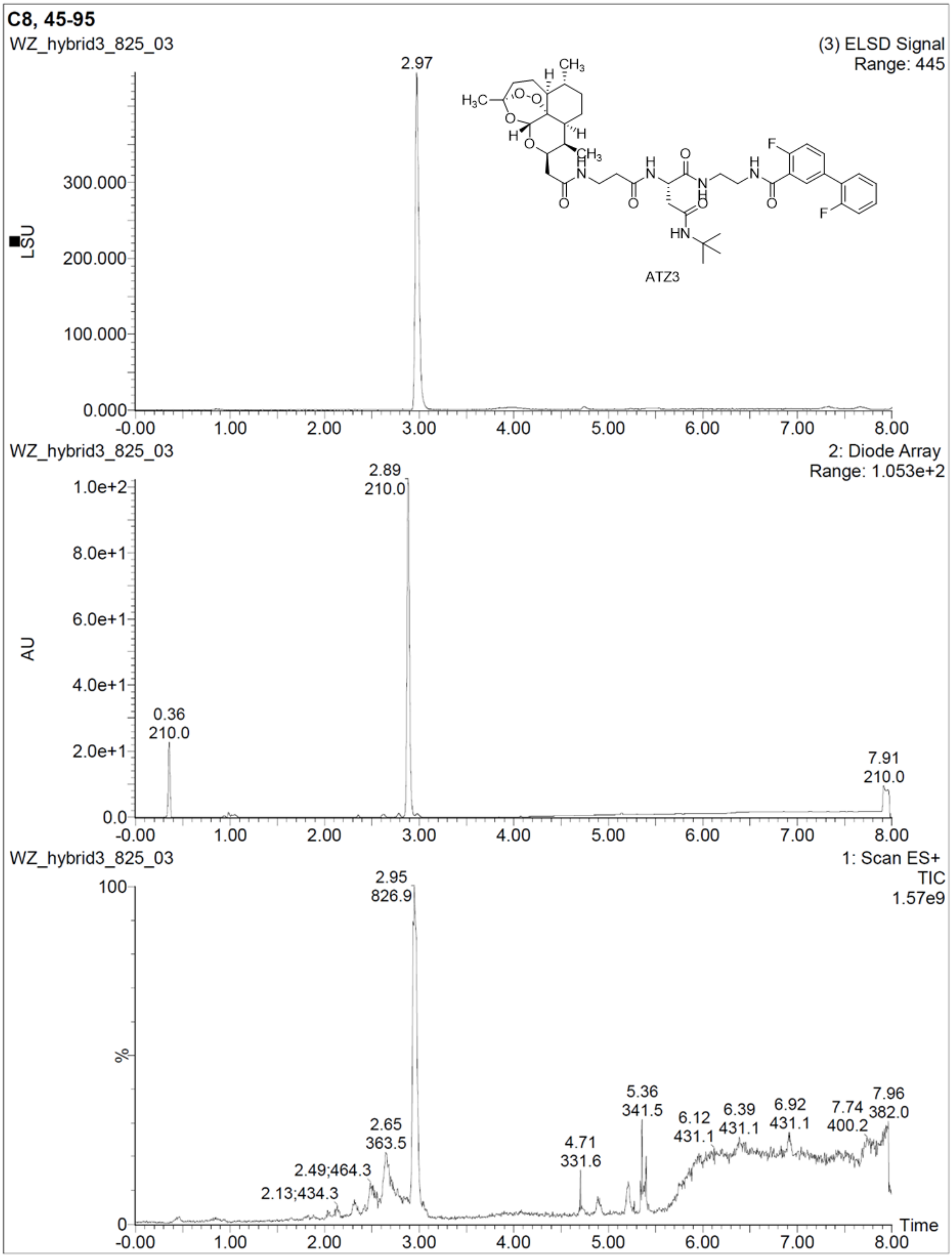
C8, 45-95
WZ_hybrid3_825_03
(3) ELSD Signal
Range: 445
2.97
LSU
300.000
200.000
100.000
0.000
ATZ3
-0.00
1.00
2.00
3.00
4.00
5.00
6.00
7.00
8.00
WZ_hybrid3_825_03
2: Diode Array
Range: 1.053e+2
2.89
210.0
1.0e+2
8.0e+1
6.0e+1
4.0e+1
2.0e+1
0.0
AU
0.36
210.0
7.91
210.0
WZ_hybrid3_825_03
1: Scan ES+
TIC
1.57e9
2.95
826.9
100
%
0
2.13;434.3
2.49;464.3
2.65
363.5
4.71
331.6
5.36
341.5
6.12
431.1
6.39
431.1
6.92
431.1
7.74
400.2
7.96
382.0
Time

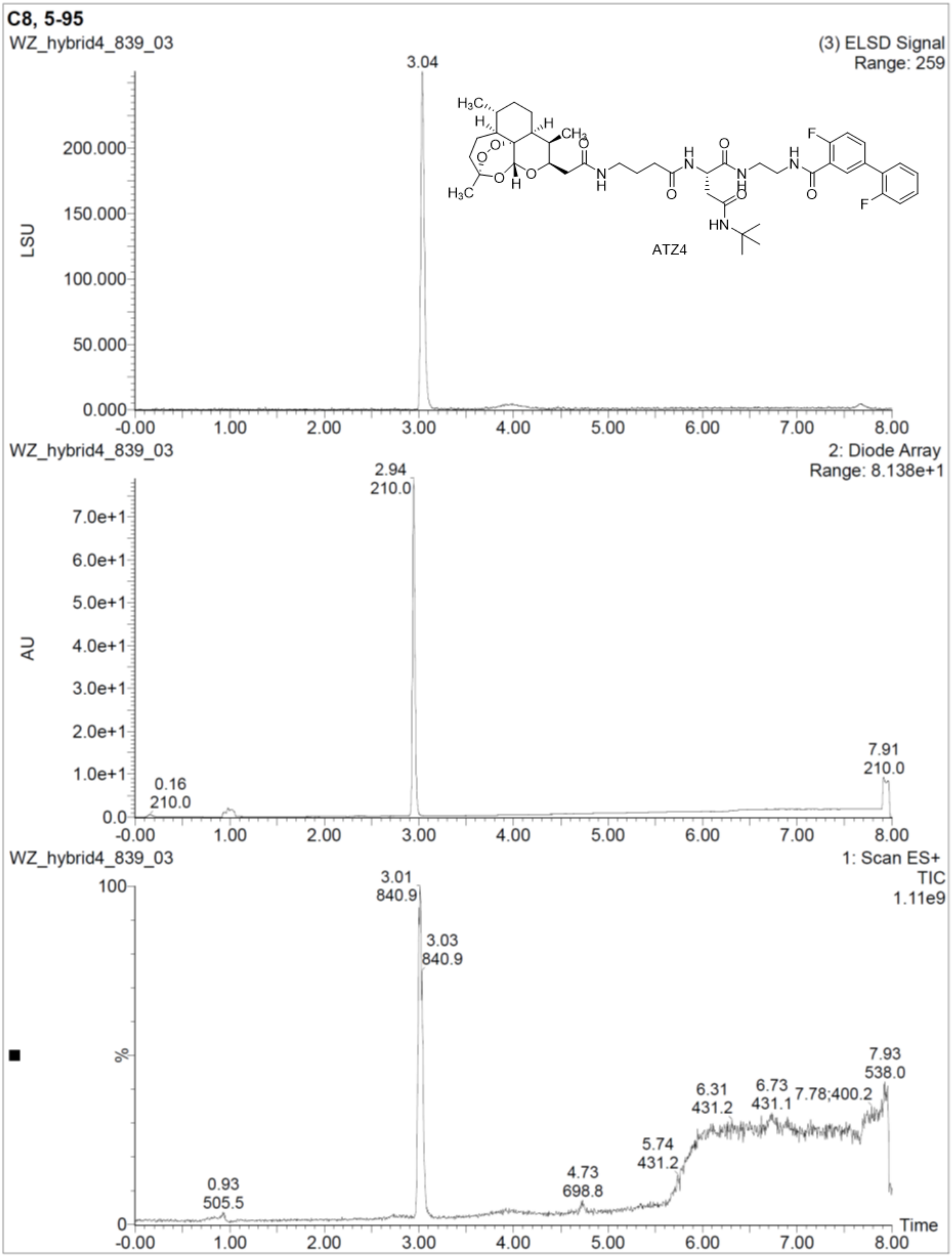
C8, 5-95
WZ_hybrid4_839_03
(3) ELSD Signal
Range: 259
3.04
LSU
ATZ4
WZ_hybrid4_839_03
2: Diode Array
Range: 8.138e+1
2.94
210.0
AU
0.16
210.0
7.91
210.0
WZ_hybrid4_839_03
1: Scan ES+
TIC
1.11e9
3.01
840.9
3.03
840.9
%
0.93
505.5
4.73
698.8
5.74
431.2
6.31
431.2
6.73
431.1
7.78;400.2
7.93
538.0
Time

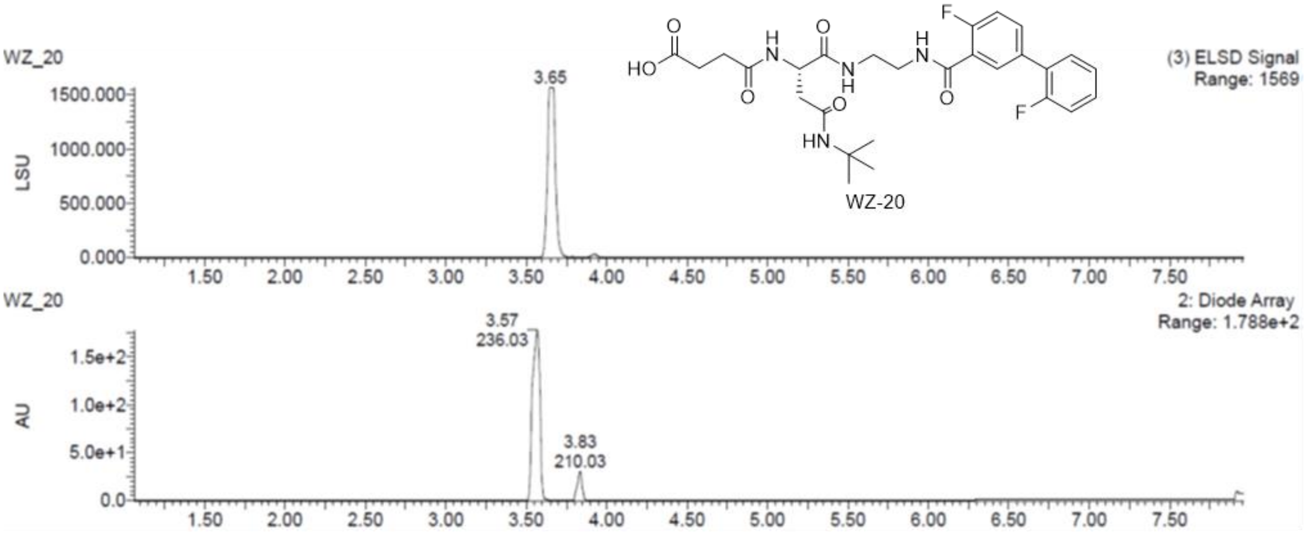
WZ_20
(3) ELSD Signal
Range: 1569
3.65
1500.000
1000.000
500.000
0.000
LSU
HO
O
H
N
F
WZ-20
WZ_20
2: Diode Array
Range: 1.788e+2
3.57
236.03
3.83
210.03
1.5e+2
1.0e+2
5.0e+1
0.0
AU
1.50 2.00 2.50 3.00 3.50 4.00 4.50 5.00 5.50 6.00 6.50 7.00 7.50

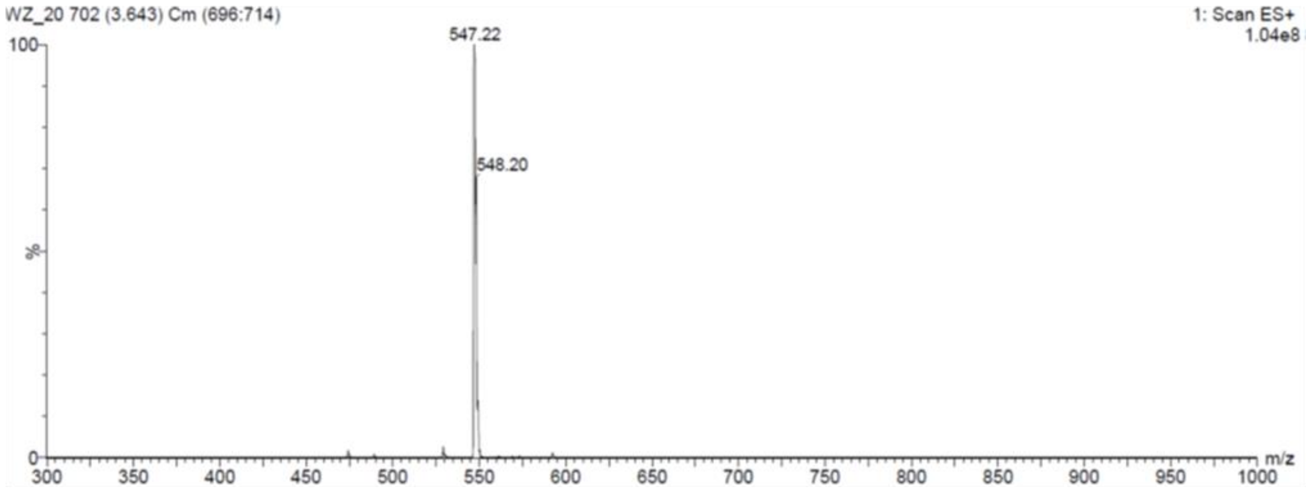
WZ_20 702 (3.643) Cm (696:714)
1: Scan ES+
1.04e8
547.22
548.20
100
%
0
300 350 400 450 500 550 600 650 700 750 800 850 900 950 1000
m/z

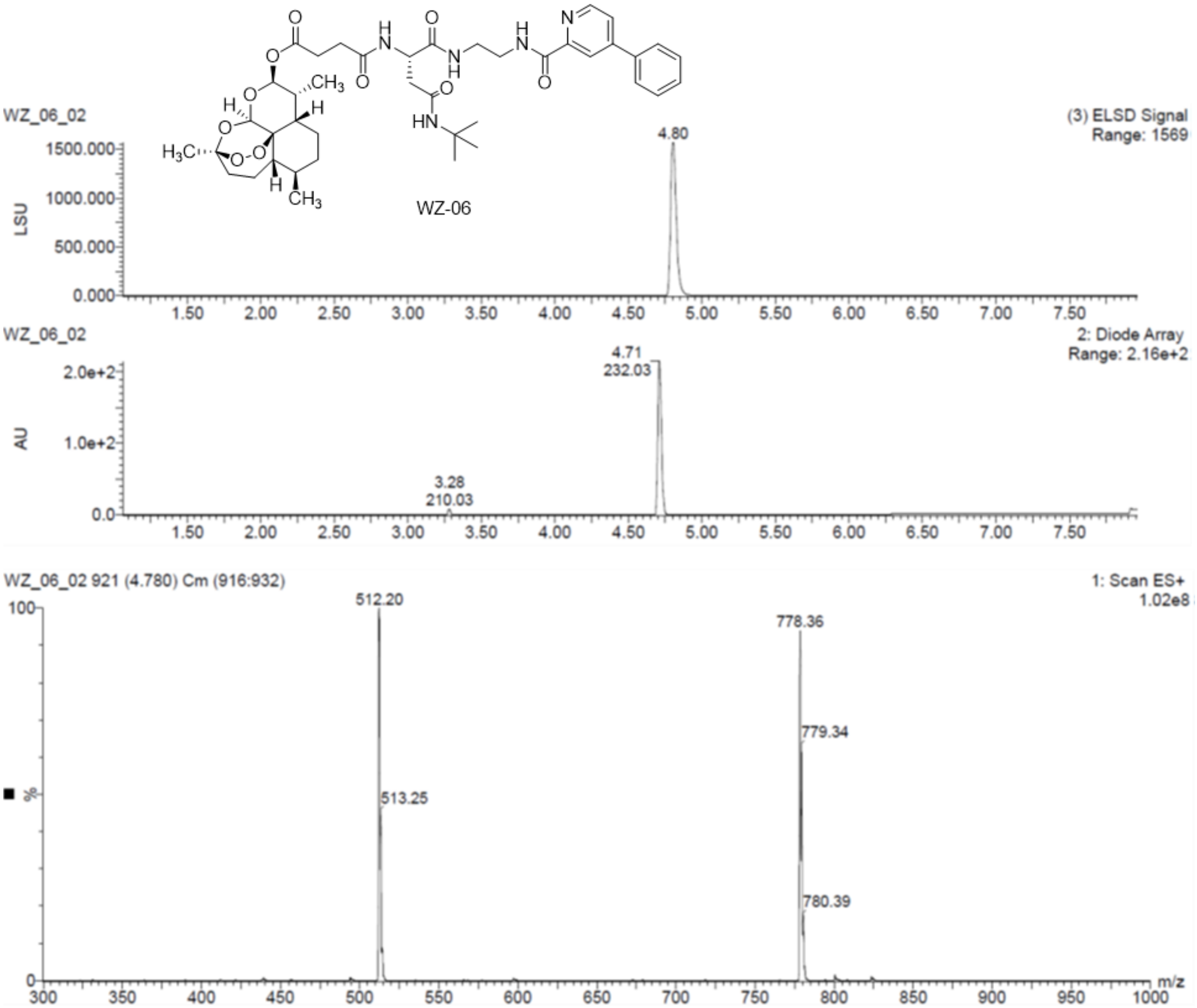
WZ-06
WZ_06_02
(3) ELSD Signal
Range: 1569
4.80
LSU
WZ_06_02
2: Diode Array
Range: 2.16e+2
4.71
232.03
3.28
210.03
AU
WZ_06_02 921 (4.780) Cm (916:932)
1: Scan ES+
1.02e8
512.20
513.25
778.36
779.34
780.39
%
m/z

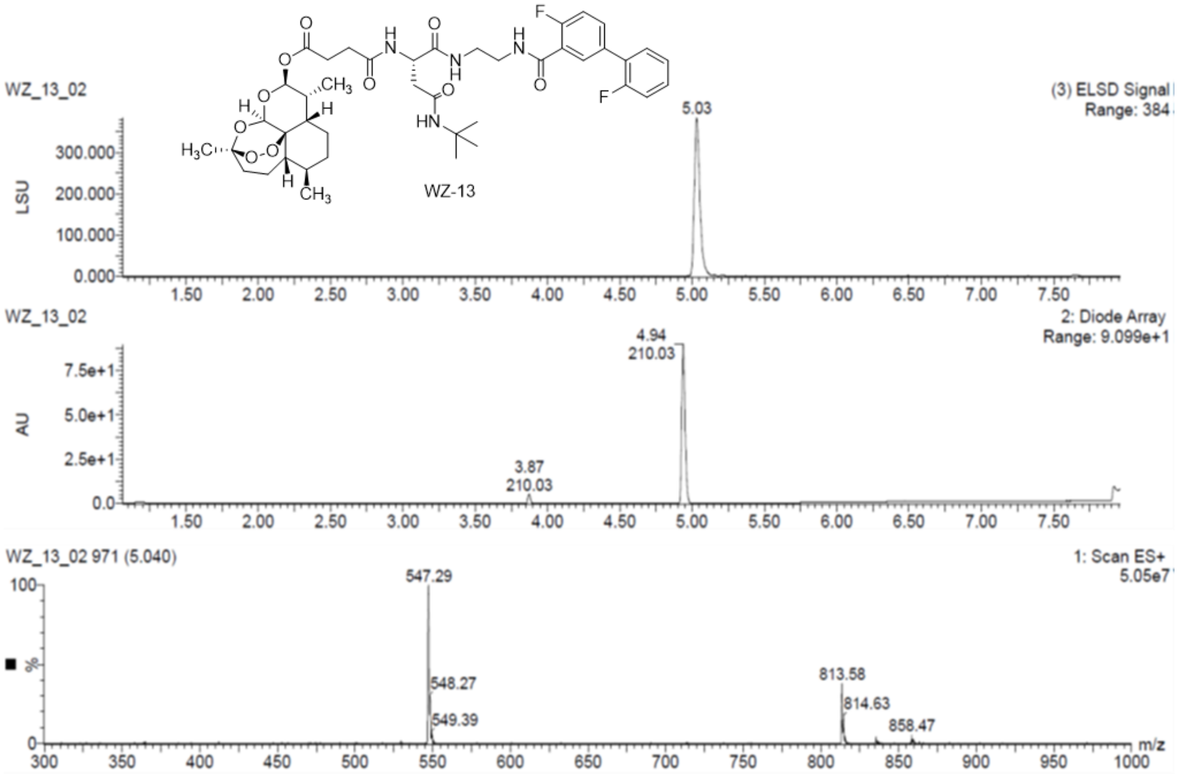
WZ_13_02
WZ-13
(3) ELSD Signal
Range: 384
LSU
300.000
200.000
100.000
0.000
5.03
2: Diode Array
Range: 9.099e+1
AU
7.5e+1
5.0e+1
2.5e+1
0.0
4.94
210.03
3.87
210.03
1.50
2.00
2.50
3.00
3.50
4.00
4.50
5.00
5.50
6.00
6.50
7.00
7.50
WZ_13_02 971 (5.040)
1: Scan ES+
5.05e7
100
%
0
547.29
548.27
549.39
813.58
814.63
858.47
m/z
300
350
400
450
500
550
600
650
700
750
800
850
900
950
1000

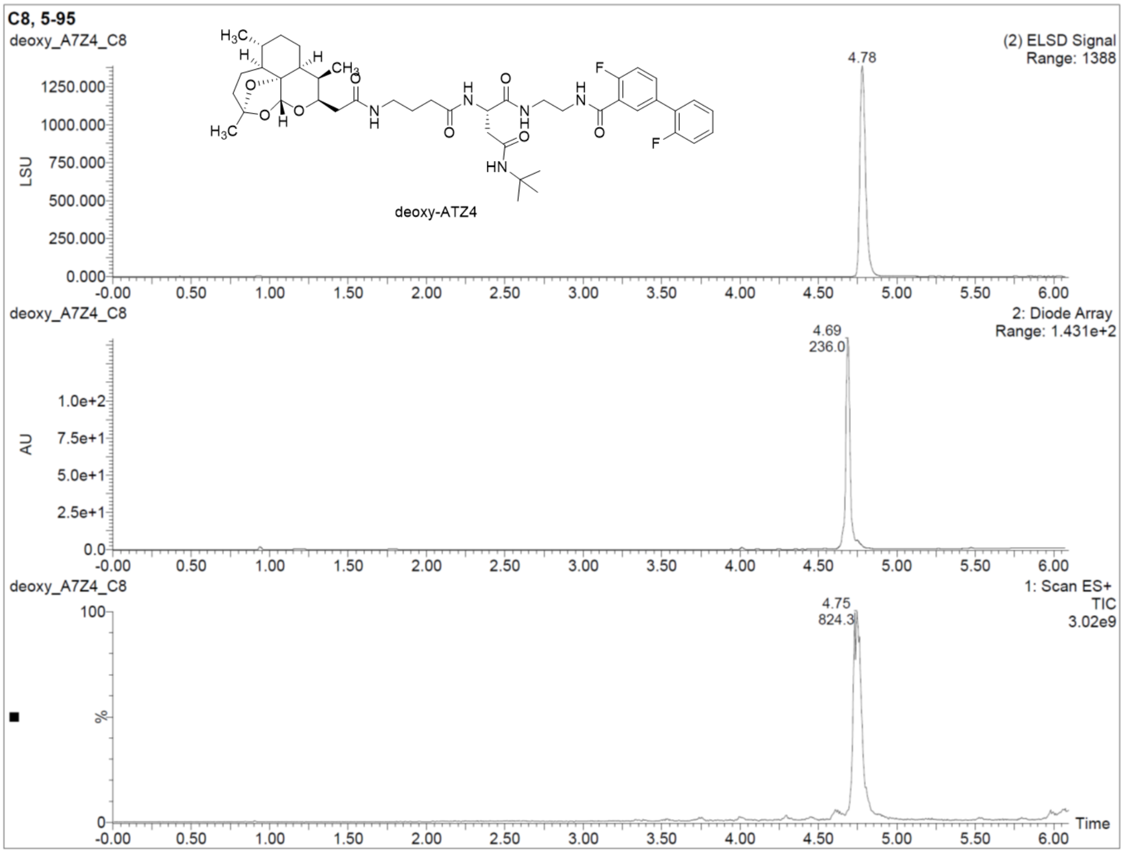
C8, 5-95
deoxy_A7Z4_C8
deoxy-ATZ4
(2) ELSD Signal
Range: 1388
LSU
1250.000
1000.000
750.000
500.000
250.000
0.000
4.78
2: Diode Array
Range: 1.431e+2
AU
1.0e+2
7.5e+1
5.0e+1
2.5e+1
0.0
4.69
236.0
1: Scan ES+
TIC
3.02e9
100
%
0
4.75
824.3
Time
-0.00
0.50
1.00
1.50
2.00
2.50
3.00
3.50
4.00
4.50
5.00
5.50
6.00

**HRMS:**

**Elemental Composition Report** Page 1

### HRMS of PI01

**Single Mass Analysis**
Tolerance = 5.0 PPM / DBE: min = -1.5, max = 50.0
Element prediction: Off
Number of isotope peaks used for i-FIT = 3

Monoisotopic Mass, Even Electron Ions
243 formula(e) evaluated with 1 results within limits (up to 50 closest results for each mass)
Elements Used:
C: 0-32 H: 0-45 N: 0-5 O: 0-6 F: 0-2 Na: 0-1

Rong
Rong
C32H43F2N5O6
WZ_18_39 9 (0.208) Cm (9:16)

NMR Analytical Core Facility
LCT Premier XE

28-Sep-2018
2::1::3

1: TOF MS ES+
5.07e+005

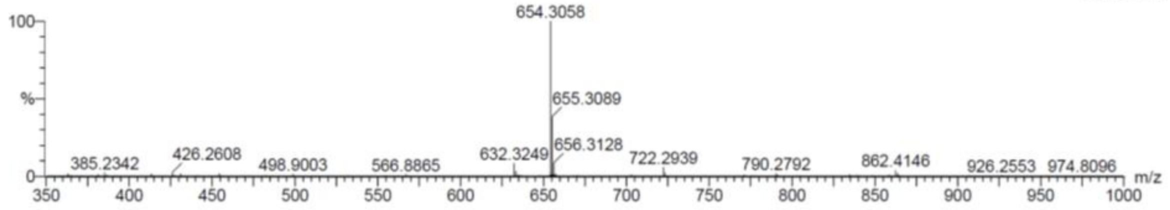

| | | | |
|---|---|---|---|
| Minimum: | | | -1.5 |
| Maximum: | 5.0 | 5.0 | 50.0 |

| Mass | Calc. Mass | mDa | PPM | DBE | i-FIT | i-FIT (Norm) | Formula |
|---|---|---|---|---|---|---|---|
| 654.3058 | 654.3079 | -2.1 | -3.2 | 12.5 | 520.8 | 0.0 | C32 H43 N5 O6 F2 Na |

**Elemental Composition Report** Page 1

### HRMS of ART1

**Single Mass Analysis**
Tolerance = 5.0 PPM / DBE: min = -1.5, max = 50.0
Element prediction: Off
Number of isotope peaks used for i-FIT = 3

Monoisotopic Mass, Even Electron Ions
32 formula(e) evaluated with 1 results within limits (up to 50 closest results for each mass)
Elements Used:
C: 0-17 H: 0-45 N: 0-1 O: 0-6 Na: 0-1

Rong
Rong
C17H27NO5
WZ_18_40 25 (0.570) Cm (25:31)

NMR Analytical Core Facility
LCT Premier XE

28-Sep-2018
2::4::2

1: TOF MS ES+
8.79e+004

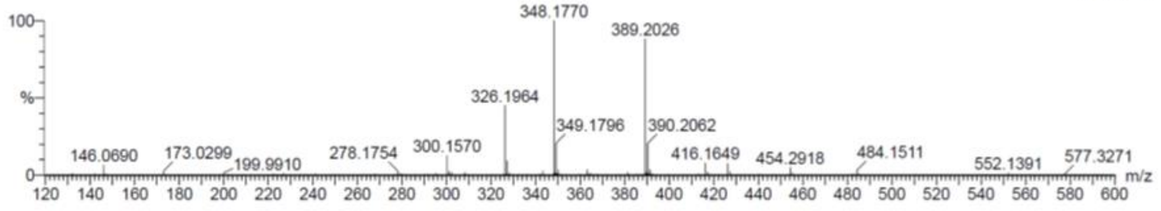

| | | | |
|---|---|---|---|
| Minimum: | | | -1.5 |
| Maximum: | 5.0 | 5.0 | 50.0 |

| Mass | Calc. Mass | mDa | PPM | DBE | i-FIT | i-FIT (Norm) | Formula |
|---|---|---|---|---|---|---|---|
| 348.1770 | 348.1787 | -1.7 | -4.9 | 4.5 | 545.8 | 0.0 | C17 H27 N O5 Na |

**Elemental Composition Report**

### HRMS of ATZ1

**Single Mass Analysis**
Tolerance = 5.0 PPM / DBE: min = -1.5, max = 50.0
Element prediction: Off
Number of isotope peaks used for i-FIT = 3

Monoisotopic Mass, Even Electron Ions
326 formula(e) evaluated with 1 results within limits (up to 50 closest results for each mass)
Elements Used:
C: 0-40 H: 0-56 N: 0-5 O: 0-9 F: 0-2 Na: 0-1

Rong
Rong
C40H52F2N4O8
WZ_Hybrid1_755 41 (0.932) Cm (41:46)

NMR Analytical Core Facility
LCT Premier XE

28-Sep-2018
2::4::0

1: TOF MS ES+
1.48e+005

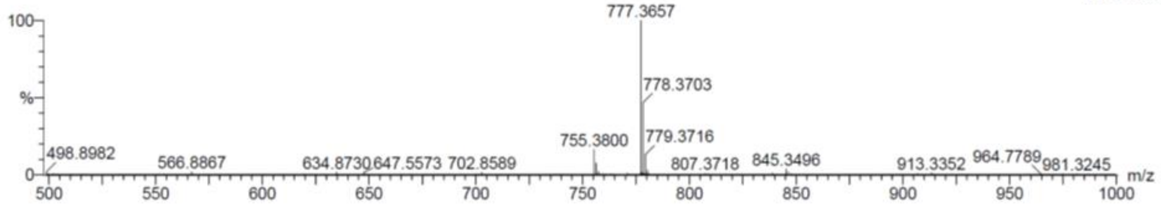

| | | | | |
|---|---|---|---|---|
| Minimum: | | | | -1.5 |
| Maximum: | | 5.0 | 5.0 | 50.0 |

| Mass | Calc. Mass | mDa | PPM | DBE | i-FIT | i-FIT (Norm) | Formula |
|---|---|---|---|---|---|---|---|
| 777.3657 | 777.3651 | 0.6 | 0.8 | 15.5 | 331.8 | 0.0 | C40 H52 N4 O8 F2 Na |

**Elemental Composition Report**

### HRMS of ATZ2

**Single Mass Analysis**
Tolerance = 5.0 PPM / DBE: min = -1.5, max = 50.0
Element prediction: Off
Number of isotope peaks used for i-FIT = 3

Monoisotopic Mass, Even Electron Ions
347 formula(e) evaluated with 1 results within limits (up to 50 closest results for each mass)
Elements Used:
C: 0-42 H: 0-56 N: 0-5 O: 0-9 F: 0-2 Na: 0-1

Rong
Rong
C42H55F2N5O9
WZ_Hybrid2_811 16 (0.350) Cm (11:16)

NMR Analytical Core Facility
LCT Premier XE

28-Sep-2018
2::8::0

1: TOF MS ES+
1.49e+005

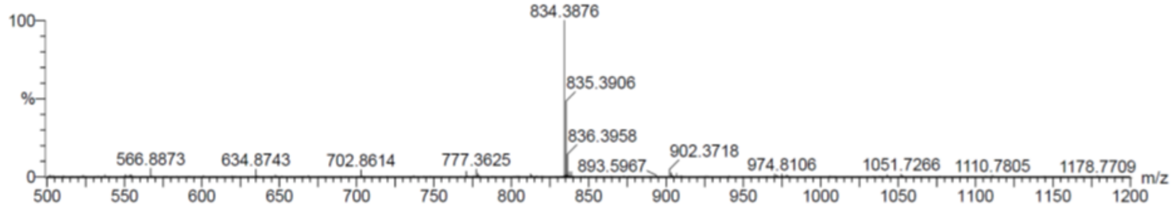

| | | | | |
|---|---|---|---|---|
| Minimum: | | | | -1.5 |
| Maximum: | | 5.0 | 5.0 | 50.0 |

| Mass | Calc. Mass | mDa | PPM | DBE | i-FIT | i-FIT (Norm) | Formula |
|---|---|---|---|---|---|---|---|
| 834.3876 | 834.3866 | 1.0 | 1.2 | 16.5 | 370.2 | 0.0 | C42 H55 N5 O9 F2 Na |

**Elemental Composition Report**

### HRMS of ATZ3

Page 1

**Single Mass Analysis**
Tolerance = 5.0 PPM / DBE: min = -1.5, max = 50.0
Element prediction: Off
Number of isotope peaks used for i-FIT = 3

Monoisotopic Mass, Even Electron Ions
347 formula(e) evaluated with 1 results within limits (up to 50 closest results for each mass)
Elements Used:
C: 0-43 H: 0-57 N: 0-5 O: 0-9 F: 0-2 Na: 0-1

Rong
Rong
C43H57F2N5O9
WZ_Hybrid3_825 29 (0.661) Cm (26:32)

NMR Analytical Core Facility
LCT Premier XE

28-Sep-2018
2::5::0

1: TOF MS ES+
4.55e+004

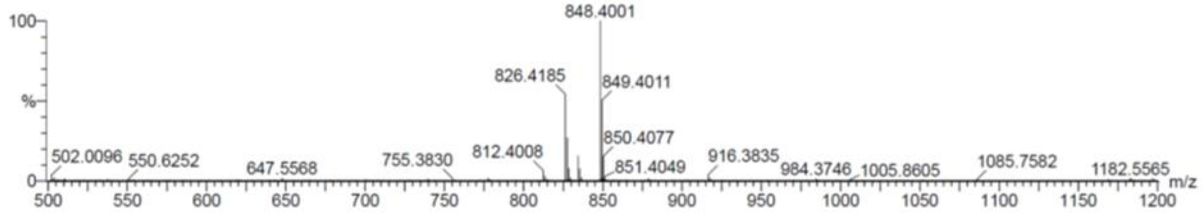

| | | | | | | | |
|---|---|---|---|---|---|---|---|
| Minimum: | | | | -1.5 | | | |
| Maximum: | | 5.0 | 5.0 | 50.0 | | | |
| Mass | Calc. Mass | mDa | PPM | DBE | i-FIT | i-FIT (Norm) | Formula |
| 848.4001 | 848.4022 | -2.1 | -2.5 | 16.5 | 304.9 | 0.0 | C43 H57 N5 O9 F2 Na |

**Elemental Composition Report**

### HRMS of ATZ4

Page 1

**Single Mass Analysis**
Tolerance = 5.0 PPM / DBE: min = -1.5, max = 50.0
Element prediction: Off
Number of isotope peaks used for i-FIT = 3

Monoisotopic Mass, Even Electron Ions
347 formula(e) evaluated with 1 results within limits (up to 50 closest results for each mass)
Elements Used:
C: 0-44 H: 0-60 N: 0-5 O: 0-9 F: 0-2 Na: 0-1

Rong
Rong
C44H59F2N5O9
WZ_Hybrid4_839 28 (0.622) Cm (28:33)

NMR Analytical Core Facility
LCT Premier XE

28-Sep-2018
2::8::8

1: TOF MS ES+
9.78e+004

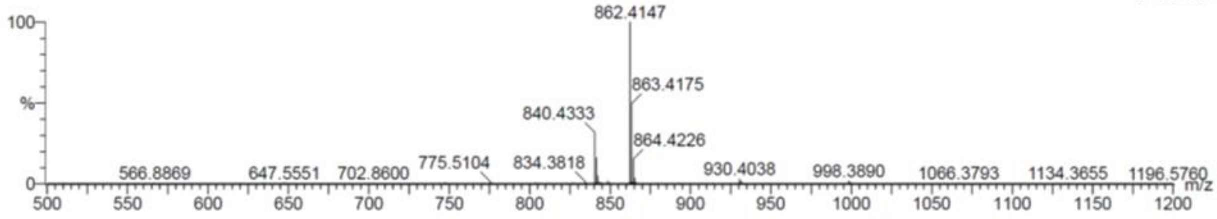

| | | | | | | | |
|---|---|---|---|---|---|---|---|
| Minimum: | | | | -1.5 | | | |
| Maximum: | | 5.0 | 5.0 | 50.0 | | | |
| Mass | Calc. Mass | mDa | PPM | DBE | i-FIT | i-FIT (Norm) | Formula |
| 862.4147 | 862.4179 | -3.2 | -3.7 | 16.5 | 327.2 | 0.0 | C44 H59 N5 O9 F2 Na |

#### Single Mass Analysis

### HRMS of WZ-20

Tolerance = 5.0 PPM / DBE: min = -1.5, max = 50.0
Element prediction: Off
Number of isotope peaks used for i-FIT = 2

Monoisotopic Mass, Even Electron Ions
112 formula(e) evaluated with 1 results within limits (up to 50 closest results for each mass)
Elements Used:
C: 0-27 H: 0-33 N: 0-5 O: 0-6 F: 0-2

Gang
Wenfu
C27H32F2N4O6
WZ_20_546 31 (0.696) Cm (31:34)

NMR Analytical Core Facility
LCT Premier XE

28-Mar-2019
1::7::3

1: TOF MS ES+
2.03e+004

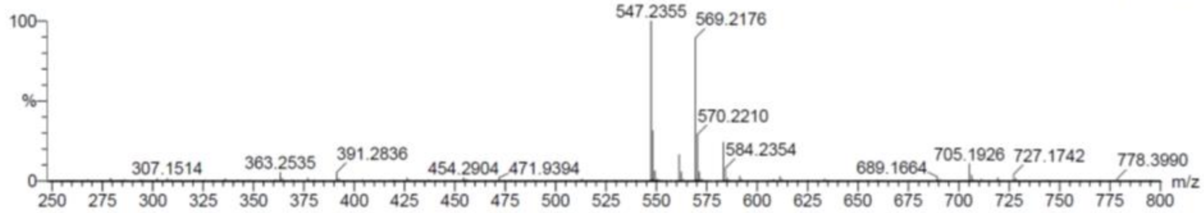

| | | | | | | | |
|---|---|---|---|---|---|---|---|
| Minimum: | | | | -1.5 | | | |
| Maximum: | | 5.0 | 5.0 | 50.0 | | | |
| Mass | Calc. Mass | mDa | PPM | DBE | i-FIT | i-FIT (Norm) | Formula |
| 547.2355 | 547.2368 | -1.3 | -2.4 | 12.5 | 132.6 | 0.0 | C27 H33 N4 O6 F2 |

#### Single Mass Analysis

### HRMS of WZ-06

Tolerance = 5.0 PPM / DBE: min = -1.5, max = 50.0
Element prediction: Off
Number of isotope peaks used for i-FIT = 2

Monoisotopic Mass, Even Electron Ions
112 formula(e) evaluated with 1 results within limits (up to 50 closest results for each mass)
Elements Used:
C: 0-27 H: 0-33 N: 0-5 O: 0-6 F: 0-2

Gang
Wenfu
C27H32F2N4O6
WZ_20_546 31 (0.696) Cm (31:34)

NMR Analytical Core Facility
LCT Premier XE

28-Mar-2019
1::7::3

1: TOF MS ES+
2.03e+004

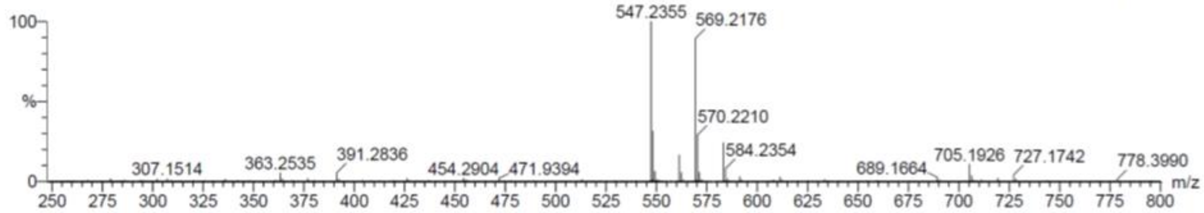

| | | | | | | | |
|---|---|---|---|---|---|---|---|
| Minimum: | | | | -1.5 | | | |
| Maximum: | | 5.0 | 5.0 | 50.0 | | | |
| Mass | Calc. Mass | mDa | PPM | DBE | i-FIT | i-FIT (Norm) | Formula |
| 547.2355 | 547.2368 | -1.3 | -2.4 | 12.5 | 132.6 | 0.0 | C27 H33 N4 O6 F2 |

**Single Mass Analysis**
Tolerance = 5.0 PPM / DBE: min = -1.5, max = 50.0
Element prediction: Off
Number of isotope peaks used for i-FIT = 2

### HRMS of WZ-13

Monoisotopic Mass, Even Electron Ions
182 formula(e) evaluated with 1 results within limits (up to 50 closest results for each mass)
Elements Used:
C: 0-42 H: 0-56 N: 0-5 O: 0-10 F: 0-2

Gang
Wenfu
C42H54F2N4O10
WZ_13_812 33 (0.752) Cm (31:36)

NMR Analytical Core Facility
LCT Premier XE

28-Mar-2019
1::3::4

1: TOF MS ES+
9.59e+004

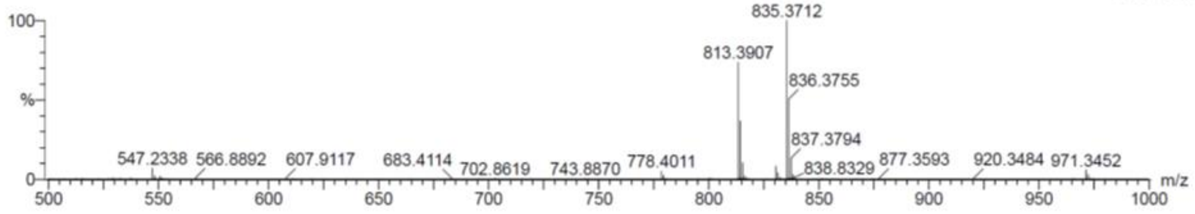

| | | | | | | | |
|---|---|---|---|---|---|---|---|
| Minimum: | | | | -1.5 | | | |
| Maximum: | | 5.0 | 5.0 | 50.0 | | | |
| Mass | Calc. Mass | mDa | PPM | DBE | i-FIT | i-FIT (Norm) | Formula |
| 813.3907 | 813.3886 | 2.1 | 2.6 | 16.5 | 179.0 | 0.0 | C42 H55 N4 O10 F2 |

**Elemental Composition Report**

### HRMS of Deoxy-ATZ4

Page 1

**Single Mass Analysis**
Tolerance = 5.0 PPM / DBE: min = -1.5, max = 50.0
Element prediction: Off
Number of isotope peaks used for i-FIT = 2

Monoisotopic Mass, Even Electron Ions
199 formula(e) evaluated with 1 results within limits (up to 50 closest results for each mass)
Elements Used:
C: 0-44 H: 0-60 N: 0-5 O: 0-8 F: 0-3

Gang
Gang
C44H59F2N5O8
deoxy_A7Z4 11 (0.244) Cm (9:11)

NMR Analytical Core Facility
LCT Premier XE

17-Feb-2021
3::8::2

1: TOF MS ES+
2.97e+004

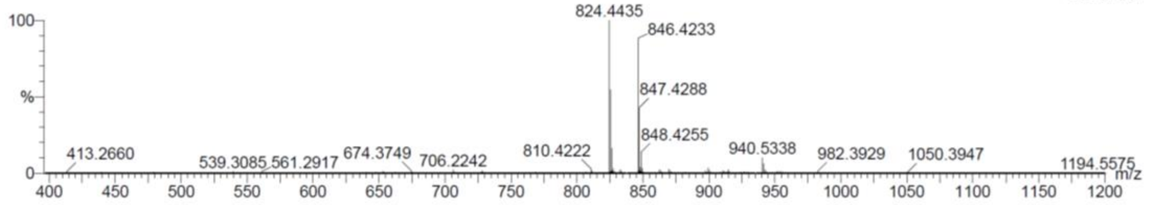

| | | | | | | | |
|---|---|---|---|---|---|---|---|
| Minimum: | | | | -1.5 | | | |
| Maximum: | | 5.0 | 5.0 | 50.0 | | | |
| Mass | Calc. Mass | mDa | PPM | DBE | i-FIT | i-FIT (Norm) | Formula |
| 824.4435 | 824.4410 | 2.5 | 3.0 | 16.5 | 138.2 | 0.0 | C44 H60 N5 O8 F2 |
